# Glycosylation of serine/threonine-rich intrinsically disordered regions of membrane-associated proteins in streptococci

**DOI:** 10.1101/2024.05.05.592596

**Authors:** Mohammad M. Rahman, Svetlana Zamakhaeva, Jeffrey S. Rush, Catherine T. Chaton, Cameron W. Kenner, Yin Mon Hla, Ho-Ching Tiffany Tsui, Vladimir N. Uversky, Malcolm E. Winkler, Konstantin V. Korotkov, Natalia Korotkova

## Abstract

Proteins harboring intrinsically disordered regions (IDRs) lacking stable secondary or tertiary structures are abundant across the three domains of life. These regions have not been systematically studied in prokaryotes. Our genome-wide analysis identifies extracytoplasmic serine/threonine-rich IDRs in several biologically important membrane-associated proteins in streptococci. We demonstrate that these IDRs are glycosylated with glucose by glycosyltransferases GtrB and PgtC2 in *Streptococcus pyogenes* and *Streptococcus pneumoniae*, and with N-acetylgalactosamine by a Pgf-dependent mechanism in *Streptococcus mutans*. The absence of glycosylation leads to a defect in biofilm formation under ethanol-stressed conditions in *S. mutans*. We link this phenotype to the C-terminal IDR of the post-translocation chaperone PrsA. Our data reveal that *O*-linked glycosylation protects the IDR-containing proteins from proteolytic degradation and is critical for the biological function of PrsA in biofilm formation.

## Introduction

Many protein regions, and even entire proteins, do not adopt a defined three-dimensional structure under physiological conditions. These protein regions are known as intrinsically disordered regions (IDRs) ^1, 2, 3^. IDRs are abundant in eukaryotes, where more than 30% of proteins are predicted to have extended IDRs with disordered segments longer than 30 residues ^4, 5^. These regions play a wide variety of essential roles and are associated with many human diseases, including cancer, cardiovascular disease, diabetes, and neurodegeneration^6^. A critical feature of many characterized IDRs is weak, transient, multivalent interactions with various binding partners, including proteins, DNA, RNA, and small molecules. These attractive intermolecular and intramolecular interactions between IDRs have been shown to contribute to the formation of condensed liquid-like droplets that play important biological roles in the compartmentalization of proteins and regulation of biochemical reactions ^7^. Conformations, dynamics, and interactions of IDRs are particularly sensitive to the crowded milieu of the cell ^8, 9, 10^. In spatially constricted regions of a cell membrane, IDRs of membrane proteins can promote oligomerization and clustering required for signal amplification in signal transduction pathways ^9, 10, 11^. Another important consequence of IDRs is a large increase in hydrodynamic radius and high conformational entropy ^12, 13^. IDRs of membrane proteins generate a substantial entropic force at the point of attachment ^14^ to drive membrane bending ^15^. Furthermore, some IDRs can prevent protein crowding by entropically excluding large neighboring molecules. This property of IDRs promotes stability and folding of adjacent domains and has been used to design fusion tags to enhance the soluble expression of recombinant proteins ^16, 17^. Because disordered domains are highly accessible for modifying enzymes, IDRs frequently serve as the sites for a broad spectrum of post-translational modifications ^18^. In eukaryotes, *O*-linked glycosylation has been predicted to be preferentially located within IDRs of extracellular proteins ^19^ and implicated in the protection of these regions from proteolysis ^20^ and regulation of protein aggregation and stability ^21, 22^. However, understanding the impact of glycosylation on the biological function of IDRs and its role in most cellular processes remains to be elucidated.

In contrast, few proteins bearing IDRs have been reported in prokaryotes ^23^. Protein disorder in prokaryotes has not been systematically identified and characterized even in model organisms such as *Escherichia coli* and *Bacillus subtilis*. However, recent experimental evidence highlights the importance of prokaryotic IDRs in various aspects of cell physiology, including signaling pathways, protein sorting mechanisms, cell division, and liquid-liquid phase separation ^24, 25, 26, 27, 28, 29, 30, 31^. Bacterial proteins that function in the external milieu play a crucial role in growth and pathogenesis by maintaining cell envelope integrity, providing protection from various environmental stimuli, facilitating the uptake and secretion of different products, and sensing external signals. In Gram-positive bacteria, these processes occur in the highly crowded space between the plasma membrane and the cell wall peptidoglycan layer, which is also exposed to fluctuations in the external environment. Extended disordered regions have been reported in cell wall-anchored adhesins and secreted enzymes in these organisms ^32^. However, little is known about the function of extracytoplasmic IDRs in membrane-associated proteins. Our study addresses this important aspect of the proteome in three Gram-positive bacteria: *Streptococcus mutans*, *Streptococcus pyogenes*, and *Streptococcus pneumoniae*. We reveal that these important human pathogens possess a unique subset of extracytoplasmic IDRs in membrane and secreted proteins involved in critical cellular processes such as protein folding, cell wall biosynthesis, cell division, and signal transduction. We demonstrate that the *S. pyogenes* and *S pneumoniae* IDRs are decorated extracellularly with glucose (Glc) by a previously undescribed mechanism, and the *S. mutans* IDRs are glycosylated with GalNAc by Pgf-dependent pathway enzymes. Previously, *O*-glycosylated proteins have been reported to be present on the surface of different Gram-positive bacteria ^33^. The identified glycoproteins are cell wall-anchored adhesins with serine-rich repeat regions. Decoration of these adhesins with sugars has been shown to take place in the cytoplasm, followed by their export across the plasma membrane by the accessory Sec system ^33^. In contrast, *O*-glycosylation of the *S. mutans* cell wall-anchored adhesins Cnm and WapA with *O*-N-acetylhexosamines, presumably N-acetylgalactosamine (GalNAc) and N-acetylglucosamine (GlcNAc), was reported to require the Pgf gene cluster encoding the GT-A type glycosyltransferase PgfS, an epimerase PgfE, and two GT-C type glycosyltransferases, PgfM1 and PgfM2 ^34, 35, 36^. The Pgf mechanism is unusual because the PgfM1 conserved catalytic residues are located on the outer face of the membrane, strongly supporting the notion that *O*-glycosylation occurs in extracellular space.

*S. mutans* is the major contributor to human dental plaque formation, primarily living in biofilms on the tooth surfaces ^37^. We discover that *O*-glycosylation of the C-terminal IDR of PrsA by the Pgf-dependent mechanism is required for protein-based biofilm formation of *S. mutans* under ethanol stress. PrsA homologs in Gram-positive bacteria are lipoproteins that play crucial roles in the export and maturation of membrane and secreted proteins, including numerous virulence factors ^38, 39, 40, 41, 42, 43, 44, 45, 46, 47^. We further demonstrate that glycosylation of some IDRs protects the glycoproteins from proteolytic cleavage and impacts their protein level. Hence, our report not only reveals the mechanism utilized by *S. mutans* to modulate biofilm synthesis but also provides a roadmap for future functional studies of glycosylated IDRs in prokaryotes.

## Results

### GtrB and PgtC2 catalyze glucosylation of *S. pyogenes* membrane-associated proteins

In bacteria, GT-A type glycosyltransferases typically transfer a sugar moiety from nucleotide-sugars to a lipid carrier, undecaprenyl phosphate (Und-P), which subsequently serves as sugar donor to a wide range of acceptor molecules, including proteins and polysaccharides ^48, 49^. We recently demonstrated that in *S. mutans,* a GT-A type glycosyltransferase SccN is a UDP-glucose:Und-P glucosyl transferase, which synthesizes β-Glc-P-Und ^50^. This glycolipid is required for modification of the cell wall serotype *c* rhamnopolysaccharide, SCC, with α-glucose (Glc) side-chains ^51^. Many Gram-positive bacteria, including streptococcal species such as *S. pyogenes* and *S. pneumoniae*, encode an SccN homolog, GtrB ^48^, of previously unknown function (Supplementary Fig. 1). To investigate the enzymatic activity of this enzyme in *S. pyogenes* (M5005_Spy_420, hereafter GtrB^Spy^), we expressed the protein on a plasmid in the *E. coli* JW2347 strain which is devoid of glucosyl phosphoryl undecaprenol (Glc-P-Und) synthase activity ^52^. When GtrB^Spy^ activity was assayed in *E. coli* membrane fractions *in vitro* using UDP-[^3^H]Glc and exogenously added Und-P ^49^, we observed robust incorporation of [^3^H]Glc into the glycolipid fraction. No incorporation of radioactivity into [^3^H]glycolipid was noted when the reaction was conducted with the *E. coli* strain carrying an empty plasmid (Supplementary Table 1). A kinetic analysis demonstrated that GtrB^Spy^ possesses a high affinity for UDP-Glc and Und-P (8 μM and 51 μM, respectively), suggesting that GtrB^Spy^ is an active Glc-P-Und synthase (Supplementary Fig. 2a, b). Analysis of the products by TLC on silica gel detected a single peak of radioactivity, with chromatographic mobility expected for [^3^H] Glc-P-Und (Supplementary Fig. 2c). To investigate GtrB^Spy^ activity in the *S. pyogenes* membranes, we constructed a GtrB-deficient mutant in the *S. pyogenes* MGAS2221 background (2221Δ*gtrB*) and complemented 2221Δ*gtrB* with a plasmid expressing the WT copy of *gtrB^Spy^* (2221Δ*gtrB*:p*gtrB*). When the membranes of these strains were incubated with UDP-[^3^H]Glc and Und-P, two classes of glucolipids were detected in wild-type (WT) and 2221Δ*gtrB*:p*gtrB*: a major glucolipid product with properties consistent with a glucosyldiglyceride and a minor glucolipid, presumably Glc-P-Und (Fig. 1a, c), which co-migrates on TLC with the lipid accumulating in the *E. coli* pBAD33_GtrB^Spy^ membranes (Supplementary Fig. 2c). Only the major glucolipid product was detected in 2221Δ*gtrB* (Fig. 1b). When glucosyldiglyceride was eliminated by mild alkaline de-acylation, the WT membranes contained a single peak of radioactivity corresponding to Glc-P-Und (Supplementary Fig. 3a). The membranes of 2221Δ*gtrB* did not catalyze the synthesis of Glc-P-Und (Supplementary Fig. 3b). The complementation of the mutant with the plasmid resulted in a robust increase in Glc-P-Und synthase activity (Supplementary Fig. 3c). The minor lipid product was stable to mild alkaline methanolysis, but sensitive to mild acid treatment (Supplementary Table 2), properties consistent with its identification as Glc-P-Und ^49^. Thus, these results implicate GtrB^Spy^ in the synthesis of Glc-P-Und.

**Fig. 1.**
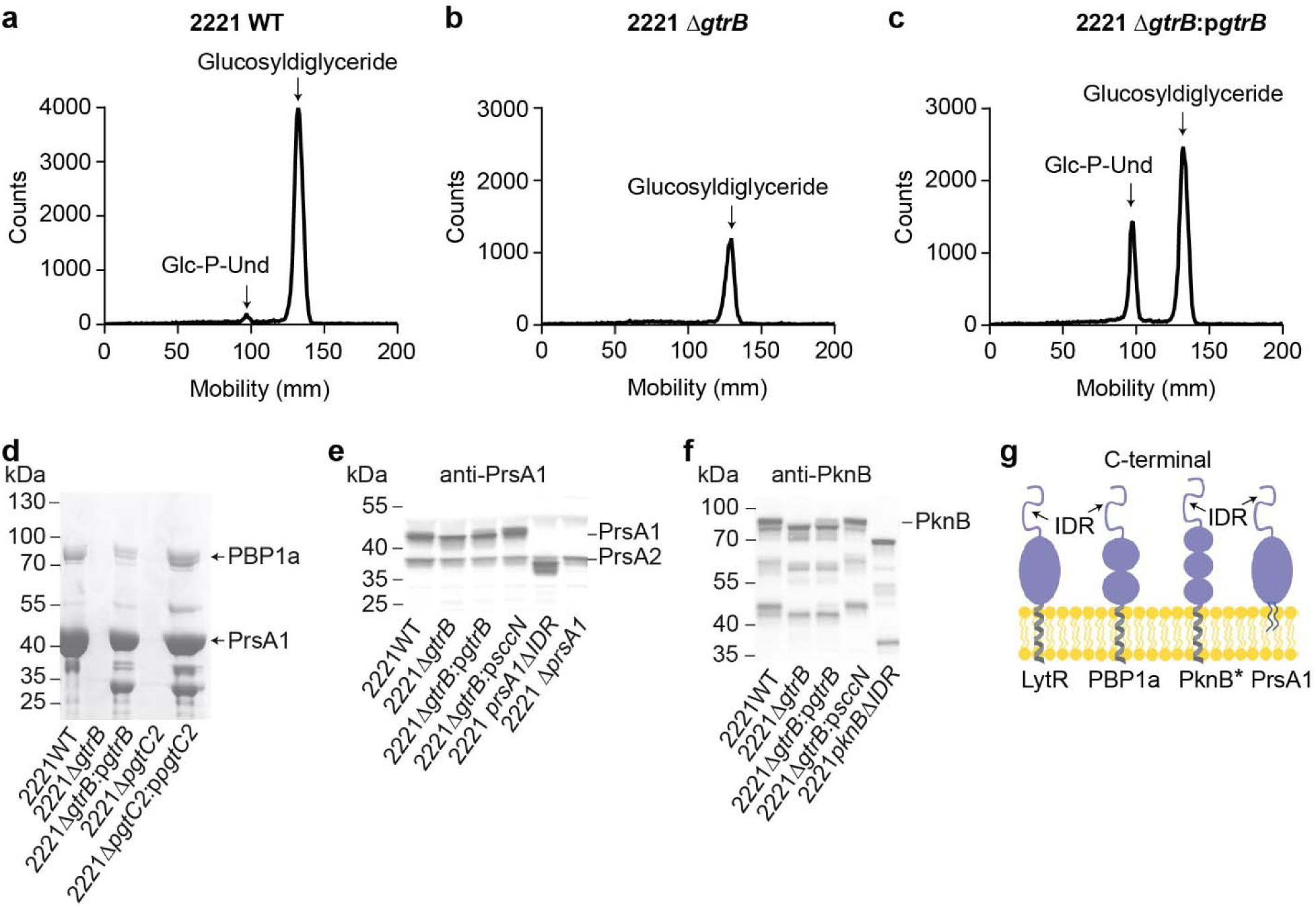
Glucosylation of *S. pyogenes* membrane-associated proteins. **a**, **b**, **c**, Representative TLC analysis of [^3^H]Glc-lipids extracted from *in vitro* incubations of UDP-[^3^H]Glc with the membrane fractions isolated from *S pyogenes* WT (**a**), Δ*gtrB* (**b**), and Δ*gtrB*:p*gtrB* (**c**). **d**, Coomassie-stained gel of *S. pyogenes* membrane proteins purified by conA affinity chromatography. Proteins in excised bands were identified by LC-MS/MS analysis as described in Methods. The major identified proteins are indicated. **e**, **f**, Immunoblot analysis of PrsA1 (**e**) and PknB (**f**) in *S. pyogenes* WT, Δ*gtrB*, Δ*gtrB*:p*gtrB*, and Δ*gtrB*:p*sccN* using specific antibodies. Anti-PrsA1 antibodies also recognize PrsA2. Proteins were separated on a 4-12% SurePAGE™ Bis-Tris gel (Genscript) in MES buffer in **d**, **e**, **f**. The experiments were performed independently three times in **a**, **b**, **c**, **e**, and **f**, and two times in **d**, yielding the same results. A representative image from one experiment is shown. **g,** Topology of extracytoplasmic domains of *S. pyogenes* conA-bound proteins. * The cytoplasmic domain of PknB is omitted for clarity. Just one monomer of the dimer for PrsA1 and PknB is shown for clarity. PrsA1 is depicted as a diacylated lipoprotein because lipoproteins in streptococci are diacylated as N-acyl-glyceryl-cysteine ^96, 97^. Source data for **a**, **b**, **c**, **d**, **e** and **f** are provided as a Source Data file.

Since *S. pyogenes* cell wall polysaccharide does not contain Glc moieties ^53^, we hypothesized that GtrB^Spy^ is involved in the first step of protein glucosylation. To test the role of GtrB^Spy^ in glucosylation of membrane proteins, we employed lectin affinity chromatography on sepharose-bound concanavalin A (ConA) to purify glycoproteins from detergent-solubilized extracts of the WT, 2221Δ*gtrB*, and 2221Δ*gtrB*:p*gtrB* membranes. ConA lectin selectively recognizes α-mannose/α-glucose residues ^54, 55^. Bound putative glycoproteins were eluted by the hapten sugar, methyl α-D-mannopyranoside. SDS-PAGE analysis demonstrated that ConA-bound glycoproteins are present in WT and 2221Δ*gtrB*:p*gtrB*, but not in 2221Δ*gtrB* (Fig. 1d). This result establishes that *S. pyogenes* membrane proteins undergo glucosylation that requires the participation of GtrB^Spy^. The fact that glycoproteins are bound tightly by ConA confirms the presence of non-reducing terminal α-Glc units.

We hypothesized that the final step of protein glucosylation is catalyzed by an unknown GT-C fold glycosyltransferase, which transfers α-Glc from Glc-P-Und to membrane proteins. A similar enzyme family participates in asparagine-linked *N-* glycosylation and serine or threonine-linked *O-*glycosylation in higher-order eukaryotes and some Gram-negative bacteria ^56^. *S. pyogenes* encodes two GT-C type glycosyltransferases, M5005_Spy_1794 and M5005_Spy_1862 (hereafter PgtC1 and PgtC2^Spy^), with unknown functions. PgtC2^Spy^ belongs to the YfhO protein family, which is universally present in Firmicutes (Supplementary Fig. 4). Interestingly, in *B. subtilis* and *Listeria monocytogenes*, YfhO homologs are involved in the transfer of sugar moieties to teichoic acid polymers ^57^. We constructed the PgtC1 and PgtC2^Spy^ deletion mutants (2221Δ*pgtC1* and 2221Δ*pgtC2*) and the corresponding complemented strains (2221Δ*pgtC1*:p*pgtC1* and 2221Δ*pgtC2*:p*pgtC2*) in *S. pyogenes*. We used these strains for the purification of glycoproteins on ConA as described above. This approach revealed that glucosylation of membrane proteins requires the participation of PgtC2^Spy^, but PgtC1 likely has no or a minor role in this mechanism (Fig. 1d and Supplementary Fig. 5).

### *S. pyogenes* PrsA1, PBP1a, LytR and PknB are glycoproteins

Mass spectrometry (MS)-based proteomic analysis of *S. pyogenes* glycoproteins present in the gel bands (Fig. 1d, lane 1), described above, identified a conserved post-translational chaperone lipoprotein of peptidyl-prolyl cis/trans isomerase family PrsA1 ^39^ and a class A penicillin-binding protein PBP1a as the most abundant proteins in the WT eluate. Additionally, the analysis detected LytR, which belongs to LytR-Cps2A-Psr (LCP) family proteins involved in the attachment of polysaccharides to peptidoglycan ^58^, serine/threonine (Ser/Thr) protein kinase PknB ^59, 60^ and a homolog of PrsA1, PrsA2. Mass spectrometry identification of proteins purified by lectin affinity chromatography is provided in Supplementary Data 1. To validate the results of lectin affinity enrichment, we evaluated the expression of PrsA1, PrsA2, and PknB in WT, 2221Δ*gtrB*, and 2221Δ*gtrB*:p*gtrB* using protein-specific polyclonal antibodies. Deletion of *gtrB^Spy^* resulted in increased electrophoretic mobility of PrsA1 and PknB, but not PrsA2 (Fig. 1e, f and Supplementary Fig. 6a). Because PrsA1 and PrsA2 have 65% amino acid identity (Supplementary Fig. 6b), it is possible that PrsA2 was either misidentified by proteomics or co-purified in a complex with PrsA1. The electrophoretic behavior of PrsA1 and PknB in 2221Δ*gtrB* is consistent with an alteration of molecular weight due to a loss of Glc moieties. Interestingly, complementation of 2221Δ*gtrB* with the *S. mutans sccN* on a plasmid more efficiently restored PrsA1 and PknB electrophoretic behavior than the plasmid-expressed *gtrB*^Spy^ (Fig. 1e, f). These experiments support the conclusion that the observed change in the molecular weight of PrsA1 and PknB in 2221Δ*gtrB* is due to the absence of α-Glc, donated by β-Glc-P-Und.

### Serine/Threonine (Ser/Thr)-rich IDR of *S. pyogenes* PrsA1 is glycosylated

Multiple studies reported that ConA binds weakly to α-mannose/α-glucose monosaccharides, whereas the lectin exhibits a high affinity to oligosaccharides ^54, 55^. The strength of the association of ConA with *S. pyogenes* glycoproteins implies the presence of clusters of α-Glc units that provide a tight binding site for ConA. When we analyzed the AlphaFold2 ^61^ structural models of PrsA1, PBP1a, PknB, and LytR, we observed that their C-terminal regions lack secondary structure and have low pLDDT (predicted local distance difference test) confidence scores (Fig. 1g and Supplementary Fig. 7). The low pLDDT scores correlate with predicted disorder and can serve as a predictor of intrinsic disorder ^62, 63^. Visual inspection of the amino acid sequences of C-terminal regions revealed the presence of serine (Ser) and threonine (Thr) rich regions, suggesting *O*-glucosylation of these sites (Fig. 2a). To examine whether the PrsA1 IDR carries Glc residues, we constructed *S. pyogenes* mutant, 2221*prsA1*ΔIDR, lacking this region. Glycoprotein enrichment on ConA using the solubilized membranes of 2221*prsA1*ΔIDR determined that the IDR-less PrsA1 does not bind the lectin (Fig. 2b). These results suggest that the GtrB/PgtC2 mechanism specifically targets extracytoplasmic Ser/Thr IDRs for glucosylation. Interestingly, contrary to PrsA1, PrsA2 has a 9-residue C-terminal IDR, which lacks Ser and Thr residues, the potential sites for *O*-linked glycosylation (Supplementary Fig. 6b, c). This is in agreement with the observation that glycosylation does not impact the electrophoretic mobility of PrsA2.

**Fig. 2.**
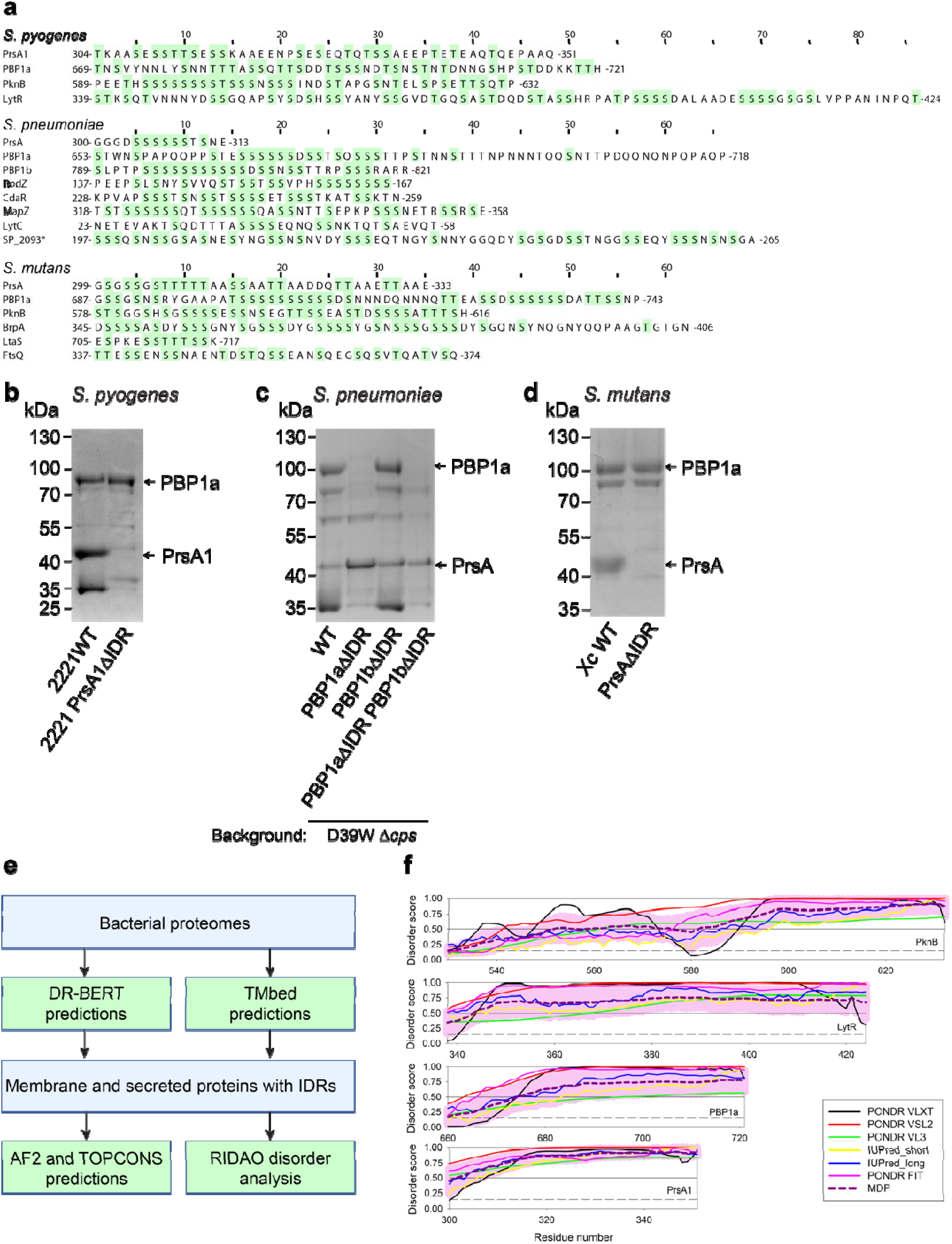
Features of Ser/Thr-rich IDRs identified in streptococcal extracytoplasmic proteins. **a**, IDR sequences of extracytoplasmic proteins. Ser and Thr residues are highlighted in green. Numbers correspond to amino acid residue starting and ending the IDR sequence. A * symbol denotes a TIGR4-specific protein. **b**, A Coomassie-stained gel of *S. pyogenes* membrane proteins purified by ConA affinity chromatography. **c**, A Coomassie-stained gel of *S. pneumoniae* D39W membrane proteins purified by ConA affinity chromatography. **d**, A Coomassie-stained gel of *S. mutans* membrane proteins purified on the WFA-conjugated agarose. The proteins were eluted with 300 mM methyl α-D-mannopyranoside in **b** and **c**, and with 50 mM GalNAc in **d**. Proteins were separated on a 4-12% SurePAGE™ Bis-Tris gel (Genscript) in MES buffer. The experiments were performed independently two times in **b**, and **d**, and three times in **c**. A representative image from one experiment is shown. **e**, A flowchart outlining a genome-wide analysis of IDRs in streptococci. **f**, Evaluation of the intrinsic disorder status of the Ser/Thr regions in *S. pyogenes* proteins analyzed in this study. Plots represent the per-residue disorder profiles generated for the indicated proteins by the Rapid Intrinsic Disorder Analysis Online (RIDAO) platform ^75^. RIDAO integrates six established disorder predictors, such as PONDR® VLS2, PONDR® VL3, PONDR® VLXT, PONDR® FIT, IUPred-Long, and IUPred-Short into a single, unified platform. The mean disorder profiles (MDP) for these proteins were calculated for each protein by averaging the disorder profiles of individual predictors. The light pink shade represents the MDP error distribution. The thin black line at the disorder score of 0.5 is the threshold between order and disorder, where residues/regions above 0.5 are disordered, and residues/regions below 0.5 are ordered. The dashed line at the disorder score of 0.15 is the threshold between order and flexibility, where residues/regions above 0.15 are flexible, and residues/regions below 0.15 are mostly ordered. Analysis of *S. pneumoniae* and *S. mutans* proteins is shown in Supplementary Fig. 10. Source data for **b, c** and **d** are provided as a Source Data file.

### Membrane-associated proteins harboring Ser/Thr-rich IDRs are widespread in streptococci and other Firmicutes

Previous studies have reported that the glycoproteins in Firmicute organisms are species-specific, frequently involved in adherence, and modified by species-specific protein glycosylation mechanisms ^33, 35^. In light of the observation that *gtrB* and *pgtC2* are widespread in streptococcal species, we suggested that glycosylation of IDRs is a common feature of extracellular membrane-associated proteins in these bacteria. We considered that the ConA-enrichment strategy may fail to capture all *S. pyogenes* glycoproteins. Therefore, to uncover a comprehensive picture of the streptococcal glycoproteome, we developed an automated pipeline that uses protein language-based models TMbed and DR-BERT to annotate membrane and secreted proteins, and disorder propensities for all proteins in the bacterial proteome (Fig. 2e, Methods) ^64, 65^. We focused the analysis on the proteomes of three streptococcal species, *S. pyogenes*, *S. pneumoniae*, and *S. mutans*, that are phylogenetically distinct belonging to three separate species groups — “Pyogenic”, “Mitis”, and “Mutans” ^66^ with the goal of understanding whether these bacteria possess the same set of membrane-associated proteins harboring Ser/Thr-rich IDRs.

In the proteomes of *S. pyogenes* MGAS2221, *S. mutans* Xc, and *S. pneumoniae* TIGR4, we identified membrane and secreted proteins with predicted IDRs containing at least 10 consecutive residues (Supplementary Data 2, 3, and 4). TOPCONS ^67^ and AlphaFold2 ^61^ were employed to assign the relative membrane topology of selected membrane proteins harboring predicted IDRs. Additionally, we analyzed AlphaFold2 ^61^ models to validate the disorder predictions. Remarkably, we found that all analyzed streptococci possess proteins with Ser/Thr-rich IDRs (Fig. 2a and Supplementary Table 3). Among the proteins harboring this type of IDRs, we identified PrsA and PBP1a in all streptococci, PknB and LytR in *S. pyogenes* and *S. mutans* (LytR is known as BrpA ^68^), the extracytoplasmic regulator of c-di-AMP synthesis CdaR ^69^, cell division proteins MapZ ^70, 71, 72^ and RodZ ^73^ in *S. pneumoniae*, and lipoteichoic acid synthase LtaS and a divisome protein FtsQ ^74^ in *S. mutans* (Supplementary Figs. 7, 8 and 9, Supplementary Table 3). Additionally, these Ser/Thr-rich regions were shown to be highly disordered by several commonly used disorder predictors assembled into the RIDAO platform ^75^. In fact, in all the cases analyzed in this study, the Ser/Thr-rich regions are invariantly disordered and are often located within longer IDRs (Fig. 2f, Supplementary Fig. 10). This is an important but expected observation, since the amino acid sequences of many IDRs are known to be characterized by the presence of repeats, and since it was indicated that the level of repeat perfection correlates with their tendency to be intrinsically disordered or unstructured, with more perfect repeats being less structured ^76^. Interestingly, this type of IDR is frequently located at the C-termini of the membrane-associated proteins. Furthermore, protein sequence alignments revealed the presence of Ser/Thr-rich IDRs in the homologs of the identified proteins from streptococcal species and other Firmicutes (Supplementary Fig. 11, 12). To understand if IDRs carry important functional features, we generated sequence logos (WebLogo ^77^) for each subset of IDRs (Supplementary Fig. 13, 14). The results showed that the main feature of the PrsA, PBP1a, LtaS, PBP1b, PknB, and CdaR IDRs is the presence of poly-Ser/Thr tracks adjacent to a folded domain.

### GtrB and PgtC2 glucosylate Ser/Thr-rich IDRs in *S. pneumoniae*

To test a hypothesis that the GtrB/PgtC2 mechanism is involved in glucosylation of extracytoplasmic Ser/Thr-rich IDRs in *S. pneumoniae*, we employed the ConA enrichment strategy, described above, to purify the membrane glycoproteins from the *S. pneumoniae* serotype 2 D39W derivative strain (D39Δ*cps*) and the *S. pneumoniae* serotype 4 TIGR4 strain (Fig. 3a and Supplementary Fig. 15a). Proteomic analysis of captured proteins in excised bands revealed that both strains produce five putative glycoproteins: PrsA, PBP1a, MapZ, RodZ and CdaR. Additionally, we identified peptidoglycan hydrolase LytC ^78^ in D39W, and membrane protein SP_2093 in TIGR4. Furthermore, in both strains, in a protein band at around 100 kDa, the analysis identified predominantly the peptides corresponding to PBP1a and small amounts of peptides corresponding to PBP1b (Supplementary Data 1). As outlined above, C-terminal Ser/Thr-rich IDRs are present in PrsA, PBP1a, PBP1b, and CdaR. MapZ, RodZ, and SP_2093 possess linker IDRs. In contrast, LytC has an N-terminal IDR enriched with Ser and Thr (Fig. 2a, 3b and Supplementary Fig. 9). Significantly, contrary to *S. pyogenes* PknB, the *S. pneumoniae* PknB homolog, known as StkP, lacks the C-terminal IDR (Supplementary Fig. 15b) and does not bind to ConA sepharose. We constructed the GtrB (SPD_1431) deletion mutant, Δ*gtrB*, in the D39W derivative strain, and PgtC2 (SPD_2058) deletion mutant, Δ*pgtC2*, in the D39W and TIGR4 strain backgrounds to examine the roles of these enzymes in protein glucosylation. The glycoproteins were absent in the ConA-bound fractions of the mutants, indicating that the protein glucosylation pathway in *S. pneumoniae* is similar to *S. pyogenes* (Fig. 3a, Supplementary Fig. 15a). To examine how deletion of the C-terminal IDRs of PBP1a and PBP1b impacts binding of these proteins to ConA, we constructed single mutants, *pbp1a*ΔIDR and *pbp1b*ΔIDR, and the double mutant *pbp1a*ΔIDR*pbp1b*ΔIDR in the D39W derivative strain. The SDS-PAGE analysis of membrane proteins captured by ConA revealed that a 100 kDa protein band is absent in *pbp1a*ΔIDR and *pbp1a*ΔIDR*pbp1b*ΔIDR but present in WT and *pbp1b*ΔIDR (Fig. 2c). This result confirms PBP1a as the major glycoprotein in this band and demonstrates that the PBP1a IDR is critical for binding to ConA suggesting the presence of Glc residues on the IDR.

**Fig. 3.**
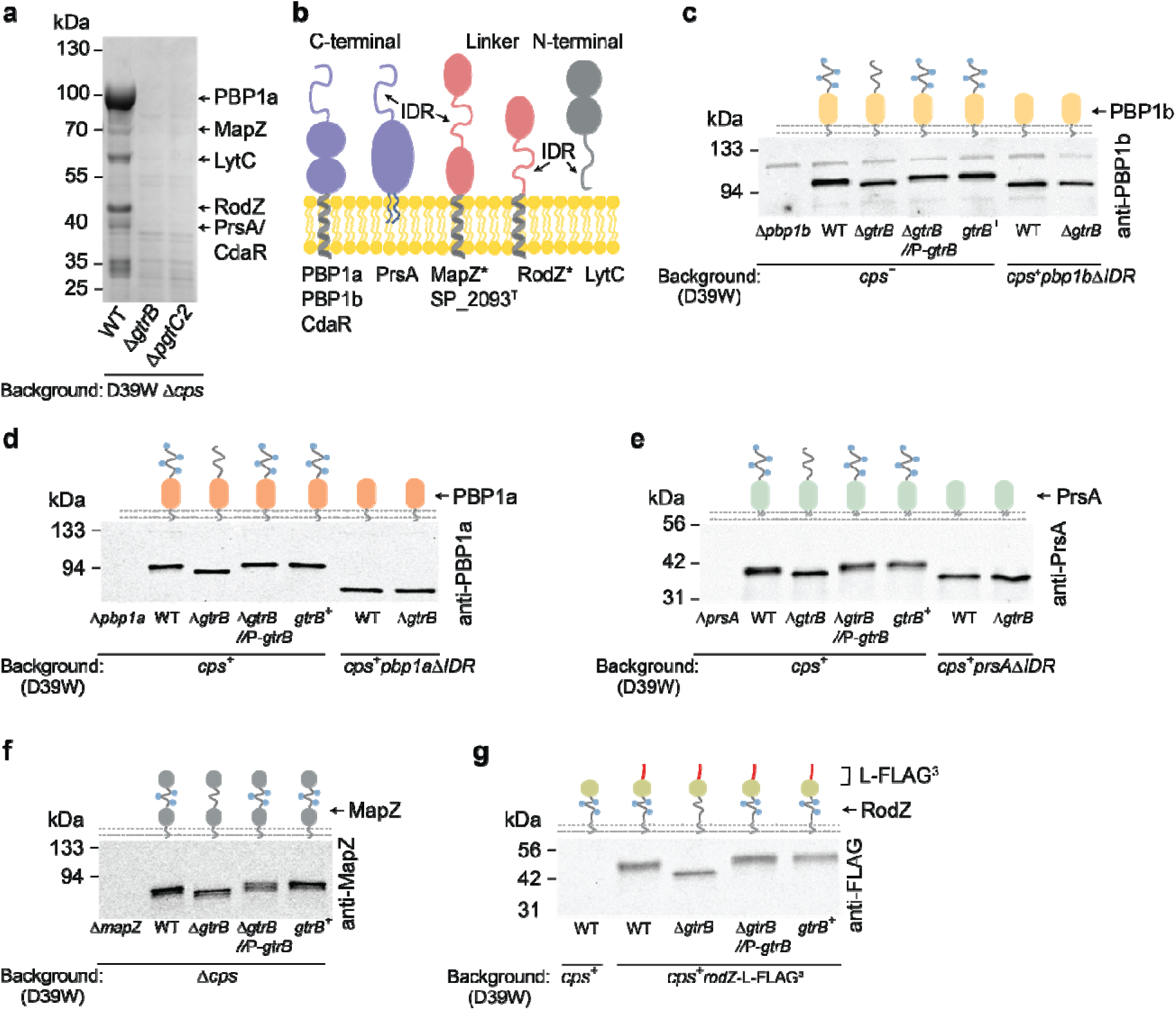
Glucosylation of *S. pneumoniae* extracytoplasmic proteins. **a**, Coomassie-stained gel of *S. pneumoniae* membrane proteins obtained from D39W Δ*cps* (IU1824), the isogenic Δ*gtrB* mutant (IU18454), and Δ*pgtC2* mutant (IU18456) purified by conA affinity chromatography. Proteins in excised bands were identified by LC-MS/MS analysis as described in Methods. The identified proteins are indicated. **b**, Topology of extracytoplasmic domains of *S. pneumoniae* conA-bound proteins. * The cytoplasmic domains of RodZ and MapZ are omitted for clarity. ^T^ denotes TIGR4 protein. Just one monomer of the dimer for PrsA is shown for clarity. PrsA is depicted as a diacylated lipoprotein. **c**, **d**, **e**, Immunoblot analysis of PBP1b (**c**), PBP1a (**d**), PrsA (**e**) in *cps*^+^ WT and the isogenic mutants. **f**, Immunoblot analysis of MapZ in Δ*cps* WT and the isogenic mutants. **g**, Immunoblot analysis of the RodZ-L-FLAG^3^ variant ^73^ in *cps* WT and the isogenic mutants. Samples obtained from *pbp1a*, *pbp1b, prsA*, *mapZ* deletion strains (left lanes) were used to demonstrate the specificity of the antibodies in **c**, **d**, **e**, **f**. The *cps*^+^ WT strain was used as a negative control in **g**. Proteins were separated on a 4-12% SurePAGE™ Bis-Tris gel (Genscript) in MES buffer in **a**, 7.5% Mini-PROTEAN® TGX™ precast protein gel (Biorad 4561025) in **c** and **d**, 10% precast protein gel (Biorad 4561035) in **e**, and 4–15% Mini-PROTEAN® TGX™ Precast Protein Gels (Biorad 4561085) in **f** and **g**. In **c**, **d**, **e**, **f** and **g** schematic representations of PBP1b, PBP1a, PrsA MapZ and RodZ are shown (top panel). Each of the single blob of PBP1b and PBP1a represents two extracellular domains. The cytoplasmic domains of RodZ and MapZ are omitted for clarity. Just one monomer of the dimer for PrsA is shown. Glc is depicted as blue circles. The experiments were performed independently two times in **a** and three times in **c**, **d**, **e, f, and g**. A representative image from one experiment is shown. Quantifications of protein loads and immunoblot band intensities of PBP1a, PBP1b, PrsA MapZ and RodZ are provided in Supplementary Fig. 15c, d, e, g and Supplementary Fig. 16. Source data for **a, c, d, e, f** and **g** are provided as a Source Data file.

To further confirm the results of the lectin-affinity enrichment, the electrophoretic behavior of PBP1b, PBP1a, PrsA, MapZ, RodZ, and StkP was assessed by immunoblot analysis in the D39W derivative strains (WT control), the corresponding isogenic Δ*gtrB* mutants and the mutants in which *gtrB^Spn^* was ectopically expressed from a constitutive promoter in the WT or Δ*gtrB* backgrounds. We detected that in the Δ*gtrB* mutant, PBP1b, PBP1a, PrsA, MapZ, and RodZ displayed an increase in the electrophoretic mobility, indicating a loss of sugar moieties (Fig. 3c, d, e, f, g). In contrast, no alteration in the electrophoretic behavior of the IDR-less PBP1b, PBP1a, PrsA, and MapZ variants was observed in the analyzed strain backgrounds, supporting the hypothesis that the IDRs are the sites of Glc attachment (Fig. 3c, d, e, f). As expected, the deletion of GtrB^Spn^ did not change the electrophoretic mobility of StkP (Supplementary Fig. 15i), providing evidence that this protein does not carry Glc residues. Quantitative analysis of glycoproteins by scanning of band staining established that glucosylation exerts a positive effect on the total protein levels of PBP1b, MapZ, and RodZ, but not on PrsA and PBP1a (Fig. 3c, e, and Supplementary Fig. 15c and 16a, c, d, e), and the presence of the IDRs are important for the PBP1a and MapZ protein levels (Fig. 3d, and Supplementary Fig. 15c and 16b, d).

### *S. mutans* Pgf pathway glycosylates Ser/Thr-rich IDRs with GalNAc

Surprisingly, we recovered no ConA-bound proteins from detergent-solubilized extracts of *S. mutans* Xc membranes despite the presence of GtrB (known as SccN) and PgtC2 (Smu_2160) homologs. However, according to our bioinformatics analysis, *S. mutans* Xc genome encodes the cell wall-anchored collagen-binding adhesins (CBAs) WapA, Cnm, CnaB, and CbpA harboring Ser/Thr-rich regions. It has been reported that Cnm is decorated by the Pgf-dependent mechanism with *O-*N-acetylhexosamines sugars, GalNAc and GlcNAc ^34, 35^. In *S. mutans* Xc, the Pgf gene cluster encodes the PgfS, PgfE, PgfM1 and PgfM2 enzymes and the Cnm, CnaB, and CbpA adhesins (Fig. 4a) ^35^. PgfM1 and PgfM2 are the structural homologs of PgtC2 being annotated as containing the YfhO domain (Supplementary Fig. 17a). We hypothesized that the Pgf pathway glycosylates extracytoplasmic Ser/Thr-rich IDRs of the membrane-associated proteins. To test this idea, we first focused on the identification of sugars attached to IDRs. We constructed the XcΔ*pgfS*, XcΔ*pgfM1*, XcΔ*pgfM2*, and XcΔ*pgtC2* mutants in *S. mutans* Xc and used these strains in a co-sedimentation assay with the fluorescent GalNAc and GlcNAc-specific lectins. We observed that the GalNAc-specific lectin, *Wisteria floribunda* agglutinin (WFA), binds to the WT, XcΔ*pgfM2*, and XcΔ*pgtC2* cells but not to the XcΔ*pgfS*, XcΔ*pgfM1* bacteria, and the mutant lacking the adhesins encoded by the Pgf gene cluster, XcΔ*cnaB-cbpA-cnm* (Supplementary Fig. 17b). Furthermore, *S. mutans* WT cells, but not the XcΔ*pgfS*, XcΔ*pgfM1* and *S. pyogenes* cells (used as a negative control) associate with other GalNAc-specific lectins, *Helix pomatia* agglutinin (HPA) and *Griffonia simplicifolia* lectin (BS-I) (Supplementary Fig. 17c, d). Binding of WFA to the WT cells was completely inhibited by the hapten sugar, GalNAc (Fig. 4b). In contrast to WFA, the GlcNAc-specific lectin, wheat-germ agglutinin (WGA), associated with the WT and XcΔ*pgfS* cells, and the addition of the WGA hapten sugar, GlcNAc, did not reduce the binding, indicating a non-specific interaction of WGA with *S. mutans* cells (Supplementary Fig. 17e). Fluorescent microscopy of *S. mutans* strains confirmed that WFA binds the cell surface of WT and XcΔ*pgfS* complemented with *pgfS* on a plasmid (XcΔ*pgfS*:p*pgfS*), but not XcΔ*pgfS* (Fig. 4c). Thus, these experiments strongly suggest that the enzymes of the Pgf pathway, PgfS and PgfM1, transfer GalNAc on the cell wall-anchored adhesins encoded by *cnaB-cbpA-cnm*.

**Fig. 4.**
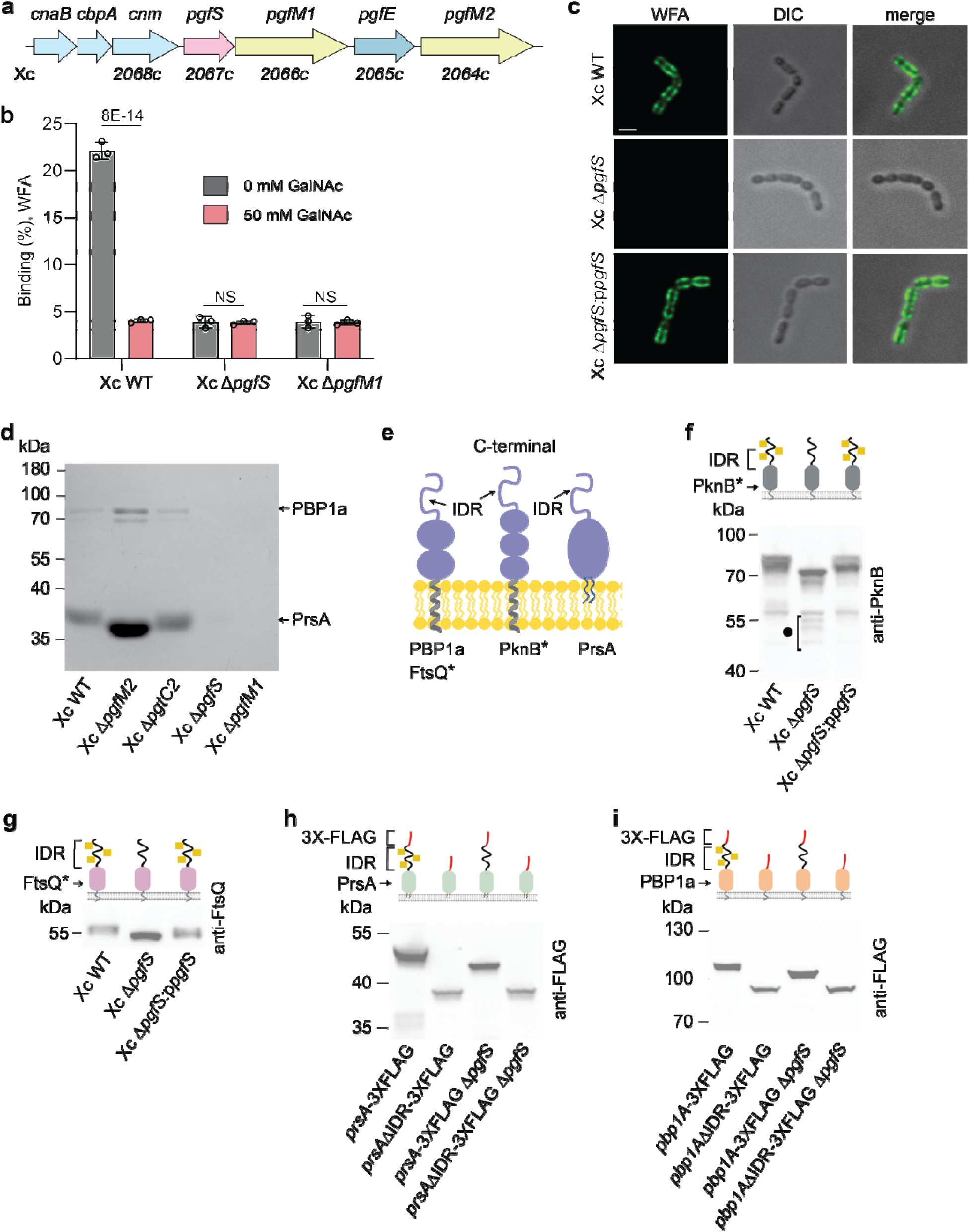
The *pgf* operon glycosylates *S. mutans* membrane-associated proteins with GalNAc. **a**, Schematic representation of the *pgf* operon in *S. mutans* Xc. **b**, Co-sedimentation of FITC-conjugated WFA with the *S. mutans* strains. 0 or 50 mM GalNAc was added to the binding buffer. Data were recorded as a total fluorescence *vs* a pellet fluorescence. Columns and error bars represent the mean and S.D., respectively (n = 3 biologically independent replicates). A two-way ANOVA with Bonferroni’s multiple comparisons test was performed to determine *P* values. **c**, Fluorescence and DIC images of bacteria bond to FITC-conjugated WFA. The scale bar is 1 µm. **d**, Coomassie-stained gel of *S. mutans* membrane proteins purified on the WFA-conjugated agarose column. Proteins in excised bands were identified by LC-MS/MS analysis as described in Methods. The identified proteins are indicated. **e**, Topology of extracytoplasmic domains of *S. mutans* WFA-bound proteins. * The cytoplasmic domains of PknB and FtsQ are omitted for clarity. Just one monomer of the dimer for PrsA and PknB is shown for clarity. PrsA is depicted as a diacylated lipoprotein. **f**, **g**, Immunoblot analysis of PknB (**f**) and FtsQ (**g**) in *S. mutans* cell lysates using anti-PknB and anti-FtsQ antibodies. In **f** and **g**, schematic representations of PknB and FtsQ are shown (top panel). Each of the single blobs of PknB and FtsQ represents three- and two extracellular domains, respectively. * The cytoplasmic domains are not shown in the schematic. **h**, **i**, Immunoblot analysis of 3×FLAG-tagged variants of PrsA (**h**) and PBP1a (**i**) in the WT and XcΔ*pgfS* backgrounds examined by anti-FLAG antibodies. Schematic of PrsA and PBP1a variants is shown in **h** and **i** top panels, respectively. Each of the single blobs of PBP1a represents two extracellular domains. GalNAc is depicted as yellow squares. Proteins were separated on a 15% SDS-PAGE gel in Tris-glycine buffer in **d** and a 4-12% SurePAGE™ Bis-Tris gel (Genscript) in MES buffer in **f**, **g**, **h**, and **i**. The experiments in **c**, **f**, **g**, **h,** and **i** were performed independently three times yielding the same results. A representative image from one experiment is shown. Quantifications of immunoblot band intensities of PrsA (**h**) and PBP1a (**i**) are provided in Supplementary Fig. 24. Source data for **b, d, f, g, h** and **i** are provided as a Source Data file.

Next, to identify *S. mutans* membrane proteins decorated with GalNAc, we employed WFA agarose for glycoprotein enrichment as we described for *S. pyogenes* and *S. pneumoniae*. Bound proteins were eluted with GalNAc, separated on the SDS-PAGE gel, and identified by MS analysis. The analysis of the WT eluate revealed the presence of peptides derived from PrsA and PBP1a in protein bands at around 40 and 80 kDa, respectively. We also detected small amounts of peptides derived from PknB, BrpA (LytR), and the cell division protein FtsQ (Supplementary Data 1). The major protein band, corresponding to PrsA, was detected in the membranes of WT, XcΔ*pgfM2*, and XcΔ*pgtC2* (Fig. 4d, lanes 1, 2, and 3). In contrast, no glycoproteins were recovered from the XcΔ*pgfS* and XcΔ*pgfM1* membranes (Fig. 4d, lanes 4 and 5). Furthermore, increased electrophoretic mobility of PrsA was observed in XcΔ*pgfM2* (Fig. 4d, lane 2).

All identified *S. mutans* glycoproteins were predicted to possess the C-terminal Ser/Thr-rich IDRs (Fig. 2a and Fig. 4e and Supplementary Fig. 8). We confirmed glycosylation of PknB and FtsQ by comparing the electrophoretic mobilities of the proteins in the cell extracts of *S. mutans* WT, XcΔ*pgfS,* and XcΔ*pgfS*:p*pgfS* using protein-specific polyclonal antibodies as we outlined for *S. pyogenes* and *S. pneumoniae* glycoproteins (Fig. 4f, g). To validate PrsA and PBP1a glycosylation and to identify the location of the glycosites, we constructed the 3×FLAG-tagged PrsA and PBP1a variants at their native chromosomal loci in the WT and XcΔ*pgfS* backgrounds. We fused the tag with the C-termini of the full-lengths and the IDR-truncated variants of the proteins, creating PrsA-3×FLAG, PBP1a-3×FLAG, PrsAΔIDR-3×FLAG and PBP1aΔIDR-3×FLAG. Immunoblot analysis with the anti-FLAG antibodies demonstrated that deletion of *pgfS* increased the electrophoretic mobility of the full-length proteins but not the IDR-less variants (Fig. 4h, i). Additionally, we used WFA agarose to purify the glycoproteins from the membranes of WT and the *prsA*ΔIDR mutant lacking the PrsA IDR (the 33 C-terminal amino acid residues). Comparison of the glycoproteins by SDS-PAGE analysis revealed the absence of the band corresponding to PrsA in the *prsA*ΔIDR mutant, confirming a loss of GalNAc moieties (Fig. 2d). Taken together, these experiments conclude that the Pgf pathway enzymes in *S. mutans* glycosylate PrsA, PBP1a, PknB, and FtsQ with GalNAc, and sugar decorations are likely present on the C-terminal Ser/Thr-rich IDRs.

### *S. mutans* protein-based biofilm under ethanol stress depends on *O*-glycosylation and the PrsA IDR

Previous studies have shown that *S. mutans* exhibits defective biofilm formation due to deletion of PrsA, PBP1a, PknB, or BrpA ^44, 79, 80, 81^. To understand the role of protein glycosylation in *S. mutans*, we analyzed the biofilm biomasses of WT, XcΔ*pgfS*, and XcΔ*pgfS*:p*pgfS* grown overnight in a medium supplemented with sucrose or glucose. *S. mutans* grown in a sucrose medium produces an exopolysaccharide-based biofilm. However, when sucrose is replaced with glucose, the bacterium forms a protein-based biofilm mediated by protein-protein interactions of the surface proteins ^82^. The biofilm assay showed that glycosylation is dispensable for exopolysaccharide- or protein-biofilm development in normal conditions (Supplementary Fig. 18a, c). Previously, it has been reported that *S. mutans* remodels biofilm in response to different environmental stresses ^83^. Ethanol, xylitol, caffeine, SOS-inducer mitomycin C, and high temperatures induce the regulatory mechanisms that stimulate the production of a cell wall-attached adhesin responsible for the remodeling of biofilms ^83, 84, 85^. We observed that the addition of 3.5% ethanol into a sucrose medium did not impact the exopolysaccharide-based biofilm of the analyzed strains (Supplementary Fig. 18b). However, when glucose medium was supplemented with ethanol, deletion of *pgfS* significantly impaired the protein-based biofilm (Fig. 5a).

**Fig. 5.**
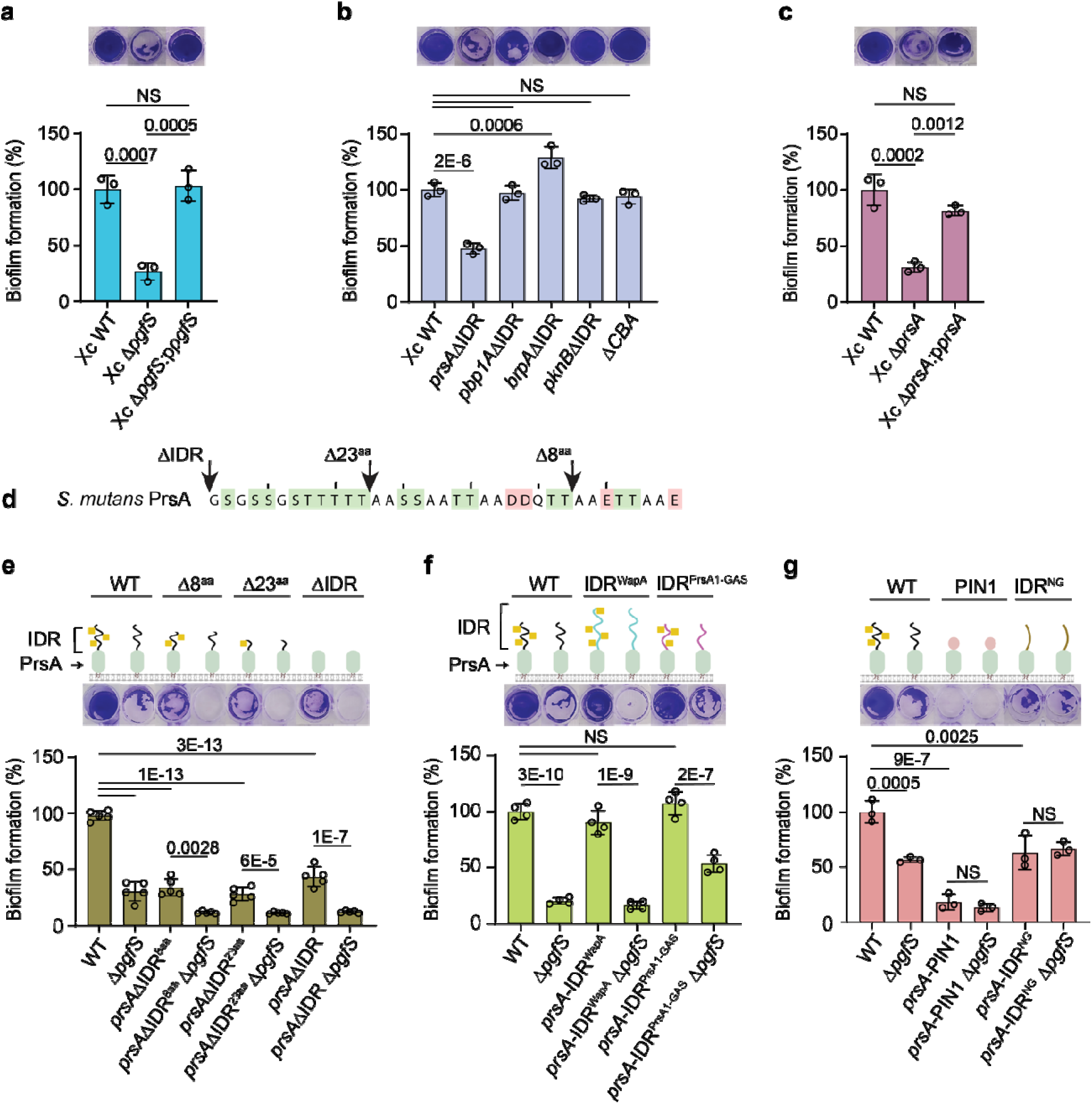
PrsA IDR regulates *S. mutans* protein-based biofilm under ethanol stress. **a**, Biofilm formation of *S. mutans* WT, XcΔ*pgfS,* and XcΔ*pgfS*:p*pgfS*. **b**, Biofilm formation of *S. mutans* strains expressing the IDR-less variants of PrsA, PknB, BrpA, PBP1a (*prsA*ΔIDR, *pknB*ΔIDR, *brpA*ΔIDR and *pbp1a*ΔIDR) or deficient in four cell wall-associated adhesins WapA, Cnm, CnaA and CnaB (ΔCBA). The *pbp1a*ΔIDR mutant is the *pbp1a*ΔIDR-3×FLAG strain. **c**, Biofilm formation of *S. mutans* WT, XcΔ*prsA,* and XcΔ*prsA*:p*prsA*. **d**, Terminal positions of the truncated variants of *S. mutans* PrsA, *prsA*ΔIDR_8 and *prsA*ΔIDR_23, are indicated by arrows. **e**, Biofilm phenotype of *S. mutans* strains expressing PrsA with truncations in the C-terminal IDR. **f**, **g,** Biofilm formation of *S. mutans* strains expressing PrsA variants with the chimeric IDRs. In **e, f** and **g**, schematic representations of PrsA variants are shown (top panel). GalNAc is depicted as yellow squares. In **a**, **b**, **c**, **e**, **f** and **g,** bacterial strains were grown in the UFTYE-glucose medium supplied with 3.5% ethanol for 24 h. Representative biofilms stained with crystal violet are shown in the top panels. Quantifications of biofilm formation are shown in the bottom panels. Columns and error bars represent the mean and S.D., respectively (n = 3 in **a**, **b**, **c** and **g;** n = 4 in **f** and n = 5 in **d** biologically independent replicates. *P* values were calculated by one-way ANOVA with Tukey’s (**a**, **c**, **e**, **f, g**) and Dunnett’s (**b**) multiple comparisons test. Source data for **a, b, c, e, f** and **g** are provided as a Source Data file.

To identify a glycoprotein linked to this phenotype, we analyzed biofilm formation of the PrsA, PknB, BrpA, and PBP1a mutants lacking C-terminal IDRs (*prsA*ΔIDR, *pknB*ΔIDR, *brpA*ΔIDR, and *pbp1a*ΔIDR-3×FLAG), and the ΔCBA mutant deficient in all four collagen-binding adhesins, WapA, CnaB, CbpA, Cnm, that are likely glycosylated by the Pgf pathway. Similar to XcΔ*pgfS*, the *prsA*ΔIDR mutant produced biofilm in the normal conditions but was biofilm-defective in the ethanol-stressed conditions (Fig. 5b, Supplementary Fig. 19a). The *prsA*ΔIDR and XcΔ*pgfS* strains were not attenuated for growth in the presence of ethanol (Supplementary Fig. 20), indicating that a significant decline of the biofilm biomass of these mutants is not due to the change in cell growth. The isogenic *prsA* deletion mutant, XcΔ*prsA*, was also biofilm-defective only in the ethanol-stressed conditions (Fig. 5c, Supplementary Fig. 19b). However, this mutant displayed a modest growth defect (Supplementary Fig. 20), indicating that PrsA is also necessary to support cell growth under ethanol stress. Furthermore, deletion of the C-terminal 8 or 23 amino acid residues of PrsA (*prsA*ΔIDR_8 and *prsA*ΔIDR_23, respectively, Fig. 5d) also resulted in biofilm deficiency only in the ethanol-stressed conditions (Fig. 5e, Supplementary Fig. 19c). Based on these results, we concluded that the full-length PrsA IDR is critical for *S. mutans* protein-based biofilm development under ethanol stress. Interestingly, a more prominent biofilm defect was observed when *pgfS* was deleted in the *prsA*ΔIDR_8, *prsA*ΔIDR_23, and *prsA*ΔIDR strains, implying that *O*-glycosylation maintains biofilm synthesis not only via the PrsA IDR (Fig. 5e).

### C-terminal disorder and *O*-glycosylation are required for PrsA function in protein-based biofilm under ethanol stress

To understand if C-terminal disorder is critical for PrsA function in protein-based biofilm development in the ethanol-stressed conditions, we replaced the PrsA IDR from its *S. mutans* chromosomal locus with a) the 74-residue Ser/Thr-rich region of WapA (hereafter PrsA-IDR^WapA^) which is likely *O*-glycosylated by the Pgf mechanism ^34, 35^, b) the 48-residue PrsA1 IDR of *S. pyogenes* which is glycosylated by PgtC2 (hereafter PrsA-IDR^PrsA1_GAS^), c) the 34-residue PIN1 WW domain ^86^ which was fused with PrsA via a short peptide linker (hereafter PrsA-PIN1). The *S. mutans* PrsA IDR shares a very low sequence homology with the WapA and *S. pyogenes* PrsA1 IDRs, and no sequence homology to PIN1 (Supplementary Fig. 21a, b). PIN1 domain has six Ser and Thr residues, and it is one of the smallest structured proteins adopting a β-sheet fold (Supplementary Fig. 21b, c, d) ^86^. Finally, to understand if sugar decorations on the PrsA IDR are necessary for PrsA function in biofilm formation, we constructed mutants in the *S. mutans* WT and XcΔ*pgfS* backgrounds in which the PrsA IDR was mutated to replace Ser and Thr residues with asparagine and glycine residues (hereafter PrsA-IDR^NG^, Supplementary Fig. 21e). Similar to the PrsA IDR, IDR^NG^ is also predicted as a highly disordered region by DR-BERT, disorder predictors assembled into the RIDAO platform and structure prediction by AlphaFold2 (Supplementary Fig. 21f, g). Strikingly, the WT background strains expressing the PrsA-IDR^WapA^ or PrsA-IDR^PrsA1_GAS^ chimeras produced protein-based biofilm under ethanol stress. But, the deletion of *pgfS* in these strains resulted in reduced protein-based biofilm only in the ethanol-stressed conditions (Fig. 5f, Supplementary Fig. 19d). In contrast, the WT and XcΔ*pgfS* strains expressing the PrsA-PIN1 chimera or PrsA-IDR^NG^ displayed a significant defect in biofilm biomass in the presence of ethanol (Fig. 5g, Supplementary Fig. 19e). The impact was more dramatic in the strains expressing the PrsA-PIN1 chimera. Taken together, these data demonstrate that both the C-terminal disorder and *O*-glycosylation are necessary for the function of PrsA in *S. mutans* protein-based biofilm development under ethanol stress.

### Glycosylation protects some IDRs from proteolytic cleavage

Remarkably, examination of PrsA expression in *S. mutans* using the anti-PrsA polyclonal antibodies revealed that PrsA migrates as a broad band at around 43-47 kDa in WT and XcΔ*pgfS*:p*pgfS*, but as two bands at around 38 and 40 kDa in XcΔ*pgfS* (Fig. 6a, lanes 1-3). We hypothesized that the 40 kDa band (lane 2, red arrow) is a non-glycosylated form of PrsA and the 38 kDa band (lane 2, black circle) is a proteolytically processed form of PrsA which was not detected in the 3×FLAG-tagged strains by the anti-FLAG antibodies (Fig. 4h). Upon exposure of cells to 3.5% ethanol, heterogeneity of PrsA was significantly increased in WT and XcΔ*pgfS*:p*pgfS*. The protein migrated between 38 and 47 kDa (Fig. 6a, lanes 4 and 6). Bands corresponding to the non-glycosylated and proteolytically processed forms of PrsA were also detected in these cells, implying that ethanol inhibits glycosylation. These observations were further corroborated by the analysis of PrsA in the *S. mutans* strains harboring the PrsA-3×FLAG variant (Fig. 6b). Immunoblot analysis with the anti-FLAG antibodies demonstrated that the ethanol-subjected WT cells produced an additional low molecular weight band (Fig. 6b, lane 3) which co-migrates with the non-glycosylated form of PrsA-3×FLAG (Fig. 6b, lane 2, red arrow). When we used the anti-PrsA antibodies to assess the PrsA-3×FLAG variant in the cells grown under normal conditions, the protein migrated as two bands in the XcΔ*pgfS* background (Fig. 6c, lane 2), consistent with the idea that the smallest fragment (black circle) is formed by proteolytic cleavage. Upon exposure of the cells to ethanol stress, the PrsA-3×FLAG cleavage product was also detected in the WT background (Fig. 6c, lane 3), which aligns with the hypothesis that ethanol inhibits protein glycosylation. Because the smallest fragment of the PrsA-3×FLAG variant is not detected by the anti-FLAG antibodies, it confirms that this fragment is a proteolytically processed form of PrsA lacking the 3×FLAG tag. These observations also argue that the cleavage site for an unknown protease is present on the PrsA IDR and *O*-glycosylation protects this region from proteolysis.

**Fig. 6.**
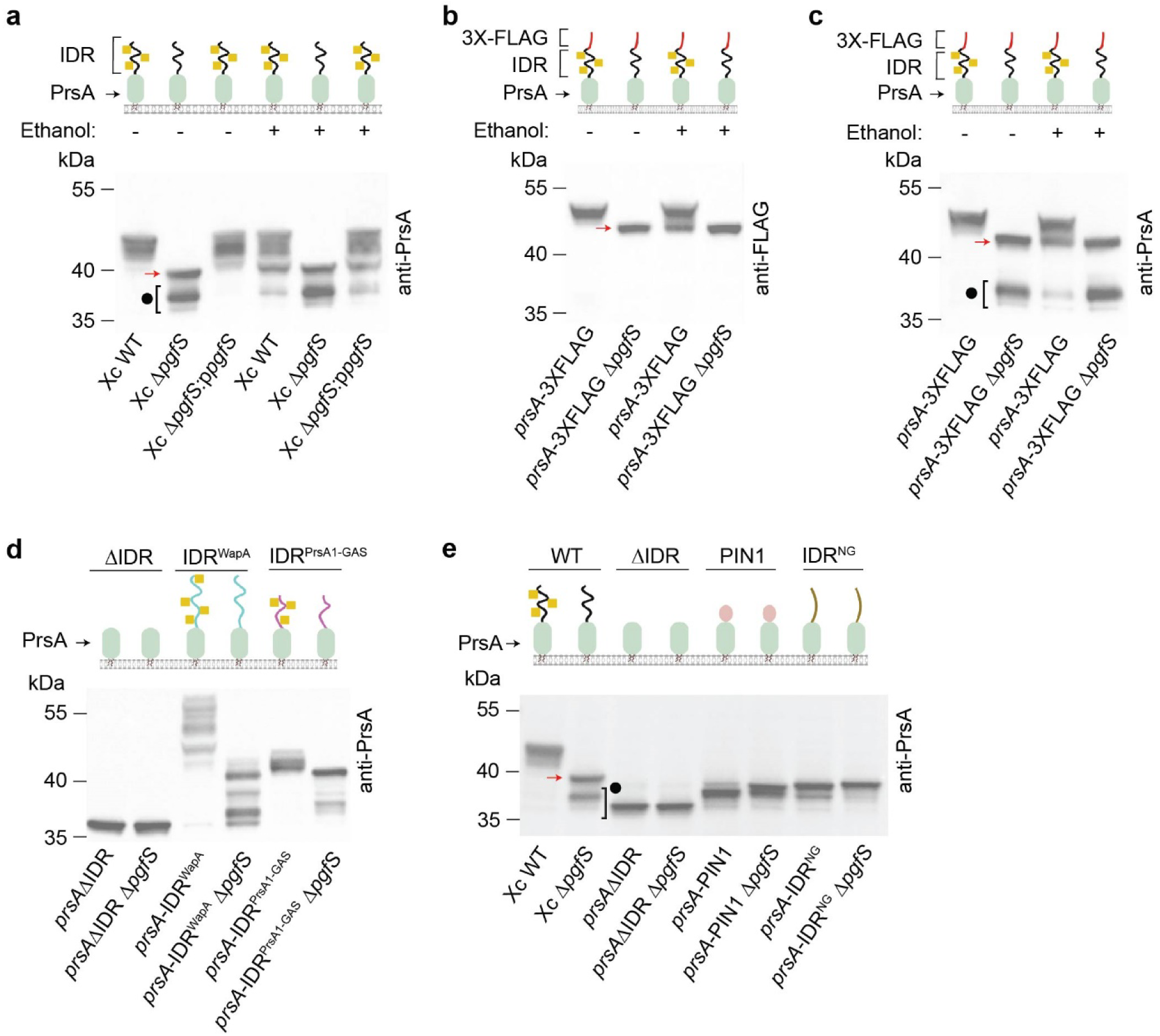
*O*-glycosylation protects the PrsA IDR from proteolysis in *S. mutans*. **a**, Immunoblot analysis of PrsA in WT, XcΔ*pgfS,* and XcΔ*pgfS*:p*pgfS* grown in normal or ethanol-stressed conditions. PrsA is probed with anti-PrsA antibodies. **b**, **c**, Immunoblot analysis of 3×FLAG-tagged variant of PrsA in the WT and XcΔ*pgfS* backgrounds grown in normal or ethanol-stressed conditions. PrsA is probed with anti-FLAG antibodies in (**b**) and anti-PrsA antibodies in (**c**). **d**, **e**, Immunoblot analysis of PrsA variants harboring the chimeric IDRs in the WT and XcΔ*pgfS* backgrounds grown in normal conditions. PrsA is probed with anti-PrsA antibodies. Proteins were separated on a 4-12% SurePAGE™ Bis-Tris gel (Genscript) in MES buffer in **a**, **b**, **c**, **d** and **e**. Schematic of PrsA variants is shown in **a**, **b**, **c**, **d**, and **e** top panels. GalNAc is depicted as yellow squares. In **a**, **b**, **c**, and **e**, the red arrow indicates a non-glycosylated form of PrsA and the black circle denotes the proteolytic products of PrsA. The experiments were performed independently three times in **a**, **b**, **d**, **e**, and two times in **c**, and they yielded the same results. A representative image from one experiment is shown. Source data for **a, b, c, d** and **e** are provided as a Source Data file.

Further investigation of the PrsAΔIDR and PrsA-IDR^NG^ variants, and three PrsA-IDR^WapA^, PrsA-IDR^PrsA1_GAS^, and PrsA-PIN1 chimeras using the anti-PrsA antibodies revealed that PrsAΔIDR migrated as a single sharp band (Fig. 6d, lane 1 and Fig. 6e, lane 3), PrsA-IDR^WapA^ migrated as multiple bands (Fig. 6d, lane 3), and PrsA-IDR^PrsA1_GAS^, PrsA-PIN1, and PrsA-IDR^NG^ migrated as broad bands (Fig. 6d, lane 5 and Fig. 6e, lanes 5 and 7). The non-glycosylated forms of PrsA-IDR^WapA^ and PrsA-IDR^PrsA1_GAS^ in the XcΔ*pgfS* strain background demonstrated multiple bands of greater mobility (Fig. 6d, lanes 4 and 6), suggesting a proteolytic cleavage of the IDRs. These data provide evidence that IDR^WapA^ and IDR^PrsA1_GAS^ are glycosylated in the WT strain background and sensitive to proteolysis when *O*-glycans are removed. In contrast, in the absence of glycosylation, the electrophoretic mobility of PrsAΔIDR, PrsA-PIN1, and PrsA-IDR^NG^ did not increase (Fig. 6d, lane 2 and Fig. 6e, lanes 4, 6, and 8) indicating that these PrsA variants are not decorated with sugars. We noted that the PrsA-PIN1 and PrsA-IDR^NG^ variants generated double bands, suggesting the proteolytic vulnerability of these proteins. To test this idea, we fused the C-terminus of PrsA-IDR^NG^ with the 3×FLAG tag, creating PrsA-IDR^NG^-3×FLAG. Immunoblot analysis with the anti-PrsA antibodies revealed that this variant displayed the major and minor bands in the WT or XcΔ*pgfS* backgrounds (Supplementary Fig. 22a). The minor band was not detected when the blot was probed with the anti-FLAG antibodies (Supplementary Fig. 22b). Moreover, the electrophoretic mobility of this PrsA variant was similar in both strain backgrounds validating that this protein does not carry sugar moieties (Supplementary Fig. 22a, b). Hence, these data confirm the sensitivity of the IDRs to proteolysis in the absence of sugar decorations.

Considering these observations, we examined the expression of other *S. mutans* glycoproteins in the WT and XcΔ*pgfS* backgrounds. Additional lower-molecular weight PknB-specific bands were detected in XcΔ*pgfS*, suggesting the function of *O*-glycosylation in protecting *S. mutans* PknB from proteolysis (Fig. 4f, lane 2, black circle).

### Some C-terminal IDRs allow for optimal protein levels

We considered that the chaperone function of PrsA likely safeguards extracellular proteins from misfolding and aggregation in proteotoxic conditions when ethanol is present in the growth medium. Therefore, we investigated whether the PrsA IDR promotes the folding of PrsA. We first explored the possibility that the IDR affects the secondary structure and the overall thermal stability of PrsA. We expressed in *E. coli* two recombinant soluble variants of PrsA that lack the signal peptide: the full-length version, ePrsA-FL, and the IDR-less version, ePrsAΔIDR. Both variants displayed similar behavior in protein thermal stability assays conducted in a wide range of buffers and salts, including 3.5% ethanol (Supplementary Fig. 23a, b), indicating that the IDR does not affect the thermal stability of PrsA. Moreover, the circular dichroism spectra of ePrsA-FL and ePrsAΔIDR did not reveal any prominent structural changes in the secondary structure content between these variants (Supplementary Fig. 23c).

We next examined the possibility that the IDR facilitates the folding and stability of PrsA following its transport across the *S. mutans* plasma membrane. Comparison of the total protein levels of the full-length FLAG-tagged PrsA with the IDR-less variant, analyzed using the anti-FLAG antibodies, revealed the positive effect of the IDR on PrsA production (Fig. 4h, Supplementary Fig. 24a). Notably, similar results were also obtained when we compared the PBP1a protein level in the PBP1a-3×FLAG and PBP1aΔIDR-3×FLAG strains using the anti-FLAG antibodies (Fig. 4i, Supplementary Fig. 24b) and the PknB protein level in the *S. mutans* WT and *pknB*ΔIDR using the anti-PknB antibodies (Supplementary Fig. 25). These data, together with the analysis of *S. pneumoniae* PBP1a (Supplementary Fig. 16b), establish that some C-terminal IDRs positively contribute to optimal protein levels.

To understand the biofilm-deficient phenotype of the *prsA*ΔIDR_8 and *prsA*ΔIDR_23 and *prsA*ΔIDR mutants, we analyzed the full-length PrsA, PrsAΔIDR_8, PrsAΔIDR_23, and PrsAΔIDR variants in the WT and XcΔ*pgfS* backgrounds using the anti-PrsA antibodies. Strikingly, PrsAΔIDR_8 and PrsAΔIDR_23 produced the major and minor bands of approximately 37 kDa in both strain backgrounds (Fig. 7a, lanes 3-6, black circle), indicating that even the removal of 8 amino acid residues from the IDR significantly inhibits the efficiency of glycosylation leading to increased proteolysis of PrsA. Moreover, the glycosylated form of PrsAΔIDR_8 was more heterogenous than the full-length PrsA, supporting the notion that this mutant has altered glycosylation (Fig. 7a, lanes 3). Quantification of all PrsA species in *S. mutans* WT and the strains with C-terminal truncations of the PrsA IDR revealed that only deletion of the 23 C-terminal amino acid residues in the XcΔ*pgfS* background and the 33 C-terminal amino acid residues in both backgrounds significantly decreased the PrsA protein level (Fig. 7b). These data confirm a positive role of the PrsA IDR for PrsA production in *S. mutans* and highlight the importance of the optimal length of the PrsA IDR for its efficient glycosylation.

**Fig. 7.**
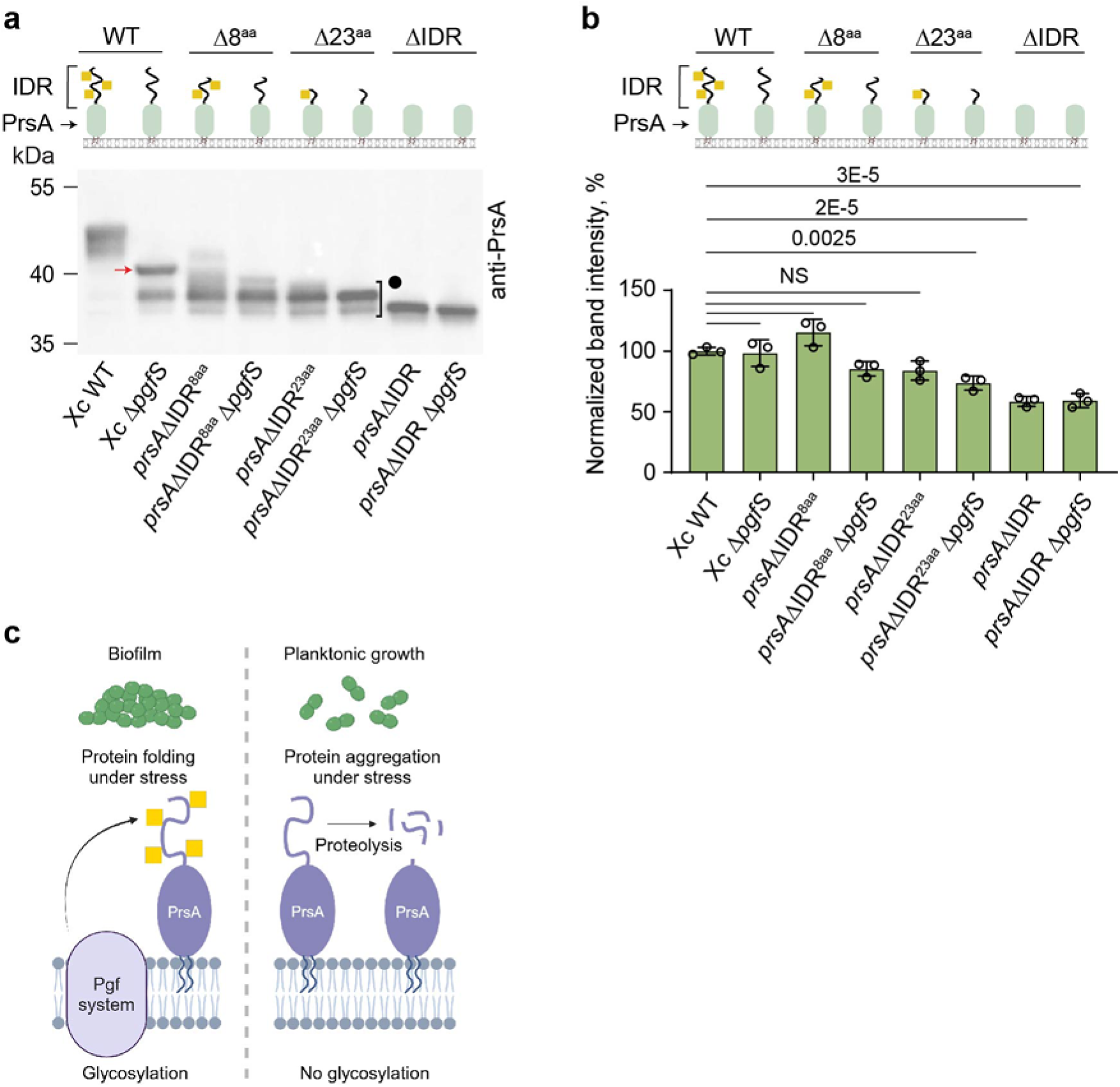
Function of *O*-glycosylation in the PrsA proteolytic stability and biofilm development in *S. mutans*. **a**, Immunoblot analysis of PrsA variants carrying the truncated IDRs in the WT and XcΔ*pgfS* backgrounds grown in normal conditions. PrsA is assessed by anti-PrsA antibodies. The red arrow and the black circle denote a non-glycosylated form- and the proteolytic products of PrsA, respectively. The experiments were performed independently three times and they yielded the same results. A representative image from one experiment is shown. **b,** Relative band intensities of *S. mutans* PrsA variants with the truncated IDRs. The representative immunoblot is shown in **a**. Total band intensities from each blot lane were calculated using ImageJ, and the data were normalized. Columns and error bars represent the mean and S.D., respectively (n = 3 biologically independent replicates). *P* values were determined by one-way ANOVA with Dunnett’s multiple comparisons test. In **a**, and **b**, schematic of PrsA variants is shown in top panel. GalNAc is depicted as yellow squares. **c**, The PrsA C-terminal IDR plays a functional role in *S. mutans* biofilm formation in ethanol-stressed cells. The Pgf-dependent protein *O*-glycosylation machinery covalently attaches GalNAc moieties to the C-terminal IDR of the post-translocation secretion chaperone PrsA. This functional form of PrsA prevents misfolding of protein clients in ethanol-stressed conditions supporting the development of a protein-based biofilm. Loss of *O*-glycosylation exposes the IDR to proteolytic attack, resulting in a non-functional PrsA, which does not aid in biofilm formation. Source data for **a** and **b** are provided as a Source Data file.

Based on the above results, we propose a model to explain the biofilm phenotype of these mutants (Fig. 7c). The C-terminal disordered region of PrsA might be necessary for preventing the misfolding and aggregation of PrsA protein clients involved in biofilm formation in proteotoxic stress conditions. Sugar decorations on the PrsA IDR protect this important region from degradation by an unknown protease and might also stimulate the chaperone function of PrsA.

## Discussion

Our glycobiological studies, in combination with genome-wide identification of IDRs, uncovered a unique class of IDRs linked to the extracytoplasmic proteins in streptococci (Supplementary Table 3). A distinct feature of these IDRs is the presence of poly-Ser/Thr residue tracks. In *S. pyogenes* and *S. pneumoniae*, we elucidated a novel Glc-type protein glycosylation mechanism in which GtrB works in pair with PgtC2 to transfer α-Glc to Ser/Thr-rich IDRs. However, we have not observed the function of the GtrB/PgtC2 mechanism in glycosylation of *S. mutans* membrane-associated proteins. Our study demonstrates that the glycosyltransferases of the Pgf protein glycosylation pathway — PgfS and PgfM1 — transfer GalNAc to Ser/Thr-rich IDRs tethered to membrane-associated proteins in *S. mutans* (Supplementary Fig. 26). This final step of sugar transfer very likely occurs in the periplasm because a catalytic DXD motif in PgtC2 and PgfM1 enzymes is located in extramembrane loop regions (Supplementary Fig. 26), consistent with other bacterial members of this superfamily of glycosyltransferases ^87, 88^. The lectin-affinity glycoprotein enrichment strategy impartially revealed a strong correlation between this subset of IDRs and glycosylation. The presence of glycans on IDRs was further confirmed by comparing the molecular weights of glycoproteins and their IDR-less variants in the WT and glycosylation-deficient backgrounds and by inspecting the binding of different protein variants to specific lectins. The Pgf mechanism was previously shown to *O*-glycosylate Ser residues on Cnm adhesin ^34^. In this study, using the PrsA IDR, we further confirm the function of this mechanism in *O*-glycosylation of Ser/Thr-rich IDRs. Moreover, a structural homology of PgfM1 to *S. pyogenes* and *S. pneumoniae* PgtC2 (Supplementary Fig. 17a), as well as the tight binding of *S. pyogenes* and *S. pneumoniae* glycoproteins to ConA lectin, argue that the numerous Ser/Thr residues on IDRs in these bacteria are also heavily *O*-glycosylated.

Strikingly, the identified IDRs tend to localize at the C-termini of membrane-associated proteins involved in critical cellular processes such as protein folding, cell wall biosynthesis, cell division, and signal transduction. Among the proteins harboring this type of IDRs, we identified a conserved post-translational chaperone lipoprotein PrsA, class A penicillin-binding proteins PBP1a and PBP1b, LytR-CpsA-Psr glycopolymer transferase LCP involved in the attachment of polysaccharides to peptidoglycan, serine/threonine protein kinase PknB, the extracytoplasmic regulator of c-di-AMP synthesis CdaR, and a divisome protein FtsQ. Sequence alignments of identified glycoproteins with the bacterial orthologs provide evidence that this class of IDRs occurs universally in streptococci and is present in other Firmicutes, suggesting *O*-glycosylation of these regions. In *B. subtilis*, the major penicillin-binding protein PBP1 also harbors a C-terminal IDR, which has been shown to enhance cell wall biosynthesis ^24^. PBP1 has been detected as two isoforms, PBP1a and PBP1b, which are different by molecular weight but encoded by the same gene, *ponA* ^89^. Intriguingly, an alternative form of PBP1 is formed by post-translational processing of the PBP1 IDR by an unknown mechanism ^89^. Although the *B. subtilis* PBP1 IDR lacks the poly-Ser/Thr, it has Ser/Thr residues and a large number of asparagine residues (Supplementary Fig. 12), implying that PBP1a is generated by *O*- or *N*-glycosylation of PBP1b. This example and our study argue that C-terminal IDRs modulate the respective functions of adjacent structured domains.

To understand the biological function of Ser/Thr-rich IDRs, we focused on *S. mutans*, a major etiological agent of human dental caries. Dental caries progression involves the attachment of *S. mutans* to the tooth surface and the development of virulent dental plaque. Surface proteins play a key role in these processes. We found that under ethanol stress, the formation of protein-based biofilm is dependent on *O*-glycosylation of the PrsA IDR. While the PrsA IDR is dispensable for *S. mutans* biofilm formation in the absence of stress, it becomes critical upon ethanol exposure. The crystal structures of *B. subtilis* and *Listeria monocytogenes* PrsA homologs demonstrated that the enzymes possess two functional subdomains, the chaperone-like, and the peptidyl-prolyl isomerase domain ^90, 91^. It has been proposed that PrsA has a dual role in protein folding by providing a chaperone function to control the aggregation of protein clients and catalyzing the cis/trans isomerization of peptide bonds with proline residues ^90, 91^. The chaperone function of PrsA is likely critically important in ethanol-stressed cells when misfolding of proteins is increased. By generating the PrsA variants in which Ser/Thr residues of the IDR were mutated, or this region was swapped with the structured peptide or the glycosylated IDRs, we determined that both the C-terminal disorder and *O*-glycosylation are critical for the functionality of PrsA in protein-based biofilm under ethanol stress.

It is currently unknown how this region contributes to PrsA activity. Interestingly, IDRs are abundant and functionally important in different folding chaperones, including *E. coli* DnaK, GroEL, HtpG, Hsp33, and HdeA ^92^. The disordered regions of some folding chaperones enable them to bind multiple clients, preventing toxic aggregation. If the PrsA IDR binds protein clients, the interactions should involve the sugar residues. Another possible function of the PrsA IDR is to increase protein stability and the folding of PrsA through steric hindrance acting as a so-called “entropic bristle” by entropically excluding large neighboring molecules ^17, 93^. There is also growing evidence that protein N-glycosylation in eukaryotes positively affects protein folding and stability by hindering hydrophobic aggregation-prone regions and reducing the conformational entropy of proteins ^94, 95^. Thus, the dual effect of disorder and glycosylation might substantially enhance the folding of adjacent domains. Our data demonstrating that the IDRs enhance the protein levels of *S. mutans* PrsA, PBP1a, PknB, and *S. pneumoniae* PBP1a and MapZ, and glycosylation positively impacts the production of *S. pneumoniae* MapZ, RodZ, Pbp1b support this idea. Because PrsA is a folding chaperone, it can be expected that the PrsA IDR also slows the aggregation of protein clients, thereby promoting their correct folding.

Our work also showed that the absence of *O*-glycosylation in *S. mutans* causes proteolytic degradation of the PrsA IDR, including the chimeric IDRs, supporting the idea that *O*-glycans shield proteins from membrane and extracellular proteases. The efficiency of PrsA glycosylation drops with the reduction of the IDR length, leading to increased degradation of the IDR and, as a result, to a biofilm defect. This effect of glycosylation is likely due to obstructing proteases through steric hindrance. Thus, *S. mutans* have evolved a sophisticated mechanism to regulate the function of PrsA by the C-terminal IDR, which carries diverse features, including the cleavage sites for an unknown protease, the *O*-glycosites, and the optimal length of the IDR. Furthermore, these features are not restricted to PrsA because *S. mutans* PknB lacking GalNAc decorations also displays increased proteolytic processing.

Another key finding of our study is that glycosylation of the PrsA IDR is suppressed in the ethanol-stressed *S. mutans* cells, probably because the last step of glycan transfer, which occurs in the periplasmic space, is highly vulnerable to small toxic molecules such as ethanol that can pass through the porous peptidoglycan. This observation highlights the tunable nature of IDRs to external stimuli and pinpoints a direct mechanism bacteria can use to modulate the homeostasis of extracytoplasmic proteins in response to specific environments. Future investigation of the functions of IDRs identified in our study may provide fundamentally new insights into how these regions sense and respond to environmental changes to boost the survival of bacteria. Finally, current advances in deep neural network-based methods have presented the opportunity to systematically evaluate IDRs in other prokaryotes. We anticipate that these approaches may expand our observation to multiple bacterial species and lead to the discovery of novel functional motifs in the prokaryotic IDRs.

## Methods

### Bacterial strains, growth conditions, and media

All plasmids, strains, and primers used in this study are listed in Supplementary Tables 4-8. *S. pyogenes* MGAS 2221*, S. mutans* Xc, and *S. pneumoniae* strains D39 and TIGR4 were used in this study. *S. pneumoniae* D39 strains were derived from unencapsulated strains IU1824 (D39 Δ*cps rpsL1*), which were derived from the encapsulated serotype-2 D39W progenitor strain IU1690 ^98, 99^. Unless otherwise indicated, *S. pyogenes* and *S. mutans* were grown in BD Bacto Todd-Hewitt broth supplemented with 1% yeast extract (THY) without aeration at 37 °C. *S. pneumoniae* was cultured statically in Becton-Dickinson brain heart infusion (BHI) broth at 37 °C in an atmosphere of 5% CO_2_, and growth was monitored by OD_620_ as described before ^100^. In all experiments, cells were inoculated from frozen glycerol stocks into BHI broth, serially diluted, and incubated for 12–15 h. The next day, cultures at OD_620_ ∼0.1–0.4 were diluted to OD_620_ ∼0.005 in BHI broth. *S. pneumoniae* D39 cultures were sampled for western analysis at OD_620_ ∼0.15–0.2 (early to mid-exponential phase). *E. coli* strains were grown in Lysogeny Broth (LB) medium or on LB agar plates at 37 °C. When required, antibiotics were included at the following concentrations: erythromycin at 500 µg mL^-1^ for *E. coli* and 5 µg mL^-1^ for streptococci; chloramphenicol at 10 µg mL^-1^ for *E. coli* and 4 µg mL^-1^ for streptococci; spectinomycin at 200 µg mL^-1^ for *E. coli* and 300 µg mL^-1^ for streptococci; kanamycin at 50 µg mL^-1^ for *E. coli* and 300 µg mL^-1^ for streptococci.

### Construction of mutants in streptococci

To construct 2221Δ*gtrB*, we used a PCR overlapping mutagenesis approach, as previously described ^53^. Briefly, 600-700 bp fragments both upstream and downstream of the gene of interest were amplified from *S. pyogenes* 2221 chromosomal DNA with designed primers that contained 16-20 bp extensions complementary to a nonpolar kanamycin resistance cassette which was PCR-amplified from pOSKAR ^101^. Two fragments flanking the gene of interest and kanamycin cassette were purified using the QIAquick PCR purification kit (Qiagen) and fused by Gibson Assembly (SGA-DNA). A PCR was then performed on the Gibson Assembly sample using the primers listed in Supplementary Table 7 to amplify the fused fragments. The assembled DNA fragments for mutagenesis of GAS *gtrB* were cloned into pMBsacB ^102^ to generate the plasmid pMBsacB-gtrB. The plasmid was electroporated into the GAS 2221 cells, and the transformants were selected on THY agar containing erythromycin. A two-step temperature-dependent selection process was used to isolate the double-crossover mutants that are sensitive to erythromycin but resistant to kanamycin ^102^. The *S. mutans* mutants were constructed using a similar PCR overlapping mutagenesis approach. The primers used to amplify 600-700 bp fragments for Gibson Assembly are listed in Supplementary Table 7, and the antibiotic resistance cassettes are listed in Supplementary Table 4. The assembled DNA fragments were directly transformed into the *S. mutans* Xc cells by electroporation. The transformants were selected on THY agar containing the corresponding antibiotic. Double-crossover recombination in the *S. pyogenes* and *S. mutans* mutant clones was confirmed by PCR and Sanger sequencing using the primers listed in Supplementary Table 7. Some deletion mutants were verified by whole-genome sequencing (Supplementary Table 9). For complementation of *S. pyogenes* and *S. mutans* mutants, the promoterless genes were amplified from the corresponding bacterial strains using the primers listed in Supplementary Table 7. The PCR products were ligated into pDC123 vector. The resultant plasmids (Supplementary Table 4) were transformed into the corresponding mutants by electroporation. Chloramphenicol-resistant single colonies were picked and checked for the presence of the transformed gene using PCR and Sanger sequencing.

The *S. pneumoniae* mutants (Supplementary Table 5) were constructed using the background strains, primers and DNA templates listed in Supplementary Tables 6 and 8. *S. pneumoniae* strains containing antibiotic markers were constructed by transformation of CSP1-induced competent pneumococcal cells with linear DNA amplicons synthesized by overlapping fusion PCR ^100, 103, 104^. Strains containing markerless alleles in native chromosomal loci were constructed using allele replacement via the P_c_-[*kan*-*rpsL*^+^] (Janus cassette) ^105^. Bacteria were grown on plates containing trypticase soy agar II (modified; Becton-Dickinson), and 5% (vol/vol) defibrinated sheep blood (TSAII-BA). TSAII-BA plates used for selection were incubated at 37 °C in an atmosphere of 5% CO_2_ and contained antibiotics at concentrations described previously ^100, 104^. Ectopic expression *gtrB* for complementation was achieved with a constitutive P*_ftsA_* promoter in the ectopic *bgaA* site ^106^. All constructs were confirmed by PCR and DNA sequencing of chromosomal regions corresponding to the amplicon region used for transformation.

### Construction of the plasmids for protein expression in *E. coli* LOBSTR(DE3) cells

The genes encoding the extracellular domain of *S. mutans* PrsA (hereafter ePrsA-FL) and the IDR truncated version of the extracellular domains of *S. mutans* PrsA, FtsQ and PknB (hereafter ePrsAΔIDR, eFtsQΔIDR and ePknBΔIDR^Smu^) were PCR-amplified from *S. mutans* Xc. The gene encoding the IDR truncated version of the extracellular domain of *S. pyogenes* PknB (hereafter ePknBΔIDR^Spy^) was PCR-amplified from *S. pyogenes* 2221. The corresponding primers used for cloning are listed in Supplementary Table 7. The PCR products were digested by the corresponding restriction enzymes and subcloned into a pRSF-NT vector. The resultant plasmids (Supplementary Table 4) express genes fused at the N-terminus with a His-tag followed by a TEV protease recognition site. They were transferred into competent *E. coli* LOBSTR(DE3) (Kerafast)

### Construction of the plasmid for expression of GtrB^Spy^ in *E. coli* JW2347

To create a vector for expression of GtrB^Spy^, the gene was amplified from *S. pyogenes* 2221 chromosomal DNA using the primer pair listed in Supplementary Table 7. The PCR product was digested by the corresponding restriction enzymes and subcloned into a pBAD33 vector. The resultant plasmid, pBAD33_GtrB^Spy^ was transferred into competent *E. coli* JW2347 strain that has a deletion of the *gtrB* gene ^52^.

### Overexpression and purification of recombinant proteins from *E. coli*

To purify recombinant proteins, cells were grown to exponential phase (OD_600_ ∼0.6) at 37 °C in an LB medium supplemented with kanamycin and chloramphenicol. Protein overexpression was induced by adding 0.25 mM isopropyl β-*D*-1-thiogalactopyranoside (IPTG), and growth was continued for approximately 18 h at 18 °C. Cells were harvested by centrifugation and lysed in 20 mM Tris-HCl pH 7.5, 300 mM NaCl, 1× bacterial protease inhibitor cocktail (Pierce Chemical Co.), 1 µg mL^-1^ DNase, and 25 µg mL^-1^ lysozyme using a microfluidizer cell disrupter. The soluble fractions were purified by Ni-NTA chromatography as previously reported for ePplD ^107^. The eluate was dialyzed overnight into 20 mM Tris-HCl pH 7.5 and 300 mM NaCl to remove imidazole. Protein concentration was determined by Pierce™ BCA protein assay kit (Thermo Fisher Scientific) and purity was estimated by SDS-PAGE.

### Preparation of *S. pyogenes* and *E. coli* JW2347 membrane fractions to assess GtrB^Spy^ activity

*S. pyogenes* strains grown in THY medium to an OD_600_∼0.8 were collected by centrifugation at 18,000 g for 10 min. Cells were resuspended in 50 mM HEPES pH 7.4, 0.6 M sucrose, recombinant PlyC (50 µg mL^-1^), and incubated at 37 °C for 1 hr. The cell suspensions were rapidly diluted with 100 mL ice-cold 10 mM HEPES pH 7.4, 2 mM MgCl_2_, 1× bacterial protease inhibitor cocktail, 5 µg mL^-1^ DNAse, 5 µg mL^-1^ RNAse, and incubated on ice for 10 min. The suspension was homogenized with a Dounce homogenizer followed by adjustment of sucrose and EDTA in the suspension to a final concentration of 0.25 M and 10 mM, respectively. The suspension was centrifuged (2,500 g, 10 min, 4 °C) followed by centrifugation of the supernatant (40,000 g, 30 min, 4 °C). The resulting pellet that contained the membrane fraction was resuspended in 10 mM HEPES pH 7.4, 0.25 M sucrose, 1 mM EDTA.

For expression of GtrB^Spy^, *E. coli* JW2347 cells carrying pBAD33_GtrB^Spy^ plasmid were grown in LB at 37 °C, and induced with 13 mM L-arabinose at OD_600_ ∼0.8, followed by 3 hours of growth at 25 °C. The cells were lysed in 20 mM Tris-HCl pH 7.5, 300 mM NaCl with two passes through a microfluidizer cell disrupter. The lysate was centrifuged (1000 g, 15 minutes, 4 °C), followed by centrifugation of the supernatant (40,000 g, 60 min, 4 °C) to isolate the membrane fraction. A similar approach was used to prepare the membrane fraction of *E. coli* JW2347 cells carrying an empty plasmid (pBAD33).

### *In vitro* analysis of Glc-P-Und synthase activity

Reactions for the assay of Glc-P-Und synthase activity contained 50 mM HEPES pH 7.4, 20 mM MgCl_2_, 1 mM DTT, 1 mM sodium orthovanadate, the indicated amounts of UDP-[^3^H]Glc (50-500 cpm/pmol), Und-P, CHAPS and bacterial enzyme (10-25 μg membrane protein from either *S. pyogenes* or *E. coli*) in a total volume of 0.02 mL. Where indicated, Und-P was added as a sonicated dispersion in 1% CHAPS, with a final reaction concentration of 0.35% CHAPS. Following incubation at 37 °C for 10-20 min, reactions were stopped by the addition of 2 mL CHCl_3_/CH_3_OH (2:1). Reactions were freed of unincorporated radioactivity by Folch partitioning as described previously ^49^ and analyzed for radioactivity by scintillation spectrometry or by TLC on silica gel G developed in CHCl_3_/CH_3_OH/H_2_O/NH_4_OH (65:30:4:1). Reactions with membrane fractions from *S. pyogenes* were subjected to mild alkaline de-acylation with 0.1 N KOH in methanol/toluene (3:1) for 60 min at 0 °C, to destroy [^3^H]glucosylglycerolipids and freed of released radioactivity by partitioning as described previously ^49^. Degradation of Glc-lipids by mild acid and mild alkaline treatment was conducted as previously described ^49^.

### Co-sedimentation assay with fluorescent lectins

*S. mutans* cells grown overnight (∼17 hours) were harvested by centrifugation and washed three times with 1× phosphate-buffered saline (PBS). The cell pellet was resuspended in 0.5 mL of PBS (OD_600_ ∼ 9). 10-50 µg mL^-1^ of FITC-conjugated lectin from *Bandeiraea simplicifolia* (Sigma-Aldrich), Alexa Fluor™ 488-conjugated lectin from *Helix pomatia* (Thermo Fisher Scientific), or FITC-conjugated lectin from *Wisteria floribunda* (WFA) (Thermo Fisher Scientific) were added to bacterial suspension. The suspension was incubated at room temperature for 1 h with agitation. Sample aliquots were assayed to record the total fluorescence. The incubations were then centrifuged (16,000 g, 3 min), washed with PBS three times to remove unbound lectin, and resuspended in 0.5 mL of PBS. Sample aliquots were assayed to determine the pellet fluorescence. Data are presented as a percentage of the fluorescence of the pellet normalized to the total fluorescence of the sample.

### Fluorescent and differential interference contrast (DIC) microscopy

Fluorescent and differential interference contrast (DIC) microscopy was performed as described previously ^51^. Briefly, exponential phase cells (OD_600_ ∼0.3) were fixed with paraformaldehyde (4% final concentration), pipetted onto poly-L-lysine coated microscope-slide-cover-glasses (high performance, D=0.17 mm, Zeiss), and allowed to settle for one hour at room temperature. Cells were then incubated with FITC-conjugated WFA (15.6 µg mL^-1^) for 15 min at room temperature. The samples were dried at room temperature after four washes with PBS and subsequently mounted on a microscope slide with ProLong Glass Antifade Kit (Invitrogen). Samples were imaged on a Leica SP8 microscope with 100×, 1.44 N.A. objective and DIC optics. Images were processed with Huygens Professional software.

### Isolation and purification of membrane glycoproteins

*S. pyogenes* and *S. mutans* strains were grown in THY and *S. pneumoniae* strains were grown in BHI to final OD_600_ ∼0.8. Bacteria (2 L) were harvested by centrifugation at 18,000 g at 4 °C and stored at −80 °C until processing. Cell pellets were resuspended in 50 mL of lysis buffer containing 20 mM Tris-HCl pH 7.5, 300 mM NaCl, 200 µg mL^-1^ mutanolysin, 250 µg mL^-1^ lysozyme, 1× protease inhibitor cocktail (Sigma-Aldrich) and 2 µg mL^-1^ DNase. To lyse *S. mutans* and *S. pyogenes*, recombinant 50 µg mL^-1^ zoocin A ^108^ and PlyC ^109^ were added to the lysis buffer, respectively. Suspension was homogenized and incubated for 1 h at room temperature. *S. mutans* and *S. pneumoniae* were lysed by bead beating (0.1 mm disruptor beads) followed by incubation of the homogenized samples at room temperature for 30 min on a spin rotator. The cell lysate was centrifuged at 5,000 g for 10 min at 4 °C to remove beads and cell debris. The resulting supernatant was centrifuged at 104,000 g for 1 h at 4 °C. The pellet was solubilized in 50 mL of the membrane-solubilization buffer (20 mM Tris-HCl pH 7.5, 300 mM NaCl, 1% Triton X-100, 1 mM CaCl_2_, 1 mM MgCl_2_, 1 mM MnCl_2_, and 1× bacterial protease inhibitor cocktail). Membrane-associated proteins were separated from cell debris by centrifugation at 104,000 g for 1 h at 4 °C. *S. mutans* glycoproteins were purified on a column prefilled with 2 mL of agarose-bound WFA (Vector Laboratories). *S. pyogenes* and *S. pneumoniae* glycoproteins were purified on a column prefilled with 2 mL of sepharose-bound concanavalin A from *Canavalia ensiformis* (Millipore Sigma). Columns were washed with 10 volumes of the membrane-solubilization buffer and glycoproteins were eluted with 50 mM GalNAc from agarose-bound WFA and 300 mM methyl α-D-mannopyranoside from sepharose-bound concanavalin. The eluent was further concentrated to a final volume of 1 mL using an Amicon Ultra-10 mL (10,000 MWCO) spin filter followed by methanol-chloroform precipitation. The resulting protein pellet was resuspended in the SDS-PAGE loading buffer and separated on a 15% SDS-PAGE gel using Tris-glycine buffer (*S. mutans* proteins in Fig. 4d) or 4-12% SurePAGE™ Bis-Tris gel (Genscript) in MES buffer (all other protein samples) followed by Coomassie blue staining.

### LC-MS/MS analysis of proteins

Proteins were identified by LC–MS/MS proteomics analysis by the Taplin Biological Mass Spectrometric Facility, Harvard Medical School, Boston, MA (Orbitrap mass spectrometers, Thermo Fisher Scientific). Briefly, Coomassie-stained protein bands were excised and subjected to a modified in-gel trypsin digestion procedure ^110^. Peptides were reconstituted in 2.5% acetonitrile plus 0.1% formic acid, loaded *via* a Thermo EASY-LC (Thermo Fisher Scientific) and eluted with 90% acetonitrile plus 0.1% formic acid. Subsequently, peptides were subjected to electrospray ionization and loaded into either an Orbitrap Exploris 480 mass spectrometer (Thermo Fisher Scientific) or a Velos Orbitrap Pro ion-trap mass spectrometer (Thermo Fisher Scientific). Peptide sequences were determined by matching protein databases with the acquired fragmentation pattern by Sequest (Thermo Fisher Scientific). All databases include a reversed version of all the sequences, and the data was filtered to between a one and two percent peptide false discovery rate. The results of the analysis are provided in Supplementary Data 1.

### SDS-PAGE and immunoblot analysis of *S. mutans* and *S. pyogenes* cell lysates

*S. mutans* and *S. pyogenes* grown to an OD_600_ ∼0.5 were pelleted by centrifugation at 3,200 g for 15 min at 4 °C. The cell pellet was resuspended in 1 mL of lysis buffer containing 20 mM Tris-HCl pH 7.5, 100 mM NaCl, 1% Triton X-100, 1% SDS, 0.5% DOC, 1× bacterial protease inhibitor cocktail, and 2 µg mL^-1^ DNase. Cells were disrupted by bead beating for 1 h at room temperature and centrifuged at 16,000 g for 25 min. The total protein content in the supernatant fraction was determined by Pierce™ BCA protein assay kit (Thermo Fisher Scientific). An equivalent amount of total proteins was loaded in each lane and resolved on a 15% SDS-PAGE gel or 4-12% SurePAGE™ Bis-Tris gel (Genscript), transferred to nitrocellulose by electroblotting, and subjected to immunoblot analysis using as primary antibodies either the polyclonal produced in rabbits or anti-FLAG® antibody (Millipore Sigma). Rabbit polyclonal antibodies against the *S. pyogenes* PrsA1 and PrsA2 homologs were provided by Dr. Yung-Chi Chang ^39^. Rabbit polyclonal antibodies against the *S. pyogenes* PknB homolog, and the *S. mutans* PrsA, FtsQ and PknB homologs were produced by Cocalico Biologicals. Purified recombinant PrsAΔIDR, FtsQΔIDR, PknBΔIDR^Smu^, and PknBΔIDR^Spy^ were used to produce the antibodies. All anti-sera except *S. pyogenes* anti-PknB were validated by immunoblot assay using streptococcal strains expressing the WT bacteria and the mutants deficient in the corresponding tested protein. The validation of *S. pyogenes* PrsA1 and PrsA is shown in Fig. 1e and Supplementary Fig. 6a. The validation of *S. mutans* PrsA, FtsQ and PknB is shown in Supplementary Fig 27. *S. pyogenes* anti-PknB antibodies were validated using *S. pyogenes* WT and the *pknB*ΔIDR mutant expressing the IDR-less PknB (Fig. 1f)

### SDS-PAGE and immunoblot analysis of *S. pneumoniae* cell lysates

Cell lysate preparations using SEDS lysis buffer [0.1% deoxycholate (vol/vol), 150 mM NaCl, 0.2% SDS (vol/vol), 15 mM EDTA pH 8.0] and immunoblotting was performed as previously described ^73, 111^. Briefly, bacteria were grown exponentially in 5 ml BHI broth to an OD_620_ ∼ 0.15–0.2. 1.8 mL of culture were centrifuged at 4 °C at 16,100 g for 5 min, and washed once with PBS at 4 °C. Pellets were frozen on dry ice for more than 15 min, then thawed, and resuspended in volumes of SEDS lysis buffer proportional to the harvest OD_620_ values. Frozen pellets collected from 1.8 mL of cultures at OD_620_ ∼0.16 were suspended in 80 μL of SEDS lysis buffer followed by incubation at 37 °C for 15 min with shaking. Proteins were denatured by addition of 2× Laemmeli SDS loading buffer (Bio-Rad) plus β-mercaptoethanol (5% vol:vol) at 95 °C for 10 min. 8 μL of lysate were loaded in each lane and resolved on a 7.5% Mini-PROTEAN® TGX™ precast protein gel (Biorad 4561025) for PBP1a and PBP1b samples, or 10% precast protein gels (Biorad 4561035) for PrsA and StkP samples. For RodZ-L-FLAG^3^ and MapZ samples, 8 or 6 μL respectively, were loaded in each lane of 4–15% Mini-PROTEAN® TGX™ Precast Protein Gels (Biorad 4561085). For quantification, 4, 8, and 12 μL of WT samples were also loaded in additional lanes. The proteins were transferred to nitrocellulose, stained with Totalstain Q-NC reagent from Azure biosystems, and imaged as described by the manufacturer. Signal intensities from Totalstain Q-NC stain were used to normalize antibody signals. Rabbit polyclonal antibodies used were anti-StkP (1:7,000) ^112, 113^, anti-PBP1a^Spn^ (1:7,000) ^73^, anti-PBP1b^Spn^ (Dr. Cécile Morlot, 1:7,000), and anti-PrsA^Spn^ (Dr. Marien de Jonge, 1:7,000), anti-MapZ^Spn^ (Dr. Linda Doubravová), and anti-FLAG (Sigma, F7425, 1:2000). Anti-sera were validated by immunoblot assays using WT and the Δ*pbp1a*, Δ*pbp1b*, Δ*prsA* or Δ*mapZ* mutants (Fig. 3c, d, e, f) or the WT strain with no FLAG-tag (Fig. 3g). The secondary antibody used was Licor IR Dye800 CW goat anti-rabbit (926–32,211, 1:14,000). IR Signal intensities were detected with Azure biosystem 600 and normalized with total protein stain signals for each lane using Totalstain Q-NC reagent. Measurements of band intensities were performed with AzureSpot software.

### Biofilm assay

*S. mutans* cells grown overnight (17 h) were used as an inoculum at a 100-fold dilution in UFTYE (2.5% tryptone and 1.5% yeast extract containing) medium supplemented with 1% (wt/vol) sucrose or glucose. Bacterial biofilms were grown in 24-well plates (Corning) at 37 °C in the presence of 5% CO_2_ for 24 h. For ethanol stress, 3.5% ethanol was added to the growth medium. After 24 h of growth, cell suspension in each well was removed, and biofilm was washed three times with water. The biofilm was stained with crystal violet (0.2%) followed by washes and resuspension in a de-staining solution (10% methanol, 7.5% acetic acid). Finally, the absorbance of the de-staining solution was measured at 540 nm.

### Thermal protein stability assay

The protein thermal stability measurements were performed using Tycho NT.6 (NanoTemper Technologies). The purified protein samples were diluted in the test condition buffers to a range of concentrations from 1 to 2 mg mL^-1^. The samples were heated from 35 to 95 °C at a rate of 30 °C min^-1^. Protein unfolding was monitored by measuring the intrinsic fluorescence of tryptophan and tyrosine residues at emission wavelengths of 330 and 350 nm. The inflection temperature (Ti) was determined by calculating the maximum of the first derivative of the temperature-dependent change in fluorescence ratio F_350_/F_330_. The data were processed by Tycho NT.6 software.

### Circular dichroism (CD)

Purified recombinant proteins ePrsA-FL and ePrsAΔIDR were buffer exchanged into 20 mM sodium phosphate pH 7.4, and 150 mM sodium fluoride. CD spectra were documented at a protein concentration of 0.80 mg/mL over a 200–250 nm wavelength range with a scan rate of 100 nm/min using a JASCO J-810 spectropolarimeter at 25 °C. The secondary structure content of ePrsA-FL and ePrsAΔIDR was calculated using the BeStSel server ^114^.

### Prediction of extracytoplasmic IDRs

Predictions were run and aggregated using a custom Python script we developed. It takes as input and parses a genome in GenBank file format, extracting individual protein sequences for forwarding to the predictors, handles switching conda environments, and includes logic for resuming incomplete runs. After both predictors are run, it concatenates the results as both a residue map and provides a list of every gene that contains predicted transmembrane regions, signal peptides, and disordered regions. Transmembrane regions and signal peptides were predicted using serial runs of the protein language model TMbed on the coding regions of *S. pyogens* MGAS2221 (GenBank: CP043530.1), *S. mutans* Xc (GenBank: CP183775.1), *S. pneumoniae* D39 (GenBank: CP000410.2) and *S. pneumoniae* TIGR4 (https://genomes.atcc.org/genomes/641decb736d446cc) with default batch size ^64^. Disordered region prediction was performed using the compact protein language model DR-BERT using the pretrained weights ^115^. To identify extracytoplasmic IDRs, the resultant subset of membrane-associated and secreted proteins possessing IDRs (Supplementary Data 2, 3 and 4) was analyzed by TOPCONS ^67^. The membrane topology was also validated by analysis of structural models generated by AlphaFold2 prediction pipeline ^61^. Selected proteins were additionally analyzed using protein disorder predictors PONDR® VLS2, PONDR® VL3, PONDR® VLXT, PONDR® FIT, IUPred-Long, and IUPred-Short as implemented in the Rapid Intrinsic Disorder Analysis Online (RIDAO) platform ^75^.

### Statistical analysis

Unless otherwise indicated, statistical analysis was performed using GraphPad Prism version 10 on pooled data from at least three independent biological experiments. A two-tailed Student’s *t*-test and one-way ANOVA with Tukey’s multiple comparisons test were performed as described for individual experiments. A *P*-value equal to or less than 0.05 was considered statistically significant.

## Data availability

All data generated during this study are included in the article and supplementary information files. Source data are provided with this paper.

## Code availability

Code for automated running of TMbed and DR-BERT has been released on GitHub at: https://github.com/ctch225/ScanDis (DOI: 10.24433/CO.9809200.v1).

## Acknowledgments

The authors thank Dr. Yung-Chi Chang (National Taiwan University) for the gift of anti-*S. pyogenes* PrsA1 and PrsA2 antibodies, Dr. Cécile Morlot (Institue de Biologie Structurale, Grenoble) for the gift of anti-*S. pneumoniae* PBP1b antibodies, Dr. Marien de Jonge (Radboud University Medical Center) for the gift of anti-*S. pneumoniae* PrsA antibodies, Dr. Linda Doubravová (Institute of Microbiology, Academy of Sciences of the Czech Republic) for the gift anti-MapZ antibodies, and Dr. Catalina Velez-Ortega (University of Kentucky) for access to the Leica SP8 confocal microscope. We thank William de Souza and Joseph Mougous for their comments on the manuscript.

This work was supported by NIH grants R01DE028916 from the NIDCR, R21AI149366 from the NIAID and the Hypothesis Fund (to NK), and R35GM131767 from the NIGMS (to MEW). The funders had no role in study design, data collection and interpretation, or the decision to submit the work for publication.

## Author contributions

MMR, SZ, CTC, JSR, H-CTT, MEW, KVK, and NK designed the experiments. MMR, SZ, CTC, JSR, CWK, YMH, H-CTT, KVK, and NK performed functional and biochemical experiments. SZ performed microscopy analysis. CTC developed code. VNU conducted bioinformatics analysis. H-CTT, SZ, KVK, and NK constructed streptococci and *E. coli* strains. MMR, SZ, CTC, JSR, H-CTT, MEW, KVK, and NK analyzed the data. NK wrote the manuscript with contributions from all authors. All authors reviewed the results and approved the final version of the manuscript.

