## Supplementary Information for "Glycosylation of serine/threonine-rich intrinsically disordered regions of membrane-associated proteins in streptococci"

**Supplementary Table 1.** Activity of membrane fractions of L-arabinose-induced *E. coli* JW2347 cells, carrying empty expression vector (pBAD33) or vector with $gtrB^{Spy}$ (pBAD33_GtrB$^{Spy}$) in presence of Und-P and UDP-[$^3$H]Glc [a].

| Gene expressed | Activity, pmol/mg |
|---|---|
| None | 1.5 |
| $gtrB^{Spy}$ | 88.9 |

[a] The *E. coli* membrane fraction was prepared as outlined in Methods. Reaction mixtures contained 50 mM HEPES-OH, pH 7.4, 20 mM $MgCl_2$, 5 mM ATP, 1 mM sodium orthovanadate, 1 mM dithiothreitol, 80 μM Und-P (dispersed ultrasonically in 1 % CHAPS), 0.35 % CHAPS (final concentration in the reaction), 2 μM UDP-Glc (500 cpm/pmol) and *E. coli* membrane fraction (100-200 μg protein) in a total volume of 0.025 mL. Following 5 min at 37 °C reactions were stopped by the addition of 2 mL $CHCl_3/CH_3OH$ and the [$^3$H]glucolipid product isolated as described in Methods.

**Supplementary Table 2**. Effect of mild acid and mild alkali on the minor [$^3$H]glucolipid product purified from the membranes of *S. pyogenes* MGAS2221.

| Treatment [a] | Aqueous phase | Organic phase [b] |
|---|---|---|
| Mild acid | 96.6% | 3.4% |
| Mild alkaline methanolysis | 1.8% | 98.2% |

[a] The minor [$^3$H]glucolipid product was prepared from the membranes of *S. pyogenes* MGAS2221 as described in Methods, split into two tubes and dried. For mild acid treatment, the lipid was incubated in 0.2 mL 0.2 N HCl and isopropanol (1:1) for 1 hr at 50 °C. Following neutralization with Tris-OH, 40 vol $CHCl_3/CH_3OH$ (2:1) was added, the reactions were mixed vigorously and centrifuged briefly to separate the two phases. Aqueous and organic phases were collected, dried, and radioactivity determined by scintillation spectrometry in Econosafe counting cocktail. For mild alkaline methanolysis, the lipid was incubated in 0.2 mL toluene and methanol (1:3) containing 0.1 M KOH for 1 hr at 0 °C. Following neutralization with acetic acid, 1 mL $CHCl_3$, 0.35 mL $CH_3OH$ and 0.3 mL 0.9 M NaCl were added and the reactions were centrifuged to separate phases. The two phases were dried and processed as outlined above. The experiment was done twice with the similar results.

[b] Organic phases contain stable [$^3$H]glucolipid product.

Results are presented as a percentage of radioactivity of the sample normalized to the total radioactivity of [$^3$H]glucolipid product before the treatment.

**Supplementary Table 3.** List of membrane-associated and secreted proteins harboring Ser/Thr-rich IDRs.

| Bacteria | Strains[a] | Proteins with Ser/Thr-rich IDRs identified by DR-BERT and AlphaFold2 | Maximum number of Ser/Thr in a stretch | Proteins purified by lectin chromatography | Glycoproteins validated by Immunoblot assay[b] |
|---|---|---|---|---|---|
| *S. pyogenes* | 2221 | PrsA1 | 5 | + | + |
| | | PBP1a | 4 | + | N.D. |
| | | PknB | 12 | + | + |
| | | LytR[c] | 4 | + | N.D. |
| | | MGAS2221_0652 | 8 | - | N.D. |
| *S. pneumoniae* | TIGR4 | PrsA | 8 | + | N.D. |
| | | PBP1a | 6 | + | N.D. |
| | | PBP1b | 11 | - | N.D. |
| | | CdaR | 7 | + | N.D. |
| | | RodZ | 8 | + | N.D. |
| | | MapZ | 9 | + | N.D. |
| | | Spn_2093 | 3 | + | N.D. |
| | | LytC | 4 | - | N.D. |
| | | CbpE | 5 | +/-[d] | N.D. |
| | | SP_1604 | 6 | - | N.D. |
| | D39[e] | PrsA | 8 | + | + |
| | | PBP1a | 6 | + | + |
| | | PBP1b | 11 | + | + |
| | | CdaR | 7 | + | N.D. |
| | | RodZ | 8 | + | + |
| | | MapZ | 9 | + | + |
| | | LytC | 4 | + | N.D. |
| | | CbpE | 5 | - | N.D. |
| | | SPD_RS07595 | 6 | - | N.D. |
| *S. mutans* | Xc | PrsA | 6 | + | + |
| | | PBP1a | 10 | + | + |
| | | PknB | 4 | +/-[d] | + |
| | | FtsQ | 2 | +/-[d] | + |
| | | LytR (BrpA)[c] | 4 | +/-[d] | N.D. |
| | | LtaS | 7 | - | N.D. |
| | | Smu.63c[f] | 6 | - | N.D. |
| | | Smu.503c[f] | 5 | - | N.D. |
| | | Smu.690[f] | 5 | - | N.D. |
| | | Smu.695[f] | 6 | - | N.D. |
| | | PplD | 5 | - | N.D. |
| | | GtfB | 8 | - | N.D. |

[a] Indicates genome used for analysis by deep learning algorithms.

[b] Indicates glycoproteins validated by differences in electrophoretic mobility assessed by immunoblot analysis. N.D indicates not determined.

[c] *S. pyogenes* and *S. mutans* have two LytR-Cps2A-Psr family proteins, LCP1 and LCP2. The LCP1 homolog in *S. pyogenes* is known as LytR. The LCP1 homolog in *S. mutans* is known as BrpA.

[d] LC-MS/MS analysis revealed the presense of small amounts of peptides.

[e] NCBI Reference Sequence: NC_008533.2

[f] Indicates the protein name as assigned by UA159 genome sequencing (GenBank: AE014133.2)

**Supplementary Table 4.** *S. pyogenes* and *S. mutans* strains and plasmids.

| Strain or plasmid | Description [a] | Reference |
|---|---|---|
| *Streptococcus pyogenes* | | |
| MGAS 2221 | *S. pyogenes* M1T1-serotype strain, parent | [1] |
| 2221Δ*gtrB* | *gtrB* deletion mutant has a nonpolar kanamycin resistance cassette inserted in M5005_spy_420 (*gtrB*) in the 2221 strain; Kan$^{R}$ | This study |
| 2221Δ*gtrB*:p*gtrB* | 2221Δ*gtrB* is complemented with p*gtrB* carrying *S. pyogenes gtrB;* Kan$^{R}$, Cam$^{R}$ | This study |
| 2221Δ*gtrB*:p*sccN* | 2221Δ*gtrB* is complemented with p*sccN* carrying *S. mutans sccN* [2]; Kan$^{R}$, Cam$^{R}$ | This study |
| 2221Δ*pgtC1* | *pgtC1* deletion mutant has a nonpolar spectinomycin resistance cassette inserted in M5005_spy_1794 in the 2221 strain; Spec$^{R}$ | This study |
| 2221Δ*pgtC1*:p*pgtC1* | 2221Δ*pgtC1* is complemented with p*pgtC1* carrying *S. pyogenes pgtC1;* Spec$^{R}$, Cam$^{R}$ | This study |
| 2221Δ*pgtC2* | *pgtC2* deletion mutant has a nonpolar kanamycin resistance cassette inserted in M5005_spy_1862 (*pgtC2*) in the 2221 strain; Kan$^{R}$ | This study |
| 2221Δ*pgtC2*:p*pgtC2* | 2221Δ*pgtC2* is complemented with p*pgtC2* carrying *S. pyogenes pgtC2;* Kan$^{R}$, Cam$^{R}$ | This study |
| 2221 *prsA1*ΔIDR | *prsA1* IDR deletion mutant has a nonpolar spectinomycin resistance cassette inserted in the C-terminal IDR region of *prsA1;* Spec$^{R}$ | This study |
| 2221 *pknB*ΔIDR | *pknB* IDR deletion mutant has a nonpolar spectinomycin resistance cassette inserted in the C-terminal IDR region of *pknB;* Spec$^{R}$ | This study |
| 2221Δ*prsA1* | *prsA1* deletion mutant has a nonpolar spectinomycin resistance cassette inserted in *prsA1;* Spec$^{R}$ | This study |
| 2221*prsA2* *Ω*pKV304 | *prsA2* single crossover mutant. The *prsA2* gene interrupted with pHY304 derived plasmid harboring a 700 bp fragment of *prsA2*. | This study |
| *Streptococcus mutans* | | |
| Xc | Serotype *c* strain *S. mutans*, parent | [3] |
| XcΔ*pgfS* | *pgfS* deletion mutant has a nonpolar kanamycin resistance cassette inserted in *pgfS* in the Xc background*;* Kan$^{R}$ | This study |
| XcΔ*pgfS*: p*pgfS* | Δ*pgfS* is complemented with p*pgfS* in the Xc background*;* Spec$^{R}$, Cam$^{R}$ | This study |
| XcΔ*pgfM1* | *pgfM1* deletion mutant has a nonpolar kanamycin resistance cassette inserted in *pgfM1* in the Xc background; Kan$^{R}$ | This study |
| XcΔ*pgfM2* | *pgfM2* deletion mutant has a nonpolar erythromycin resistance cassette inserted in *pgfM2* in the Xc background; Erm$^{R}$ | This study |
| XcΔ*cnaB-cbpA-cnm* | c*nm-cnaB-cbpA* deletion mutant has a nonpolar erythromycin resistance cassette replacing c*nm-cnaB-cbpA* gene region in the Xc background; Erm$^{R}$ | This study |

| | | |
|---|---|---|
| XcΔ*prsA* | *prsA* deletion mutant has a nonpolar kanamycin resistance cassette inserted in *prsA* in the Xc background; $Kan^R$ | This study |
| XcΔ*prsA*:p*prsA* | XcΔ*prsA* is complemented with p*prsA* carrying *S. mutans prsA*; $Kan^R$, $Cam^R$ | This study |
| *prsA*ΔIDR | *prsA* IDR deletion mutant has a nonpolar erythromycin resistance cassette inserted in the C-terminal IDR region of *prsA;* $Erm^R$ | This study |
| *prsA*ΔIDR Δ*pgfS* | *prsA* IDR deletion mutant in the XcΔ*pgfS* background; $Erm^R$, $Spec^R$ | This study |
| *pknB*ΔIDR | *pknB* IDR deletion mutant has a nonpolar erythromycin resistance cassette inserted in the C-terminal IDR region of *pknB* in the Xc strain; $Erm^R$ | This study |
| *brpA*ΔIDR | *brpA* IDR deletion mutant has a nonpolar erythromycin resistance cassette inserted in the C-terminal IDR region of *brpA* in the Xc strain; $Erm^R$ | This study |
| Δ*wapA*Δ*cnaB-cbpA-cnm* | *wapA* deletion mutant in the Δ*cnaB-cbpA-cnm* background (has a nonpolar spectinomycin resistance cassette inserted in *wapA;* $Erm^R$, $Spec^R$ | This study |
| *prsA*-3×FLAG | 3xFLAG tag was fused with the C-terminus of *prsA* creating *prsA*-3XFLAG followed by a nonpolar erythromycin resistance cassette; $Erm^R$ | This study |
| *prsA*-3×FLAG Δ*pgfS* | *prsA*-3XFLAG in the Δ*pgfS* background; $Erm^R$, $Spec^R$ | This study |
| *prsA*ΔIDR-3×FLAG | 3xFLAG tag was fused with *prsA* lacking the IDR creating *prsA*ΔIDR-3XFLAG followed by a nonpolar erythromycin resistance cassette; $Erm^R$ | This study |
| *prsA*ΔIDR-3×FLAG Δ*pgfS* | *prsA*ΔIDR-3XFLAG in the XcΔ*pgfS* background; $Erm^R$, $Spec^R$ | This study |
| *pbp1A*-3×FLAG | 3xFLAG tag was fused with the C-terminus of PBP1a creating *pbp1A*-3xFLAG followed by a nonpolar erythromycin resistance cassette; $Erm^R$ | This study |
| *pbp1A*-3×FLAG Δ*pgfS* | *pbp1A*-3xFLAG in the XcΔ*pgfS* background; $Erm^R$, $Spec^R$ | This study |
| *pbp1A*ΔIDR-3×FLAG | 3xFLAG tag was fused with the C-terminus of PBP1a lacking the IDR followed a nonpolar erythromycin resistance cassette; $Erm^R$ | This study |
| *pbp1A*ΔIDR-3×FLAG Δ*pgfS* | *pbp1A*ΔIDR-3XFLAG in the XcΔ*pgfS* background; $Erm^R$, $Spec^R$ | This study |
| *prsA*-$IDR^{wapA}$ | C-terminal IDR of *prsA* was replaced with the *wapA* IDR (has a nonpolar erythromycin resistance cassette); $Erm^R$ | This study |
| *prsA*-$IDR^{wapA}$ Δ*pgfS* | C-terminal IDR of *prsA* was replaced with the *wapA* IDR in the XcΔ*pgfS* background; $Erm^R$, $Spec^R$ | This study |
| *prsA*-$IDR^{PrsA1\text{-}GAS}$ | C-terminal IDR of *prsA* was replaced with the MGAS *prsA1* IDR in the Xc strain (has a nonpolar erythromycin resistance cassette); $Erm^R$ | This study |
| *prsA*-$IDR^{PrsA1\text{-}GAS}$ Δ*pgfS* | C-terminal IDR of *prsA* was replaced with the MGAS 2221 *prsA1* IDR in the XcΔ*pgfS* background; $Erm^R$, $Spec^R$ | This study |

| | | |
|---|---|---|
| *prsA*-PIN1 | C-terminal IDR of *prsA* was replaced with PIN1 domain (has a nonpolar erythromycin resistance cassette); $Erm^R$ | This study |
| *prsA*-PIN1 Δ*pgfS* | C-terminal IDR of *prsA* was replaced with PIN1 domain in the XcΔ*pgfS* background; $Erm^R$, $Spec^R$ | This study |
| *prsA*-$IDR^{NG}$ | All Ser and Thr residues in the C-terminal IDR of PrsA were replaced with asparagine and glycine residues. The mutant has a nonpolar erythromycin resistance cassette downstream of the *prsA* variant; $Erm^R$ | This study |
| *prsA*-$IDR^{NG}$ Δ*pgfS* | *prsA* -$IDR^{NG}$ was constructed in the XcΔ*pgfS* background; $Erm^R$, $Spec^R$ | This study |
| *prsA*-$IDR^{NG}$-3×FLAG | 3xFLAG tag was fused with the C-terminal $IDR^{NG}$ of PrsA. The mutant has a nonpolar erythromycin resistance cassette; $Erm^R$ | This study |
| *prsA*-$IDR^{NG}$-3×FLAG Δ*pgfS* | *prsA*-$IDR^{NG}$-3XFLAG in the XcΔ*pgfS* background; $Erm^R$, $Spec^R$ | This study |
| *prsA*Δ$IDR^{8aa}$ | C-terminal 8 amino acids of IDR of *prsA* was deleted in the Xc strain (has a nonpolar erythromycin resistance cassette); $Erm^R$ | This study |
| *prsA*Δ$IDR^{23aa}$ | C-terminal 23 amino acids of IDR of *prsA* was deleted in the Xc strain (has a nonpolar erythromycin resistance cassette); $Erm^R$ | This study |
| *prsA*Δ$IDR^{8aa}$ Δ*pgfS* | C-terminal 8 amino acids of IDR (A326-E333) of *prsA* was deleted in the XcΔ*pgfS* background; $Erm^{R;}$ $Spec^R$ | This study |
| *prsA*Δ$IDR^{23aa}$ Δ*pgfS* | C-terminal 23 amino acids of IDR (A311-E-333) of *prsA* was deleted in the XcΔ*pgfS* background; $Erm^R$, $Spec^R$ | This study |
| XcΔ*pknB* | *pknB* deletion mutant has a nonpolar erythromycin resistance cassette inserted in *pknB* in the Xc strain; $Erm^R$ | This study |
| XcΔ*ftsQ* | *ftsQ* deletion mutant has a nonpolar erythromycin resistance cassette inserted in *ftsQ* in the Xc strain; $Erm^R$ | This study |
| XcΔ*pgtC2* | *pgtC2* (*smu.2160*) deletion mutant has a nonpolar erythromycin resistance cassette inserted in *smu.2160* in the Xc strain; $Erm^R$ | This study |
| XcΔ*sccN* | *sccN* deletion mutant has a nonpolar spectinomycin resistance cassette inserted in *sccN*; $Spec^R$ | [2] |
| *Escherichia coli* | | |
| DH5α | *E. coli* cells used for cloning | Invitrogen |
| LOBSTR(DE3)pRARE2 | *E. coli* cells used for protein expression; $Cam^R$ | Kerafast |
| JW2347 | *E. coli* JW2347 strain that has a deletion in the *gtrB* homolog | [4] |
| *Plasmids* | | |
| pDC123 | *E. coli-streptococcus* shuttle vector, JS-3 replicon, chloramphenicol resistance cassette; $Cam^R$ | [5] |
| p*pgfS* | pDC123 derived plasmid expressing *pgfS;* $Cam^R$ | This study |

| | | |
|---|---|---|
| p*gtrB* | pDC123 derived plasmid expressing *gtrB*$^{Spy}$; $Cam^R$ | This study |
| p*pgtC2* | pDC123 derived plasmid expressing *pgtC2*$^{Spy}$; $Cam^R$ | This study |
| p*sccN* | pDC123 derived plasmid expressing *sccN;* $Cam^R$ | [2] |
| p*prsA* | pDC123 derived plasmid expressing *prsA;* $Cam^R$ | This study |
| pRSF-NT | A modified pRSF-Duet1 (Novagen) vector that allows the creation of N-terminus His-tagged proteins with a TEV protease cleavage site; $Kan^R$ | [6] |
| pBAD33_GtrB$^{Spy}$ | pBAD33 derived plasmid expressing GtrB$^{Spy}$; $Amp^R$, $Cam^R$ | This study |
| pRSF-NT-PrsAΔIDR | pRSF-NT derived plasmid for expression of *S. mutans* PrsAΔIDR (PrsA lacking the IDR) fused at the N-terminus with a His-tag followed by a TEV protease recognition site; $Kan^R$ | This study |
| pRSF-NT-FtsQΔIDR | pRSF-NT derived plasmid for expression of *S. mutans* FtsQΔIDR (FtsQ lacking the IDR) fused at the N-terminus with a His-tag followed by a TEV protease recognition site; $Kan^R$ | This study |
| pRSF-NT-PknBΔIDR$^{Smu}$ | pRSF-NT derived plasmid for expression of *S. mutans* PknBΔIDR (PknB lacking the IDR) fused at the N-terminus with a His-tag followed by a TEV protease recognition site; $Kan^R$ | This study |
| pRSF-NT-PrsA | pRSF-NT derived plasmid for expression of *S. mutans* PrsA fused at the N-terminus with a His-tag followed by a TEV protease recognition site; $Kan^R$ | This study |
| pRSF-NT-PknBΔIDR$^{Spy}$ | pRSF-NT derived plasmid for expression of *S. pyogenes* PknBΔIDR (PknB lacking the IDR) fused at the N-terminus with a His-tag followed by a TEV protease recognition site; $Kan^R$ | This study |
| pLR16T | Vector encoding spectinomycin resistance cassette, $Spec^R$ | [7] |
| pOSKAR | Vector encoding kanamycin resistance cassette, $Kan^R$ | [8] |
| pHY304 | Vector encoding erythromycin resistance cassette, $Erm^R$ | [9] |
| pKV304 | pHY304 derived plasmid harboring a 700 bp fragment of *prsA2*. | This study |

[a] Antibiotic resistance markers: $Erm^R$, erythromycin; $Kan^R$, kanamycin; $Spec^R$, spectinomycin; $Cam^R$, chloramphenicol; $Str^R$, streptomycin.

**Supplementary Table 5.** *S. pneumoniae* strains used in this study.

| Strain number | Abbreviated genotype | Genotype (description)[a] |
|---|---|---|
| D39 *rpsL1* Δ*cps* strains | | |
| IU1824 | Δ*cps* WT | D39 *rpsL1* Δ*cps* |
| IU9175 | Δ*cps* Δ*mapZ* | D39 *rpsL1* Δ*cps* Δ*mapZ* |
| IU18448 | Δ*cps* Δ*gtrB*::P$_c$-*erm* | D39 *rpsL1* Δ*cps* Δ*gtrB* (*spd_1431*)::P$_c$-*erm* |
| IU18454 | Δ*cps* Δ*gtrB*::P$_c$-*kan-rpsL*$^+$ | D39 *rpsL1* Δ*cps* Δ*gtrB* (*spd_1431*)::P$_c$-*kan-rpsL*$^+$ |
| IU18456 | Δ*cps* Δ*pgtC2* (Δ*spd_2058*) | D39 *rpsL1* Δ*cps* Δ*pgtC2*::P$_c$-*kan-rpsL*$^+$ |
| IU20268 | Δ*cps* *pbp1a*ΔIDR | D39 *rpsL1* Δ*cps* *pbp1a*(ΔIDR(S669-P719)) |
| IU20272 | Δ*cps* *pbp1b*ΔIDR | D39 *rpsL1* Δ*cps* *pbp1b*(ΔIDR(S794-R821)) |
| IU20229 | Δ*cps* *prsA*ΔIDR | D39 *rpsL1* Δ*cps* *prsA*(ΔIDR(S304-E313)) |
| IU20556 | Δ*cps* Δ*gtrB* complementation with ectopic *gtrB,* Δ*gtrB*//P-*gtrB* | D39 *rpsL1* Δ*cps* Δ*gtrB*::P$_c$-*erm* //Δ*bgaA*::*kan*-P$_{ftsA}$-*gtrB* |
| IU20558 | Δ*cps* *gtrB*$^+$ in both native and ectopic site, *gtrB* overexpression, *gtrB*$^+$ | D39 *rpsL1* Δ*cps* Δ*bgaA*::*kan*-P$_{ftsA}$-*gtrB* |
| IU20663 | Δ*cps* *mapZ*ΔIDR | D39 *rpsL1* Δ*cps* *mapZ*(ΔIDR(T318 to S357)) |
| IU20690 | Δ*cps* *mapZ*ΔIDR Δ*gtrB* | D39 *rpsL1* Δ*cps* *mapZ*(ΔIDR(T318 to S357)) Δ*gtrB*::P$_c$-*erm* |
| IU20915 | Δ*cps* *pbp1a*ΔIDR *pbp1b*ΔIDR | D39 *rpsL1* Δ*cps* *pbp1a*(ΔIDR(S669-P719)) *pbp1b*(ΔIDR(S794-R821)) |
| D39 *rpsL1 cps*$^+$ strains | | |
| IU1781 | *cps*$^+$ WT | D39 *cps*$^+$ *rpsL1* |
| IU17989 | *cps*$^+$ Δ*pbp1b* | D39 *rpsL1 cps*$^+$ Δ*pbp1b*::P$_c$-*erm* |
| IU18498 | *cps*$^+$ Δ*gtrB* | D39 *rpsL1 cps*$^+$ Δ*gtrB* (*spd_1431*)::P$_c$-*erm* |
| IU18715 | *cps*$^+$ *gtrB*$^+$ in both native and ectopic site, *gtrB* overexpression, *gtrB*$^+$ | D39 *rpsL1 cps*$^+$ Δ*bgaA*::*kan*-P$_{ftsA}$-*gtrB* |
| IU18719 | *cps*$^+$ Δ*gtrB* complementation with ectopic *gtrB,* Δ*gtrB*//P-*gtrB* | D39 *rpsL1 cps*$^+$ Δ*gtrB*::P$_c$-*erm* //Δ*bgaA*::*kan*-P$_{ftsA}$-*gtrB* |
| IU20077 | *cps*$^+$ Δ*pbp1a* | D39 *rpsL1 cps*$^+$ Δ*pbp1a*::P$_c$-*erm* |
| IU20079 | *cps*$^+$ Δ*prsA* | D39 *rpsL1 cps*$^+$ Δ*prsA*::P$_c$-*erm* |
| IU20112 | *cps*$^+$ *rodZ*-L-FLAG$^3$ Δ*gtrB* | D39 *rpsL1 cps*$^+$ *rodZ*-L-FLAG$^3$-P$_c$-*erm* Δ*gtrB*::P$_c$-*aad9* |
| IU20118 | *cps*$^+$ *rodZ*-L-FLAG$^3$ Δ*gtrB*//P-*gtrB,* Δ*gtrB* complementation with ectopic *gtrB* | D39 *rpsL1 cps*$^+$ *rodZ*-L-FLAG$^3$-P$_c$-*erm* Δ*gtrB*::P$_c$-*aad9*//Δ*bgaA*::*kan*-P$_{ftsA}$-*gtrB* |
| IU20124 | *cps*$^+$ *rodZ*-L-FLAG$^3$ | D39 *rpsL1 cps*$^+$ *rodZ*-L-FLAG$^3$-P$_c$-*erm* |
| IU20130 | *cps*$^+$ *rodZ*-L-FLAG$^3$ P-*gtrB, gtrB*$^+$ in both native and ectopic site, *gtrB* overexpression, *gtrB*$^+$ | D39 *rpsL1 cps*$^+$ *rodZ*-L-FLAG$^3$-P$_c$-*erm* Δ*bgaA*::*kan*-P$_{ftsA}$-*gtrB* |
| IU20242 | *cps*$^+$ *prsA*ΔIDR | D39 *rpsL1 cps*$^+$ *prsA*(ΔIDR(S304-E313)) |
| IU20269 | *cps*$^+$ *pbp1a*ΔIDR | D39 *rpsL1 cps*$^+$ *pbp1a*(ΔIDR(S669-P719)) |
| IU20273 | *cps*$^+$ *pbp1b*ΔIDR | D39 *rpsL1 cps*$^+$ *pbp1b*(ΔIDR(S794-R821)) |

| | | |
|---|---|---|
| IU20289 | *cps*$^+$ *prsA*ΔIDR Δ*gtrB* | D39 *rpsL1 cps*$^+$ *prsA*(ΔIDR(S304-E313)) Δ*gtrB*::$P_c$-*erm* |
| IU20295 | *cps*$^+$ *pbp1a*ΔIDR Δ*gtrB* | D39 *rpsL1 cps*$^+$ *pbp1a*(ΔIDR(S669-P719)) Δ*gtrB*::$P_c$-*erm* |
| IU20299 | *cps*$^+$ *pbp1b*ΔIDR Δ*gtrB* | D39 *rpsL1 pbp1b*(ΔIDR(S794-R821)) Δ*gtrB*::$P_c$-*erm* |
| TIGR4 strains | | |
| - | TIGR4 | Serotype 4 strain *S. pneumoniae* TIGR4 [10] |
| - | TIGR4Δ*pgtC2* | *pgtC2* deletion mutant has a nonpolar kanamycin resistance cassette inserted in *pgtC2* in the TIGR4 strain; Kan$^R$ |

[a] Antibiotic resistance markers: Erm$^R$, erythromycin; Kan$^R$, kanamycin.

**Supplementary Table 6.** *S. pneumoniae* strains used to construct mutants listed in Table 5.

| Strain number | Genotype (description)[a] | Antibiotic resistance[a] | Reference or source |
|---|---|---|---|
| IU1690 | D39 *cps*$^+$ (D39W) | None | [11, 12] |
| IU1781 | D39 *cps*$^+$ *rpsL1* | Str$^R$ | [11] |
| IU1824 | D39 *rpsL1* Δ*cps2A'-cps2H'* = D39 *rpsL1* Δ*cps* | Str$^R$ | [11] |
| IU1945 | D39 Δ*cps2A'-cps2H'*= D39 Δ*cps* | None | [11] |
| E177 | D39 Δ*cps* Δ*pbp1a*::P$_c$-*erm* | Erm$^R$ | [13] |
| E193 | D39 Δ*cps* Δ*pbp1b*::P$_c$-*erm* | Erm$^R$ | [13] |
| E221 | D39 Δ*cps* Δ*prsA*(*spd_0868*)::P$_c$-*erm* (IU1945 X fusion Δ*prsA*::P$_c$-*erm*) | Erm$^R$ | This study |
| K164 | D39 Δ*cps* Δ*pbp1a::*P$_c$-*kan-rpsL*$^+$ | Kan$^R$ | [14] |
| K180 | D39 Δ*cps* Δ*pbp1b::*P$_c$-*kan-rpsL*$^+$ | Kan$^R$ | [15] |
| K213 | D39 Δ*cps* Δ*prsA*(*spd_0868*)::P$_c$-*kan-rpsL*$^+$ (IU1945 X fusion Δ*prsA*::P$_c$-*kan-rpsL*$^+$) | Kan$^R$ | This study |
| IU6291 | D39 Δ*cps rodZ*-L-FLAG$^3$-P$_c$-*erm* | Erm$^R$ | [16] |
| IU6987 | D39 Δcps Δ*rodZ*::P$_c$-*aad9* | | [16] |
| IU9086 | D39 *Δcps rpsL* Δ*mapZ*::P$_c$-*kan-rpsL*$^+$ | Kan$^R$ | [17] |
| IU9175 | D39 *Δcps rpsL1* Δ*mapZ* | Str$^R$ | [17] |
| IU9621 | D39 *Δcps rpsL1* Δ*khpA*//Δ*bgaA*::*kan*-t1t2-P$_{ftsA}$-*khpA*$^+$ | Kan$^R$ | [18] |
| IU17989 | D39 *rpsL1 cps*$^+$ Δ*pbp1b*::P$_c$-*erm* (IU1781 X Δ*pbp1b*::P$_c$-*erm* amplicon from E193) | Str$^R$ Erm$^R$ | This study |
| IU18448 | D39 Δ*cps rpsL1* Δ*gtrB* (*spd_1431*)::P$_c$-*erm* (IU1824 X fusion Δ*gtrB*::P$_c$-*erm* amplicon) | Str$^R$ Erm$^R$ | This study |
| IU18454 | D39 Δ*cps rpsL1*Δ*gtrB* (*spd_1431*)::P$_c$-*kan-rpsL*$^+$ (IU1824 X fusion Δ*gtrB*::P$_c$-*erm* amplicon) | Kan$^R$ | This study |
| IU18456 | D39 Δ*cps rpsL1* Δ*pgtC2* (Δ*spd_2058*)::P$_c$-*kan-rpsL*$^+$ (IU1824 X fusion Δ*spd_2058*::P$_c$-*kan-rpsL*$^+$ amplicon) | Kan$^R$ | This study |
| IU18498 | D39 *rpsL1 cps*$^+$Δ*gtrB* (*spd_1431*)::P$_c$-*erm* (IU1781 X Δ*gtrB*::P$_c$-*erm* amplicon from IU18448) | Str$^R$ Erm$^R$ | This study |
| IU18715 | D39 *rpsL1 cps*$^+$ Δ*bgaA*::*kan*-P$_{ftsA}$-*gtrB* (IU1781 X fusion Δ*bgaA*::*kan*-P$_{ftsA}$-*gtrB* amplicon) | Str$^R$ Kan$^R$ | This study |
| IU18719 | D39 *rpsL1 cps*$^+$ Δ*gtrB*::P$_c$-*erm* //Δ*bgaA*::*kan*-P$_{ftsA}$-*gtrB* (IU18498 X fusion Δ*bgaA*::*kan*-P$_{ftsA}$-*gtrB* amplicon) | Str$^R$ Erm$^R$ Kan$^R$ | This study |
| IU20077 | D39 *rpsL1 cps*$^+$ Δ*pbp1a*::P$_c$-*erm* (IU1781 X Δ*pbp1a*::P$_c$-*erm* amplicon from E177) | Str$^R$ Erm$^R$ | This study |
| IU20079 | D39 *rpsL1 cps*$^+$ Δ*prsA*::P$_c$-*erm* (IU1781 X Δ*prsA*::P$_c$-*erm* amplicon from E221) | Str$^R$ Erm$^R$ | This study |
| IU20087 | D39 *rpsL1 cps*$^+$ Δ*gtrB*::P$_c$-*aad9* (IU1781 X fusion Δ*gtrB*::P$_c$-*aad9* amplicon) | Str$^R$ Spec$^R$ | This study |
| IU20089 | D39 *rpsL1 cps*$^+$ Δ*gtrB*::P$_c$-*aad9//*Δ*bgaA*::*kan*-P$_{ftsA}$-*gtrB* (IU18715 X fusion Δ*gtrB*::P$_c$-*aad9* amplicon) | Str$^R$ Kan$^R$ Spec$^R$ | This study |
| IU20112 | D39 *rpsL1 cps*$^+$ *rodZ*-L-FLAG$^3$-P$_c$-*erm* Δ*gtrB*::P$_c$-*aad9* (IU20087 X *rodZ*-L-FLAG$^3$-P$_c$-*erm* from IU6291) | Str$^R$ Erm$^R$ Spec$^R$ | This study |

| | | | |
|---|---|---|---|
| IU20118 | D39 *rpsL1 cps*$^+$ *rodZ*-L-FLAG$^3$-$P_c$-*erm* Δ*gtrB*::$P_c$-*aad9*//Δ*bgaA*::*kan*-$P_{ftsA}$-*gtrB* (IU20089 X *rodZ*-L-FLAG$^3$-$P_c$-*erm* from IU6291) | $Str^R$ $Erm^R$ $Spec^R$ $Kan^R$ | This study |
| IU20124 | D39 *rpsL1 cps*$^+$ *rodZ*-L-FLAG$^3$-$P_c$-*erm* (IU1781 X *rodZ*-L-FLAG$^3$-$P_c$-*erm* from IU6291) | $Str^R$ $Erm^R$ | This study |
| IU20130 | D39 *rpsL1 cps*$^+$ *rodZ*-L-FLAG$^3$-$P_c$-*erm* Δ*bgaA*::*kan*-$P_{ftsA}$-*gtrB* (IU18715 X *rodZ*-L-FLAG$^3$-$P_c$-*erm* from IU6291) | $Str^R$ $Erm^R$ $Kan^R$ | This study |
| IU20199 | D39 *rpsL1* Δ*cps* Δ*pbp1a::*$P_c$-*kan-rpsL*$^+$ (IU1824 X Δ*pbp1a::*$P_c$-*kan-rpsL*$^+$ from K164) | $Kan^R$ | This study |
| IU20205 | D39 *rpsL1 cps*$^+$ Δ*pbp1a::*$P_c$-*kan-rpsL*$^+$ (IU1781 X Δ*pbp1a::*$P_c$-*kan-rpsL*$^+$ amplicon from K164) | $Kan^R$ | This study |
| IU20207 | D39 *rpsL1 cps*$^+$ Δ*pbp1b::*$P_c$-*kan-rpsL*$^+$ (IU1781 X Δ*pbp1b::*$P_c$-*kan-rpsL*$^+$ amplicon from K180) | $Kan^R$ | This study |
| IU20209 | D39 *rpsL1 cps*$^+$ Δ*prsA::*$P_c$-*kan-rpsL*$^+$ (IU1781 X Δ*prsA::*$P_c$-*kan-rpsL*$^+$ amplicon from K213) | $Kan^R$ | This study |
| IU20242 | D39 *rpsL1 cps*$^+$ *prsA*ΔIDR(S304-E313) (IU20209 X fusion *prsA*ΔIDR(S304-E313) amplicon) | $Str^R$ | This study |
| IU20268 | D39 *rpsL1* Δ*cps pbp1a*(ΔIDR(S669-P719))(IU20199 X fusion *pbp1a*ΔIDR(S669-P719) amplicon) | $Str^R$ | This study |
| IU20269 | D39 *rpsL1 cps*$^+$ *pbp1a*ΔIDR(S669-P719) (IU20205 X fusion *pbp1a*ΔIDR(S669-P719) amplicon) | $Str^R$ | This study |
| IU20273 | D39 *rpsL1 cps*$^+$ *pbp1b*ΔIDR(S794-R821) (IU20207 X fusion *pbp1b*ΔIDR(S794-R821) amplicon) | $Str^R$ | This study |
| IU20289 | D39 *rpsL1 cps*$^+$ *prsA*ΔIDR(S304-E313) Δ*gtrB*::$P_c$-*erm* (IU20242 X Δ*gtrB*::$P_c$-*erm* from IU18448) | $Str^R$ $Erm^R$ | This study |
| IU20295 | D39 *rpsL1 cps*$^+$ *pbp1a*ΔIDR(S669-P719) Δ*gtrB*::$P_c$-*erm* (IU20269 X Δ*gtrB*::$P_c$-*erm* from IU18448) | $Str^R$ $Erm^R$ | This study |
| IU20299 | D39 *rpsL1 cps*$^+$ *pbp1b*ΔIDR(S794-R821) Δ*gtrB*::$P_c$-*erm* (IU20273 X Δ*gtrB*::$P_c$-*erm* from IU18448) | $Str^R$ $Erm^R$ | This study |
| IU20556 | D39 *rpsL1* Δ*cps* Δ*gtrB*::$P_c$-*erm*//Δ*bgaA*::*kan*-$P_{ftsA}$-*gtrB* (IU18448 X Δ*bgaA*::*kan*-$P_{ftsA}$-*gtrB* from IU18715) | $Str^R$ $Erm^R$ $Kan^R$ | This study |
| IU20558 | D39 *rpsL1* Δ*cps* Δ*bgaA*::*kan*-$P_{ftsA}$-*gtrB* (IU1824 X Δ*bgaA*::*kan*-$P_{ftsA}$-*gtrB* from IU18715) | $Str^R$ $Kan^R$ | This study |
| IU20663 | D39 *rpsL1* Δ*cps mapZ*(ΔIDR(T318 to S357)) (IU9086 X fusion *mapZ*(ΔIDR(T318 to S357)) | $Str^R$ | This study |
| IU20690 | D39 *rpsL1* Δ*cps mapZ*(ΔIDR(T318 to S357)) Δ*gtrB*::$P_c$-*erm* (IU20663 X Δ*gtrB*::$P_c$-*erm* from IU18448) | $Str^R$ $Erm^R$ | This study |
| IU20896 | D39 *rpsL1* Δ*cps pbp1a*(ΔIDR(S669-P719)) Δ*pbp1b::*$P_c$-*kan-rpsL*$^+$ (IU20268 X Δ*pbp1b::*$P_c$-*kan-rpsL*$^+$ amplicon from K180) | $Kan^R$ | This study |

| | | | |
|---|---|---|---|
| IU20915 | D39 *rpsL1* Δ*cps* *pbp1a*(ΔIDR(S669-P719)) *pbp1b*(ΔIDR(S794-R821)) (IU20896 X *pbp1b* ΔIDR amplicon) from IU20273) | $Str^R$ | This study |

[a] Antibiotic resistance markers: $Erm^R$, erythromycin; $Kan^R$, kanamycin; $Str^R$, streptomycin; $Spec^R$, spectinomycin.

**Supplementary Table 7.** Primers used to construct *S. pyogenes* and *S. mutans* strains.

| Primer | Sequence[a,b] | Genetic manipulations |
|---|---|---|
| GtrB-BflII-F | GCGTAAGATCTGAAGGCTTTCATTATGATAAGG | Construction of 2221Δ*gtrB* |
| kan-GtrB-R1 | **AGTATTTAAAGATACC**CTTCATTAAAGCAGGGGACAATG | |
| GtrB-kan-F1 | CCTGCTTTAATGAAG**GGTATCTTTAAATACTGTAG** | |
| kan-GtrB-F2 | **TGAATTGTTTTAGTAC**GGATAACACCTTGGGTATTTTG | |
| GtrB-kan-R2 | ACCCAAGGTGTTATCC**GTACTAAAACAATTCATCCAG** | |
| GtrB-XhoI-R | CGTCTCTCGAGGCCTTTTTTTAGTTTCAAGG | |
| Spy1794-BamH-F | CGTCTGGATCCGAGACGATAGAACTGGCCAAC | Construction of 2221Δ*pgtC1* |
| Spec-pgtC1-R1 | **CTCACTATTTTGGTCGAC**GGATTTAGCACTTGTAAGTTC | |
| pgtC1-spec-F1 | TTACAAGTGCTAAATCC**GTCGACCAAAATAGTGAGGAG** | |
| Spec-pgtC1-F2 | **AAAATTATAAGGATCC**GCTTTTTGGACTGGCTATTC | |
| pgtC1-spec-R2 | AGCCAGTCCAAAAAGC**GGATCCTTATAATTTTTTTAATCTG** | |
| pgtC1-XhoI-R | GCGCGCCTCGAGTCGTTATCTGTTTATCTAC | |
| GtrB-EcoRI-F | GCTTCGAATTCGCTTTTGTGTTACAATTGCCC | Construction of p*gtrB* |
| GtrB-BglII-R | GCGTAAGATCTCATTAGTTATCTCCTTTCTGATG | |
| GtrB-Check-F | GAAATATCTCAGCTTTTGTG | Verification of 2221Δ*gtrB* |
| GtrB-Check-R | CATCACAGTAATGTAATTGCC | |
| PgtC1-check-f | CAACTGCTAGCTTCGATTTTTG | Verification of 2221Δ*pgtC1* |
| PgtC1-check-r | CATGACAAATAGCCACTTGAC | |
| GtrB-XbaI-F | GCGACTCTAGAGTGTTACAATTGCCCTAAAATTCC | Construction of pBAD33_GtrB$^{Spy}$ |
| GtrB-HindIII-R | CGCGCAAGCTTGGAGTAGGAAACGATTGATGAG | |
| pgtC2-BamHI-f | CGTCTGGATCCGATAATTGCTGGACTTGCTAG | Construction of 2221Δ*pgtC2* |
| Kan-pgtC2-r1 | **CAGTATTTAAAGATACC**GTAGGAAGTATTACTAGAGCAC | |
| pgtC2-Kan-f1 | CTAGTAATACTTCCTAC**GGTATCTTTAAATACTGTAG** | |
| Kan-pgtC2-f2 | **TGAATTGTTTTAGTAC**CAAGTTTAATGCTCTGCCTATG | |
| pgtC2-Kan-r2 | GCAGAGCATTAAACTTG**GTACTAAAACAATTCATCCAG** | |
| spgtC2-XhoI-r2 | GCGCGCTCGAGAACTAGATAGTCTATTTCTTACAG | |
| pgtC2-check-f | GCTTTTGTTAGGTTTACAGC | Verification of 2221Δ*pgtC2* |
| pgtC2-check-r | GGGAAATAATCCTACGTAAATC | |
| pgtC2-EcoRI-F | GCTTCGAATTCGAAGATCTCTTCCTAGCCTC | Construction of p*pgtC2* |
| pgtC2-BamHI-r | CGTCTGGATCCGGACTAAAGTAATTTAGAC | |
| pgfC-F | GGCGAAGAAAGCGGTATTG | Construction of XcΔ*pgfS* |
| Spec-pgfC-R1 | **CTCACTATTTTGGTCG**AAAAAGGTACTCTGTCTTGATTCC | |
| pgfC-Spec-F1 | CAGAGTACCTTTTT**CGACCAAAATAGTGAGGAG** | |
| pgfC-Spec-R2 | ACCAAACTTACTCTCACC**GATCCTTATAATTTTTTTAATCTG** | |
| Spec-pgfC -F2 | **AAAATTATAAGGATC**GGTGAGAGTAAGTTTGGTTTCAG | |
| pgfC-R | CCATAGCGCAGCCAAATTG | |
| pgfC-XbaI-f | GCGAGCTCTAGAGTGTTAGGCTGTTTTTTGTTC | Construction of p*pgfS* |
| pgfC-BamHI-R | CGTCTGGATCCGTTGCGTGAGTCATCCGAC | |
| pgfC_chF | GGTCAGATTTATCAGTTTCAGTCGG | Verification of XcΔ*pgfS* |
| pgfC_chR | TCGAAATCAGCTGTACACCG | |
| pgfM1-f | GCAACTGGTGATGCTGTTATG | Construction of XcΔ*pgfM1* |
| pgfM1-kan-f1 | TTTAGTGGGATGAGAC**GGTATCTTTAAATACTGTAG** | |
| Kan-pgfM1 -r1 | **CAGTATTTAAAGATACC**GTCTCATCCCACTAAATCCTC | |

| | | |
|---|---|---|
| Kan-pgfM1 -f2 | **TGAATTGTTTTAGTAC**TTGCTTCCAATGATGATTATG | |
| pgfM1-kan-r2 | TCATCATTGGAAGCAA**GTACTAAAACAATTCATCCAG** | |
| pgfM1-r | GGAAGAACACTGAAGTAATTGC | |
| pgfM1-check-f | CGGCCGCATTATATTGTAG | Verification of Xc∆*pgfM1* |
| pgfM1-check-r | CTTCCAAAAGGAAACGTATC | |
| pgfM2-F | CATTGGTGCTCACAGTTCAG | Construction of Xc∆*pgfM2* |
| Er-pgfM2-R1 | **AATTTAACTTCAATTCC**TGCCTCATATCTTGAGTTC | |
| pgfM2-Er-F1 | TCAAGATATGAGGCA**GGAATTGAAGTTAAATTAGATG** | |
| Er-pgfM2 -F2 | **GGAAATAATTCTATG**TGGTGTGATGCCATCTG | |
| pgfM2-Er-R2 | GATGGCATCACACCA**CATAGAATTATTTCCTCCCG** | |
| pgfM2-R | CCAACATAAAAATTAGGCAGAGC | |
| pgfM2-check-F | GTGATGGTATCAAGAGAAGAAAAAG | Verification of Xc∆*pgfM2* |
| pgfM2-check-R | GCTAGACTAAAGGGACTGAGAC | |
| pgfM3-F | CGTGCACTTATGGGAGATATTG | Construction of Xc∆*pgtC2* |
| Er-pgfM3 -R1 | **AATTTAACTTCAATTCC**TTGGTAGCAGAAAACTAAGACTATAG | |
| pgfM3-Er-F1 | GTTTTCTGCTACCAA**GGAATTGAAGTTAAATTAGATG** | |
| Er-pgfM3 -F2 | **GGAAATAATTCTATG**ATGCTGTCAACATCTTATGC | |
| pgfM3-Er-R2 | GATGTTGACAGCAT**CATAGAATTATTTCCTCCCG** | |
| pgfM3-R | CATATTAGGTGAATGCATTCCTTG | |
| pgfM3-check-F | GAAGAAGTTCAGGTTAAAGTAGCAC | Verification of Xc∆*pgtC2* |
| pgfM3-check-F | GTGGTAGCAGGATAAAAACATCTAA | |
| PrsA-F | CGTCCTGTTATTAAGGAAGAC | Construction of Xc∆*prsA* |
| Kan-PrsA-R1 | **GTATTTAAAGATACC**AAACCAATCTCTTTGCGG | |
| PrsA-Kan-F1 | AAAGAGATTGGTTT**GGTATCTTTAAATACTGTAG** | |
| Kan-PrsA-F2 | **TGAATTGTTTTAGTAC**AGCAGAGACAACAGCAG | |
| PrsA-Kan-R2 | CTGTTGTCTCTGCT**GTACTAAAACAATTCATCCAG** | |
| PrsA-R | GACATAGTCAATAGCAATGAC | |
| PrsA-check-F | GATGGTCTGCTGCTGATTCG | Verification of Xc∆*prsA* |
| PrsA-check-R | CATTCGTCAATCTTTACAAG | |
| PrsA-XbaI-F | GCGAG<u>CTCTAGA</u>TTGGTTTGATTGATTAAGTC | Construction of p*prsA* |
| PrsA-BamHI-R | GTCT<u>GGATCC</u>ATTCGTCAATCTTTACAAGTC | |
| Cnm-F | GAATAGCGATAGCGATGGC | Construction of Xc∆*cnaB-cbpA-cnm* |
| Er-Cnm-R1 | **AATTTAACTTCAATTCCGATAGGAAGAGAAGAGCAATTATCTG** | |
| Cnm-Er-F1 | CTCTTCTCTTCCTATC**GGAATTGAAGTTAAATTAGATG** | |
| Er-Cnm-F2 | **GGAAATAATTCTATG**CCATTAAGCTGGAGGTTCAG | |
| Er-Cnm-R2 | CTCCAGCTTAATGG**CATAGAATTATTTCCTCCCG** | |
| Cnm-R | CCTTACCATCAACAGCATCTTTAC | |
| Cnm-check-F | GAATTCTTAATTTCTAATGTAAATGTGAAAGG | Verification of Xc∆*cnaB-cbpA-cnm* |
| Cnm-check-R | CAAGCCAGTAATACTGTCATTGAAAG | |
| WapA-F | GCAGCTTTTGATGCGGCTC | Construction of Xc∆*wapA* |
| Spec-WapA-R1 | **CTCACTATTTTGGTCG**CTTAATAGTTTATGTTTC | |
| WapA-Spec-F1 | CATAAACTATTAAG**CGACCAAAATAGTGAGGAG** | |
| Spec-WapA-F2 | **AAAATTATAAGGATCGTTAACATTGATTATGC** | |
| WapA-Spec-R2 | TAATCAATGTTAAC**GATCCTTATAATTTTTTTAATCTG** | |
| WapA-R | CCACATCAATCCTTCAGC | |
| WapA-check-F | CTACAGATTAACTAAGGAG | |

| | | |
|---|---|---|
| WapA-check-R | GCTTCGATTCCAACCTCG | Verification of Xc∆*wapA* |
| BrpA-F | GACAACCATGACCAGTTTAG | Construction of *brpA*∆IDR |
| BrpA_trunk-Er-F1 | ATACGAGTATTTAA**GGAATTGAAGTTAAATTAGATG** | |
| Er-BrpA_trunk-R1 | **TTTAACTTCAATTCC**TTAAATACTCGTATCGCCTCC | |
| Er-BrpA_trunk-F2 | **GAAATAATTCTATG**CTGCAGGAACGGGAATTGG | |
| BrpA_trunk-Er-R2 | TCCCGTTCCTGCAG**CATAGAATTATTTCCTCCCG** | |
| BrpA-R | CAATCTCTTTGCACGCATTC | |
| BrpA-check-F | CAACGGCTAGTCTTTATGAG | Verification of *brpA*∆IDR |
| BrpA-check-R | GGGTATAACTGAAAAACC | |
| PknB-F | GATATTGGACAGAAGCGTTC | Construction of Xc∆*pknB* |
| Er-PknB -R1 | **TTTAACTTCAATTCC**AATCTGAATCATTGATCTC | |
| PknB-ER-F1 | CAATGATTCAGATT**GGAATTGAAGTTAAATTAGATG** | |
| ER-PknB-F2 | **GAAATAATTCTATG**CAAGTGCGACGACAACTTCAC | |
| PknB-ER-R2 | TGTCGTCGCACTTG**CATAGAATTATTTCCTCCCG** | |
| PknB-R | GAGCAAAATGATTAATCATAAC | |
| PknB-check-F | GTAGCCTTAATTCACGCCG | Verification of Xc∆*pknB* |
| PknB-check-R | CTATATCATTCAATAGTTCG | |
| FtsQ-F | CAATGGGACTGGCTAACCG | Construction of Xc∆*ftsQ* |
| Er-FtsQ-R1 | **TTTAACTTCAATTCC**GCTGCCATTCTGTCAAAG | |
| FtsQ-Er-F1 | GACAGAATGGCAGC**GGAATTGAAGTTAAATTAGATG** | |
| Er-FtsQ-F2 | **GAAATAATTCTATGTAACGCAGGCAACCGTATC** | |
| FtsQ-Er-R2 | GGTTGCCTGCGTTA**CATAGAATTATTTCCTCCCG** | |
| FtsQ-R | CAAAACTGACTTGGTAAGAGC | |
| FtsQ-check-F | CTTTTGTTAACAGATATAACATC | Verification of Xc∆*ftsQ* |
| FtsQ-check-R | GAAACTTAATTTCCTATCTC | |
| PknB_Smu-F | GCCTAAAACCAAACCACAACC | Construction of *pknB*∆IDR |
| Er-PknB_Smu-NoIDR-R1 | **TTTAACTTCAATTCC**TTAATGAGAGCCACCTGATGTTG | |
| PknB_Smu-NoIDR-Er-F1 | GTGGCTCTCATTAA**GGAATTGAAGTTAAATTAGATG** | |
| Er-PknB_Smu-F2 | **GAAATAATTCTATG**TTCAAGTGCGACGACAAC | |
| PknB_Smu-Er-R2 | TCGTCGCACTTGAA**CATAGAATTATTTCCTCCCG** | |
| PknB_Smu-R | CTTGCAAGGTTAATAAAGCG | |
| PknB_Smu-check-F | CATCTAATGATGATATTATC | Verification of *pknB*∆IDR |
| PknB_Smu-check-R | CTATATCATTCAATAGTTCG | |
| PrsA-F | CGTCCTGTTATTAAGGAAGAC | Construction of *prsA*∆IDR |
| PrsAnoIDR-Er-F1 | CAGCAAGCGGATAA**GGAATTGAAGTTAAATTAGATG** | |
| Er-PrsAnoIDR-R1 | **TTTAACTTCAATTCC**TTATCCGCTTGCTGCCGCATATTG | |

| | | |
|---|---|---|
| Er-PrsAnoIDR-F2 | **GAAATAATTCTATG**GACTTGTAAAGATTGACG | |
| PrsAnoIDR-ER-R2 | AATCTTTACAAGTC**CATAGAATTATTTCCTCCCG** | |
| PrsA-R | GACATAGTCAATAGCAATGAC | |
| PrsAnoIDR-check-F | AAACGTCTGAAAGAAATTATCG | Verification of *prsA*$\Delta$IDR |
| PrsAnoIDR-ER-R2 | AATCTTTACAAGTC**CATAGAATTATTTCCTCCCG** | |
| PknB-GAS-NdeI-F | CATTTCATATGACTAAACCAACTTCTGTGAAAG | Construction of pRSF-NT-PknB$\Delta$IDR$^{Spy}$ |
| PknB-GAS-XhoI-R | GCGCGCTCGAGTTAGTGTGTTTCTTCTGGATAAAG | |
| FtsQ-NcoI-F | GAGTCCATGGCTAGTCCTTTGAGTCGTCAAAAG | Construction of pRSF-NT-FtsQ$\Delta$IDR |
| FtsQ-XhoI-R | CAGTCTCGAGTTATTGAGATACGGTTGCC | |
| PknB_Smu-NcoI-F | AGACCATGGCAACAAAACCGAGTACTGTTTC | Construction of pRSF-NT-PknB$\Delta$IDR$^{Smu}$ |
| PknB_Smu-XhoI-R | CTCTCTCGAGTTAATGAGAGCCACCTGATG | |
| PrsA_Smu-NdeI-F | CGCGTCATATGACAAACCAAAACAGTAAAATTGC | Construction of pRSF-NT-PrsA$\Delta$IDR |
| PrsA_Smu-XhoI-R | GCGCGCTCGAGTTATGACCCTGAAGAGCCACTTC | |
| PrsA_Smu-NdeI-F | CGCGTCATATGACAAACCAAAACAGTAAAATTGC | Construction of pRSF-NT-PrsA |
| PrsA-FL_Smu-XhoI-R | GCGCGCTCGAGCAAGTCCTTTATTCTGCTGCTG | |
| PrsA-F | CGTCCTGTTATTAAGGAAGAC | Construction of *prsA*-3$\times$FLAG |
| PrsAflag-Er-F1 | AAGATTATAAAGATGATGATGATAAAGATTATAAAGATGATG ATGATAAATAGGGAATTGAAGTTAAATTAGATG | |
| Er-PrsAflag-R1 | ATCTTTATCATCATCATCTTTATAATCTTTATCATCATCATCTT TATAATCGCCAGATTCCGCTGCTGTTGTCTC | |
| Er-PrsAnoIDR-F2 | **GAAATAATTCTATG**GACTTGTAAAGATTGACG | |
| PrsAnoIDR-ER-R2 | AATCTTTACAAGTC**CATAGAATTATTTCCTCCCG** | |
| PrsA-R | GACATAGTCAATAGCAATGAC | |
| PrsAflag-check-F | CACAATATGCAGCAGCAAG | Verification of *prsA*-3$\times$FLAG |
| PrsAnoIDR-ER-R2 | AATCTTTACAAGTC**CATAGAATTATTTCCTCCCG** | |
| PrsA-F | CGTCCTGTTATTAAGGAAGAC | Construction of *prsA*$\Delta$IDR-3$\times$FLAG |
| PrsAflag-Er-F1 | AAGATTATAAAGATGATGATGATAAAGATTATAAAGATGATG ATGATAAATAGGGAATTGAAGTTAAATTAGATG | |
| PrsAnoIDRflag-R1 | ATCTTTATCATCATCATCTTTATAATCTTTATCATCATCATCTT TATAATCGCCAGATGCTGCTGCATATTGTGATAAG | |
| Er-PrsAnoIDR-F2 | **GAAATAATTCTATG**GACTTGTAAAGATTGACG | |

| | | |
|---|---|---|
| PrsAnoIDR-ER-R2 | AATCTTTACAAGTC**CATAGAATTATTTCCTCCCG** | |
| PrsA-R | GACATAGTCAATAGCAATGAC | |
| PrsAnoIDR-check-F | AAACGTCTGAAAGAAATTATCG | Verification of *prsA*ΔIDR-3×FLAG |
| PrsAnoIDR-ER-R2 | AATCTTTACAAGTC**CATAGAATTATTTCCTCCCG** | |
| Pbp1a-flag-F | CAGATGCTAAATGACTCGGTC | Construction of *pbp1A*-3×FLAG |
| Pbp1a-flag-Er-F1 | ATTATAAAGATCATGATATCGATTACAAGGATGACGATGACAAGTAA**GGAATTGAAGTTAAATTAGATG** | |
| Er-Flag-Pbp1a-R1 | GTAATCGATATCATGATCTTTATAATCACCGTCATGGTCTTTGTAGTCTGGATTAGAAGATGTCGTAGC | |
| Er-Pbp1a-F2 | **GAAATAATTCTATG**CTATTTTAAAGGCACACTCTTG | |
| Pbp1a-Er-R2 | TGCCTTTAAAATAG**CATAGAATTATTTCCTCCCG** | |
| Pbp1a-R | GTGATAAAGATAATCATTATC | |
| Pbp1a-flag-F | CAGATGCTAAATGACTCGGTC | Construction of *pbp1A*ΔIDR-3×FLAG |
| Pbp1a-Flag-Er-F1 | ATTATAAAGATCATGATATCGATTACAAGGATGACGATGACAAGTAA**GGAATTGAAGTTAAATTAGATG** | |
| Er-Flag-Pbp1anoIDR-R1 | GTAATCGATATCATGATCTTTATAATCACCGTCATGGTCTTTGTAGTCACTATTTGACCCACTTGAACC | |
| Er-Pbp1a-F2 | **GAAATAATTCTATG**CTATTTTAAAGGCACACTCTTG | |
| Pbp1a-Er-R2 | TGCCTTTAAAATAG**CATAGAATTATTTCCTCCCG** | |
| Pbp1a-R | GTGATAAAGATAATCATTATC | |
| Pbp1a-flag-chF | GATTCCAACAATAACGATC | Verification of *pbp1A*-3×FLAG |
| Pbp1a-Er-R2 | TGCCTTTAAAATAG**CATAGAATTATTTCCTCCCG** | |
| Pbp1anoIDR-flag-chF | CTGATTGGACTATGCCTGAC | Verification of *pbp1A*ΔIDR-3×FLAG |
| Pbp1a-Er-R2 | TGCCTTTAAAATAG**CATAGAATTATTTCCTCCCG** | |
| PrsA-F | CGTCCTGTTATTAAGGAAGAC | Construction of *prsA*Δ$IDR^{8aa}$ |
| PrsAstop11-Er-F1 | CGACGACTACAACATAA**GGAATTGAAGTTAAATTAGATG** | |
| Er-PrsAstop11-R1 | **TTAACTTCAATTCC**TTATGTTGTAGTCGTCGTTG | |
| Er-PrsAnoIDR-F2 | **GAAATAATTCTATG**GACTTGTAAAGATTGACG | |
| PrsAnoIDR-ER-R2 | AATCTTTACAAGTC**CATAGAATTATTTCCTCCCG** | |
| PrsA-R | GACATAGTCAATAGCAATGAC | |
| PrsAnoIDR-check-F | AAACGTCTGAAAGAAATTATCG | Verification of *prsA*Δ$IDR^{8aa}$ |
| PrsAnoIDR-ER-R2 | AATCTTTACAAGTC**CATAGAATTATTTCCTCCCG** | |
| PrsA-F | CGTCCTGTTATTAAGGAAGAC | Construction of *prsA*Δ$IDR^{23aa}$ |
| PrsAstop26-Er-F1 | ATGATCAAACAACATAA**GGAATTGAAGTTAAATTAGATG** | |
| Er-PrsAstop26-R1 | **TTAACTTCAATTCC**TTATGTTGTTTGATCATCAGC | |

| | | |
|---|---|---|
| Er-PrsAnoIDR-F2 | **GAAATAATTCTATG**GACTTGTAAAGATTGACG | |
| PrsAnoIDR-ER-R2 | AATCTTTACAAGTC**CATAGAATTATTTCCTCCCG** | |
| PrsA-R | GACATAGTCAATAGCAATGAC | |
| PrsAnoIDR-check-F | AAACGTCTGAAAGAAATTATCG | Verification of *prsA*ΔIDR$^{23aa}$ |
| PrsAnoIDR-ER-R2 | AATCTTTACAAGTC**CATAGAATTATTTCCTCCCG** | |
| PrsA-F | CGTCCTGTTATTAAGGAAGAC | Construction of *prsA*ΔIDR-IDR$^{wapA}$ |
| PrsA-wapA-F1 | GGCAGCAAGCGGAGCAGAAACAACAACTGTTAC | |
| WapA-prsA-R1 | TGTTGTTTCTGCTCCGCTTGCTGCCGCATATTG | |
| WapA-Er-F2 | AAACAAGTAACCTAA**GGAATTGAAGTTAAATTAGATG** | |
| Er-WapA-R2 | **ACTTCAATTCCTT**AGGTTACTTGTTTTGAAGTTGTTG | |
| Er-PrsAnoIDR-F2 | **GAAATAATTCTATG**GACTTGTAAAGATTGACG | |
| PrsAnoIDR-ER-R2 | AATCTTTACAAGTC**CATAGAATTATTTCCTCCCG** | |
| PrsA-R | GACATAGTCAATAGCAATGAC | |
| PrsAnoIDR-check-F | AAACGTCTGAAAGAAATTATCG | Verification of *prsA*ΔIDR-IDR$^{wapA}$ |
| PrsAnoIDR-ER-R2 | AATCTTTACAAGTC**CATAGAATTATTTCCTCCCG** | |
| PrsA-F | CGTCCTGTTATTAAGGAAGAC | Construction of *prsA*ΔIDR-IDR$^{PrsA1\text{-}GAS}$ |
| PrsA-GAS-F1 | GGCAGCAAGCGGAAGTGAAAGTTCAACAACCAG | |
| GAS-PrsA-R1 | TGAACTTTCACTTCCGCTTGCTGCCGCATATTG | |
| GAS-Er-F2 | CTGCACAATAA**GGAATTGAAGTTAAATTAGATG** | |
| Er-GAS-R2 | **TTTAACTTCAATTCC**TTATTGTGCAGCTGGCTC | |
| Er-PrsAnoIDR-F2 | **GAAATAATTCTATG**GACTTGTAAAGATTGACG | |
| PrsAnoIDR-ER-R2 | AATCTTTACAAGTC**CATAGAATTATTTCCTCCCG** | |
| PrsA-R | GACATAGTCAATAGCAATGAC | |
| PrsAnoIDR-check-F | AAACGTCTGAAAGAAATTATCG | Verification of *prsA*ΔIDR-IDR$^{PrsA1\text{-}GAS}$ |
| PrsAnoIDR-ER-R2 | AATCTTTACAAGTC**CATAGAATTATTTCCTCCCG** | |
| PIN1-ER-F1 | CAGATCTTCTGGAAGAGTTTACTACTTTAACCACATTACCAATGCATCACAATGGGAAAGACCATCTGGGAATTAA**GGAATTGAAGTTAAATTAGATG** | Construction of the *prsA*-PIN1 strain |
| PIN1-prsA-R1 | GTAGTAAACTCTTCCAGAAGATCTGCTCATACGTTTCTCCCATCCTGGAGGTAACTTCTCTTCATCAGCTCCACCACCTCCGCTTGCTGCCGCATATT | |
| FL-ER-F1 | TATAAAGATCATGATATCGATTACAAGGATGACGATGACAAGTAA**GGAATTGAAGTTAAATTAGATG** | Construction of the *prsA*-IDR$^{NG}$-3xFLAG strain |
| FL-NG-R1 | TGTAATCGATATCATGATCTTTATAATCACCGTCATGGTCTTTGTAGTCTTCCGCAGCATTGTTTTC | |

| | | |
|---|---|---|
| NG-Er-F1 | AAATGCTGCTGGAGGAGCAGCCGATGATCAAAATGGTGCTGCCGAAAACAATGCTGCGGAATAA**GGAATTGAAGTTAAATTAGATG** | Construction of the *prsA*-IDR$^{NG}$ strain |
| Er-NG-R1 | ATCGGCTGCTCCTCCAGCAGCATTTCCTGCTGCACCATTACCGTTATTTCCGCCATTATTCCCATTTCCGCTTGCTGCCGCATATT | |
| pknB-XhoI-F | GTACCGGGCCCCCCTCGAGATCAAACTAATCAAACAG | Construction of 2221*pknB*$\Delta$IDR |
| Sp-pknB-R1 | TTGGTCGTTATTCTGGATAAAGGTAAAGACTA | |
| pknB-Sp-F1 | TTATCCAGAATAA**CGACCAAAATAGTGAGGAG** | |
| pknB-Sp-R2 | AGAGTGTG**GATCCTTATAATTTTTTTAATCTG** | |
| Sp-pknB-F2 | AATTATAAGGATCCACACTCTTCTAGTAGCTC | |
| pknB-BamH-R | CGCTCTAGAACTAGTGGATCCGTATATGGAACTAACATC | |
| pknB-Up | CCAACAGTGTTAACACAGG | Verification of 2221*pknB*$\Delta$IDR |
| pknB-chF | TGCAGTCAATACCTTAACAGC | |
| prsA1d-F-XhoI | GTACCGGGCCCCCCTCGAGCTTGTTCATTATATGCG | Construction of 2221$\Delta$*prsA1* |
| Sp-prsA1d-R1 | ATTTTGGTCGTTACTTCTGAGCTCCTTTTTATC | |
| prsA1d-Sp-F1 | GAGCTCAGAAGTAA**CGACCAAAATAGTGAGGAG** | |
| prsA1d-Sp-R2 | ATAAAATCCTACT**GATCCTTATAATTTTTTTAATCTG** | |
| Sp-prsA1-F2 | AATTATAAGGATCAGTAGGATTTTATTGACTAATCAG | |
| prsA1d-BamHI | CGCTCTAGAACTAGTGGATCCCCACTCAATAGCTTC | |
| prsA1d-Up | CATAATATGCGTCGTCCTG | Verification of 2221$\Delta$*prsA1* |
| prsA1d-chF | AAAGATAAAAAGGAGCTCAGAAG | |
| prsA1d-chR | CAACCGTCACTTTTCTGTATC | |
| prsA-XhoI-F | GTACCGGGCCCCCCTCGAGGTGATACAATTAGCGTTAG | Construction of 2221*prsA1*$\Delta$IDR |
| Sp-prsA-R1 | TTTGGTCGTTATTTTTGACCAAGATTTGCAT | |
| prsA-Sp-F1 | GGTCAAAAATAA**CGACCAAAATAGTGAGGAG** | |
| prsA-Sp-R2 | AGCTTTAG**GATCCTTATAATTTTTTTAATCTG** | |
| Sp-prsA-F2 | AATTATAAGGATCCTAAAGCTGCAAGTGAAAG | |
| prsA-BamHI-R | CGCTCTAGAACTAGTGGATCCAAGCATTTCAAACATAG | |
| prsA2d2-XhoI | GTACCGGGCCCCCCTCGAGCCATGTTAAGCTTGG | Construction of 2221*prsA2* $\Omega$pKV304 |
| prsA2d2-BamH | CGCTCTAGAACTAGTGGATCCTTTCAAGCGTTTTTGGTAC | |
| prsA2d2-Up | CGTCTCGATGAAAGGAGAC | Verification of 2221*prsA2* $\Omega$pKV304 |

[a] Restriction sites are underlined.

[b] Extensions complementary to the antibiotic resistance cassettes are in bold.

**Supplementary Table 8.** Primers used to construct *S. pneumoniae* strains.

| Primers used to construct strains | | | |
|---|---|---|---|
| Primer | Sequence (5' to 3') | Template[a] | Amplicon Product |
| For construction of E221 (Δ*prsA*(*spd_0868*)::$P_c$-*erm*) | | | |
| P514 | TCAACGATGCCTTTGCAGTCTTTGAACGCC | D39 | 5' fragment with 60 bp of 5' *prsA* |
| P516 | CATTATCCATTAAAAATCAAACGGATCCTAAGCTGCTAAAGTTGCTACTGATAATAGTGT | | |
| Kan rpsL forward | TAGGATCCGTTTGATTTTTAATGGATAATG | Pc-*erm* cassette[b] | Pc-*erm* |
| Kan rpsL reverse | GGGCCCCTTTCCTTATGCTTTTG | | |
| P517 | CAAAAGCATAAGGAAAGGGGCCCTTTACCCAATATATCGGTGGTGGAGATT | D39 | 3' fragment with 60 bp of 3' *prsA* |
| P515 | AAGCTGTTCTTTGAGCAGCCTACAGCTAGCTTCC | | |
| For construction of K213 (Δ*prsA*::$P_c$-*kan-rpsL*$^+$) | | | |
| P514 | TCAACGATGCCTTTGCAGTCTTTGAACGCC | D39 | 5' fragment with 60 bp of 5' *prsA* |
| P516 | CATTATCCATTAAAAATCAAACGGATCCTAAGCTGCTAAAGTTGCTACTGATAATAGTGT | | |
| Kan rpsL forward | TAGGATCCGTTTGATTTTTAATGGATAATG | $P_c$-[*kan-rpsL*$^+$] cassette[b] | $P_c$-[*kan-rpsL*$^+$] |
| Kan rpsL reverse | GGGCCCCTTTCCTTATGCTTTTG | | |
| P517 | CAAAAGCATAAGGAAAGGGGCCCTTTACCCAATATATCGGTGGTGGAGATT | D39 | 3' fragment with 60 bp of 3' *prsA* |
| P515 | AAGCTGTTCTTTGAGCAGCCTACAGCTAGCTTCC | | |
| For construction of IU18448 (Δ*gtrB* (*spd_1431*)::$P_c$-*erm*) | | | |
| TT1427 | GTGGTGGATAATCTTGTCAATAGCAATCGTAAGAG | D39 | 5' fragment with 102 bp of 5' *gtrB* |
| TT1429 | CATTATCCATTAAAAATCAAACGGATCCTAGATTTCTGTTTCCAAATCTGGAAGTAAAGC | | |
| Kan rpsL forward | TAGGATCCGTTTGATTTTTAATGGATAATG | $P_c$-*erm* cassette[b] | $P_c$-*erm* |
| Kan rpsL reverse | GGGCCCCTTTCCTTATGCTTTTG | | |
| TT1430 | CAAAAGCATAAGGAAAGGGGCCCACCATTGGGATTCTCGGTAAGTATATCAGTAAGATTT | D39 | 3' fragment with 114 bp + stop of 3' *gtrB* |
| TT1428 | GCCATATAGGTCGCTGCACGATACATAGCTCC | | |
| For construction of IU18454 (Δ*gtrB* (*spd_1431*)::$P_c$-*kan-rpsL*$^+$) | | | |
| TT1427 | GTGGTGGATAATCTTGTCAATAGCAATCGTAAGAG | D39 | 5' fragment with 102 bp of 5' *gtrB* |
| TT1429 | CATTATCCATTAAAAATCAAACGGATCCTAGATTTCTGTTTCCAAATCTGGAAGTAAAGC | | |
| Kan rpsL forward | TAGGATCCGTTTGATTTTTAATGGATAATG | $P_c$-*kan-rpsL*$^+$ cassette[b] | $P_c$-*kan-rpsL*$^+$ |

| | | | |
|---|---|---|---|
| Kan rpsL reverse | GGGCCCCTTTCCTTATGCTTTTG | | |
| TT1430 | CAAAAGCATAAGGAAAGGGGCCCACCATTGGGATTCTCGGTAAGTATATCAGTAAGATTT | D39 | 3' fragment with 114 bp + stop of 3' *gtrB* |
| TT1428 | GCCATATAGGTCGCTGCACGATACATAGCTCC | | |
| For construction of IU18456 (Δ*pgtC2* (Δ*spd_2058*)::$P_c$-*kan-rpsL*$^+$) | | | |
| TT1431 | GCAGGCAACATCACGTAAGAAAATGCTTG | D39 | 5' fragment with 120 bp of 5' *spd_2058* |
| TT1433 | CATTATCCATTAAAAATCAAACGGATCCTATAGAGGAGAGTTGTCGCTATTCCAGTAGAT | | |
| Kan rpsL forward | TAGGATCCGTTTGATTTTTAATGGATAATG | $P_c$-*kan-rpsL*$^+$ cassette[c] | $P_c$-*kan-rpsL*$^+$ |
| Kan rpsL reverse | GGGCCCCTTTCCTTATGCTTTTG | | |
| TT1434 | CAAAAGCATAAGGAAAGGGGCCCGGAGCAATCTGTTCCTTTACTTCTCTCTTACTATTTG | D39 | 3' fragment with 72 bp + stop of 3' *spd_2058* |
| TT1432 | CAGGAGTTGGTTAATTACAATCAGGATCTGGGC | | |
| For construction of IU18715, IU18719 (Δ*bgaA*::*kan*-$P_{ftsA}$-*gtrB*) | | | |
| TT657 | CGCCCCAAGTTCATCACCAATGACATCAAC | IU9621 | 5' Δ*bgaA*::*kan*-$P_{ftsA}$ |
| TT1480 | TAAACAAGGGACGATGATTGAAATCATCATTACATCGCTTCCTCTCTATCTTCCAAGT | | |
| TT1481 | ACTTGGAAGATAGAGAGGAAGCGATGTAATGATGATTTCAATCATCGTCCCTTGTTTA | D39 | *gtrB*$^+$ |
| TT1450 | GCAACTGGTTTATGAGAAAGTAAGTTCTTTCACTTTATTTTTTCTGTAAAATCAGGAAGG | | |
| TT1451 | TCCTGATTTTACAGAAAAAATAAAGTGAAAGAACTTACTTTCTCATAAACCAGTTGCTGC | D39 | *bgaA'* to downstream |
| CS121 | GCTTTCTTGAGGCAATTCACTTGGTGC | | |
| For construction of IU20087, IU20089 (Δ*gtrB* (*spd_1431*)::$P_c$-*aad9*) | | | |
| TT1427 | GTGGTGGATAATCTTGTCAATAGCAATCGTAAGAG | D39 | 5' fragment with 102 bp of 5' *gtrB* |
| TT1429 | CATTATCCATTAAAAATCAAACGGATCCTAGATTTCTGTTTCCAAATCTGGAAGTAAAGC | | |
| Kan rpsL forward | TAGGATCCGTTTGATTTTTAATGGATAATG | IU6987 | $P_c$-*aad9* |
| Kan rpsL reverse | GGGCCCCTTTCCTTATGCTTTTG | | |
| TT1430 | CAAAAGCATAAGGAAAGGGGCCCACCATTGGGATTCTCGGTAAGTATATCAGTAAGATTT | D39 | 3' fragment with 114 bp + stop of 3' *gtrB* |
| TT1428 | GCCATATAGGTCGCTGCACGATACATAGCTCC | | |
| For construction of IU20242 (*prsA*(ΔIDR(S304-E313)) | | | |
| P514 | TCAACGATGCCTTTGCAGTCTTTGAACGCC | D39 | 5' fragment with *prsA* (aa 1-303) |
| TT1642 | GTTTTTTCCCTGACTCATTGATTTGGACTAATCTCCACCACCGATATATTGGGTAAAGAT | | |
| TT1643 | ATCTTTACCCAATATATCGGTGGTGGAGATTAGTCCAAATCAATGAGTCAGGGAAAAAAC | D39 | 3' fragment with *prsA* (aa 314 to end) |
| P515 | AAGCTGTTCTTTGAGCAGCCTACAGCTAGCTTCC | | |
| For construction of IU20269 (*pbp1a*(ΔIDR(S669-P719)) | | | |

| | | | |
|---|---|---|---|
| P234 | CCCTTGTGTTCATAGCGAGGATAAGCA | D39 | 5’ fragment with *pbp1a* (aa 1-668) |
| TT1638 | CACCCAGAAAAATCTGGATGATAAATGTTATTCAGTTGATGGGGGTTGTTGTGG | | |
| TT1639 | ACAACAACCCCCATCAACTGAATAACATTTATCATCCAGATTTTTCTGGGTG | D39 | 3’ fragment with *pbp1a* (aa 720 to end) |
| P235 | AGGCAAGCCTGCAACCATGGTCTTGAAA | | |
| For construction of IU20273 (*pbp1b*(ΔIDR(S794-R821)) | | | |
| P222 | CGTTCGTGTGGCGCTGCTTCAAATTGTT | D39 | 5’ fragment with *pbp1b* (aa 1-793) |
| TT1640 | CCATTGATTGGCCACTTCATTTAGAGAATTATGGAGTTGGTAGACTCCCCACAATAC | | |
| TT1641 | GTATTGTGGGGAGTCTACCAACTCCATAATTCTCTAAATGAAGTGGCCAATCAATGG | D39 | 3’ fragment with *pbp1b* (aa 822 to end) |
| P522 | AACGGCAACCACCAAAGGAGAAACCAAGGA | | |
| For construction of IU20663 (*mapZ*(ΔIDR(T318-S357)) | | | |
| P1523 | GAGGTCTCTATTCTCAAAGATGTGGCAACTGTC | D39 | 5’ fragment with *mapZ* (aa 1-317) |
| TT1697 | GCACTCGAGAGACCCATATTGACTTCTTGCTGGCTCTTACCAAGACTGATAGCC | | |
| TT1698 | GGCTATCAGTCTTGGTAAGAGCCAGCAAGAAGTCAATATGGGTCTCTCGAGTGC | D39 | 3’ fragment with *mapZ* (aa 358 to end) |
| P1526 | AATTGCATATCACCGTACTCAATACCATTGTG | | |
| For construction of TIGR4Δ*pgtC2*[c] | | | |
| SP2231-f | GATTGATGGTGAAACTATCTTG | TIGR4 | 5’ fragment with *pgtC2* |
| Kan-SP2231-r1 | **CAGTATTTAAAGATACC**GATTTCATGTTACCTCGTAG | | |
| SP2231-Kan-f1 | GAGGTAACATGAAATC**GGTATCTTTAAATACTGTAG** | pOSKAR | a nonpolar kanamycin resistance cassette |
| Kan-SP2231-F2 | **TGAATTGTTTTAGTAC**AAGTGGTCTAGGGCTAAAC | | |
| SP2231-Kan-R2 | TAGCCCTAGACCACTT**GTACTAAAACAATTCATCCAG** | TIGR4 | 3’ fragment with *pgtC2* |
| SP2231-R | GTTTCACGTGAAACTTAATAG | | |
| SP2231-check-f | CAGTTGGGTTAACCAACTGAG | TIGR4Δ*pgtC2* | Verification of TIGR4Δ*pgtC*: Kan cassette and 5’ and 3’ DNA flanking regions of *pgtC2* |
| SP2231-check-r | CCAAGTAATAACTAGATAGG | | |

[a] Genomic DNA of indicated *S. pneumoniae* strains was used as templates for PCR reactions, except for $P_c$-*kan-rpsL*$^+$ and $P_c$-*erm* cassettes.
[b] $P_c$-*erm* and $P_c$-*kan-rpsL*$^+$ cassettes are described in [19].
[c] Extensions complementary to the antibiotic resistance cassettes are in bold.

**Supplementary Table 9.** Spontaneous mutations detected in the mutants of *S. mutans* Xc by whole-genome sequencing.

| Strain | Target insertion of antibiotic resistance cassette [a] | Additional targeted mutations [a] | Spontaneous mutations [a] | Annotation |
|---|---|---|---|---|
| *prsA*ΔIDR | n.a. | Smu.648 S300Stop | n.d. [b] | |
| *prsA*ΔIDR Δ*pgfS* | Smu.2067 | Smu.648 S300Stop | n.d. | |
| *prsA*-3XFLAG Δ*pgfS* | Smu.2067 | Smu.648 Stop334A->3X-FLAG | n.d. | |
| *prsA*ΔIDR-3XFLAG Δ*pgfS* | Smu.2067 | Smu.648 S300D->3X-FLAG | n.d. | |
| *prsA*ΔIDR-IDR$^{\text{wapA}}$ Δ*pgfS* | Smu.2067 | Smu.648 S300->IDR from Smu.987 | n.d. | |
| *prsA*ΔIDR-IDR$^{\text{PrsA1-GAS}}$ Δ*pgfS* | Smu.2067 | Smu.648 S300->IDR from MGAS2221_1146[c] | Smu.932 L152F (TT$\underline{\text{G}}$→TT$\underline{\text{T}}$) | Predicted methylcobamide:CoM methyltransferase isozyme A |
| *prsA*ΔIDR$^{\text{8aa}}$ | n.a. | Smu.648 A326Stop | n.d. | |
| *prsA*ΔIDR$^{\text{23aa}}$ | n.a. | Smu.648 A311Stop | n.d. | |
| *prsA*ΔIDR$^{\text{8aa}}$Δ*pgfS* | Smu.2067 | Smu.648 A326Stop | n.d. | |
| *prsA*ΔIDR$^{\text{23aa}}$Δ*pgfS* | Smu.2067 | Smu.648 A311Stop | n.d. | |
| *prsA*-PIN1 | n.a. | Smu.648 S300->PIN1 | n.d. | |
| *prsA*-PIN1 Δ*pgfS* | Smu.2067 | Smu.648 S300->PIN1 | n.d. | |
| *prsA*-IDR$^{\text{NG}}$ | n.a. | Smu.648 S300->NG | n.d. | |
| *prsA*-IDR$^{\text{NG}}$ Δ*pgfS* | Smu.2067 | Smu.648 S300->NG | n.d. | |

[a] corresponds to genes in *S. mutans* UA159 (GenBank: AE014133.2)

[b] n.d. indicates not detected

[c] corresponds to genes in *S. pyogenes* MGAS 2221 (GenBank: CP043530.1)

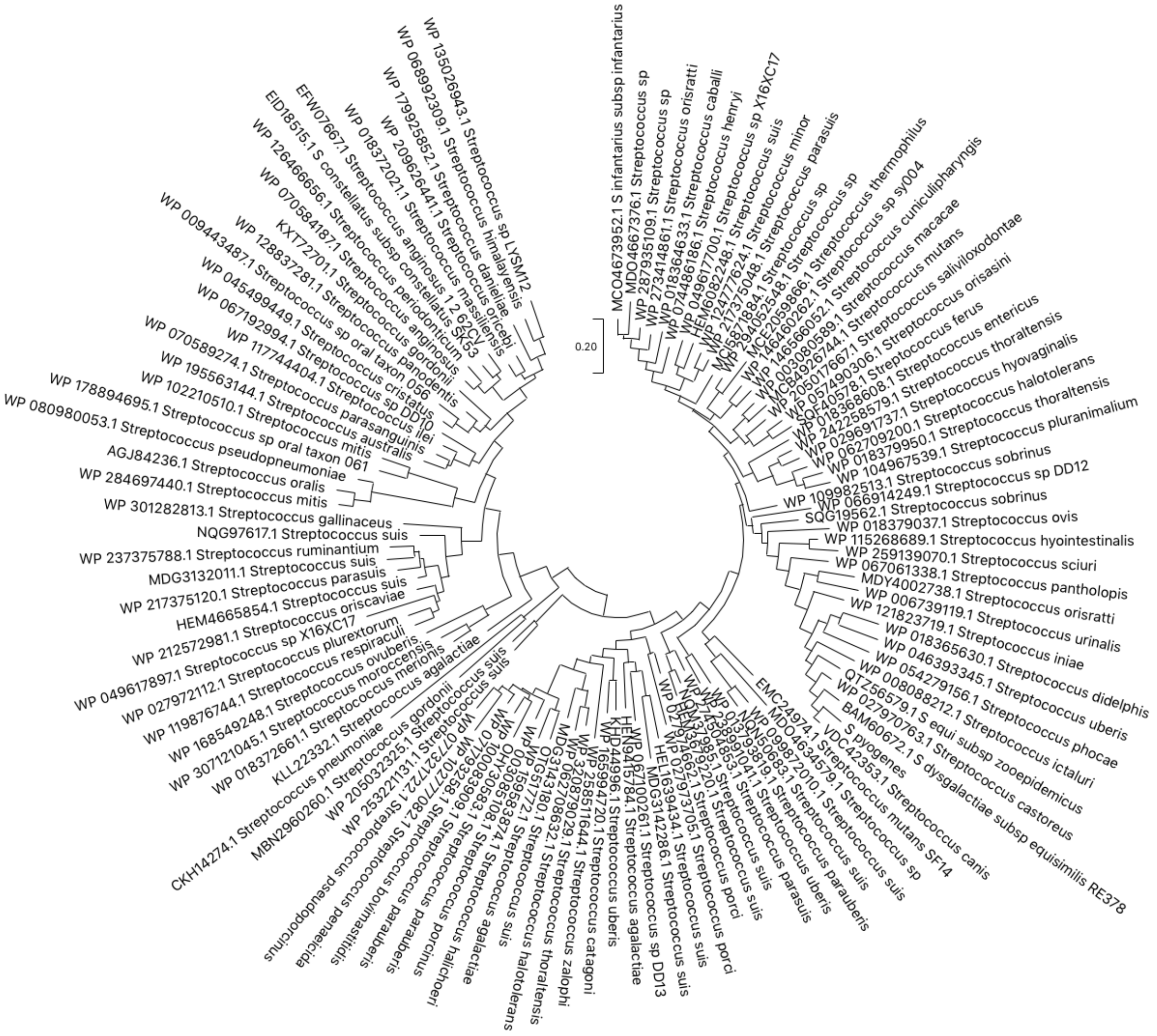


**Supplementary Fig. 1. The phylogenetic tree of GtrB$^{Spy}$ homologs.**

Sequences of GtrB$^{Spy}$ homologs from streptococci were retrieved from the NCBI database using BLAST [20]. The redundant sequences were removed using CD-HIT with 90% sequence identity cut-off [21]. The evolutionary history was inferred by using the Maximum Likelihood method and JTT matrix-based model [22]. The tree with the highest log likelihood (-28587.63) is shown. Initial tree(s) for the heuristic search were obtained automatically by applying Neighbor-Join and BioNJ algorithms to a matrix of pairwise distances estimated using the JTT model, and then selecting the topology with superior log likelihood value. The tree is drawn to scale, with branch lengths measured in the number of substitutions per site. This analysis involved 114 amino acid sequences. There were a total of 398 positions in the final dataset. Evolutionary analyses were conducted in MEGA X [23].

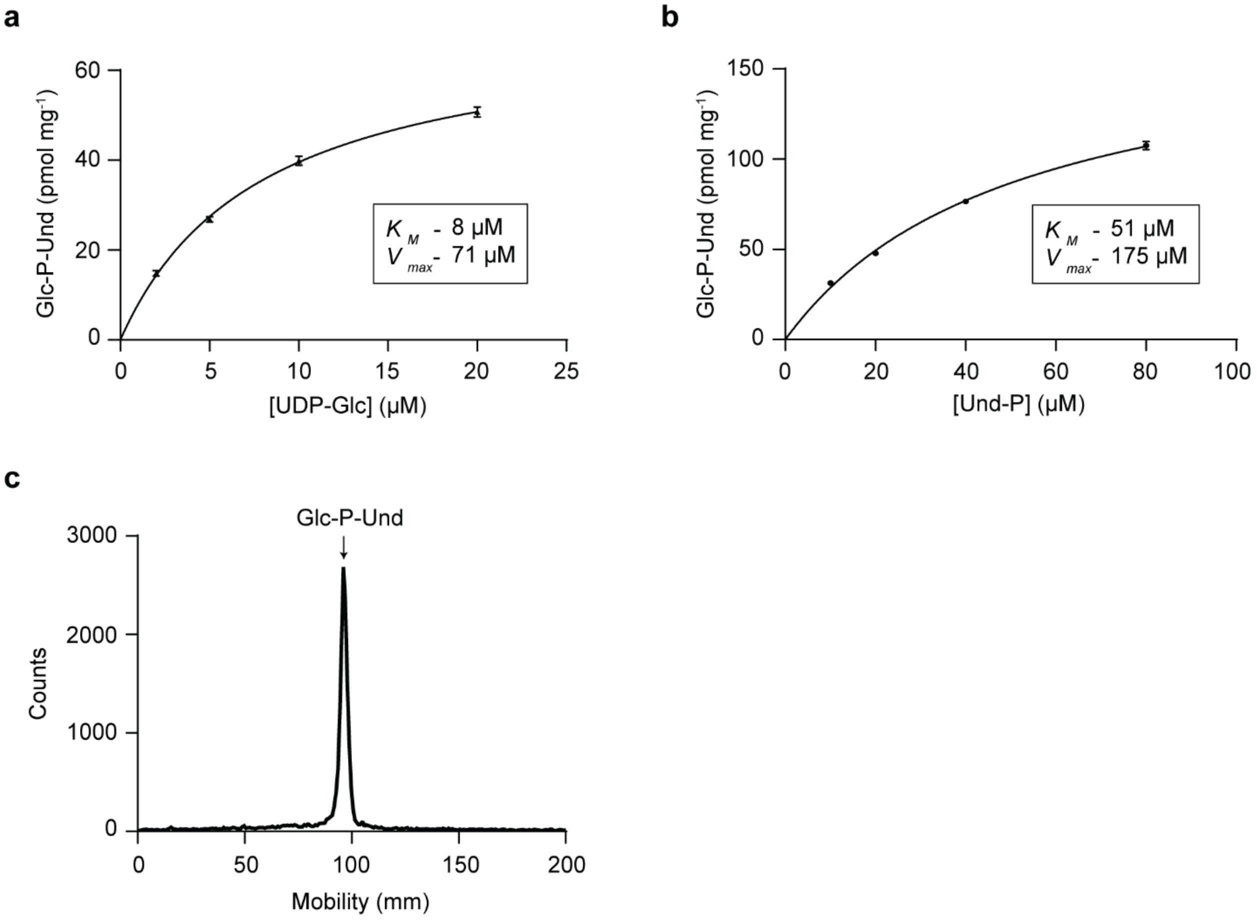


**Supplementary Fig. 2. Kinetic- and TLC analysis of $GtrB^{Spy}$ Glc-P-Und synthase activity in the *E. coli* pBAD33_$GtrB^{Spy}$ membrane fractions.**

(**a** and **b**) Kinetic analysis of Glc-lipid synthase in the *E. coli* pBAD33_$GtrB^{Spy}$ membrane fractions. Dependence of $GtrB^{Spy}$ activity on concentrations of UDP-Glc (**a**) and Und-P (**b**). Membrane fractions were analyzed for the formation of $[^3H]$Glc-lipids in the presence of increasing amounts Und-P or UDP-$[^3H]$Glc as described in Methods. Apparent kinetic parameters for UDP-Glc and Und-P were calculated using GraphPad Prism 10 software using the Michaelis-Menten enzyme kinetics tool. The experiments were performed independently three times. (**c**) Representative TLC analysis of $[^3H]$Glc-lipids extracted from *in vitro* incubations of UDP-$[^3H]$Glc with the *E. coli* pBAD33_$GtrB^{Spy}$ membrane fractions. The experiment was performed independently three times and yielded the same results. Source data for **a, b** and **c** are provided as a Source Data file.

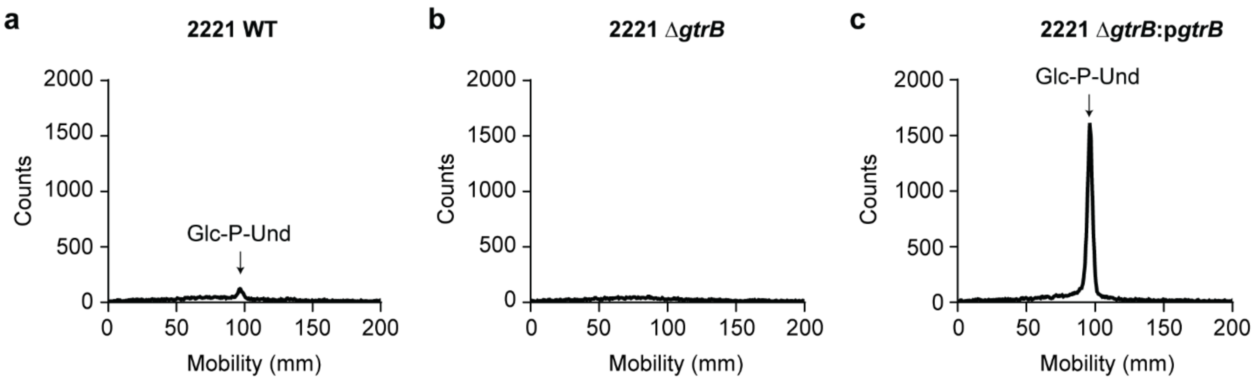


**Supplementary Fig. 3. Analysis of GtrB activity in the membrane fractions of *S. pyogenes* strains.**

(**a**, **b** and **c**) TLC of [$^3$H]Glc-lipids from *in vitro* incubations of membrane fractions from *S. pyogenes* WT (**a**), Δ*gtrB* (**b**), and Δ*gtrB*:p*gtrB* (**c**) with UDP-[$^3$H]Glc as described in Methods. Following incubation products were subjected to mild alkaline de-acylation with 0.1 N KOH to eliminate glucosyldiglyceride. Recovered [$^3$H]Glc-lipids were analyzed by TLC. The experiment was performed independently two times and yielded the same results. Source data for **a, b** and **c** are provided as a Source Data file.

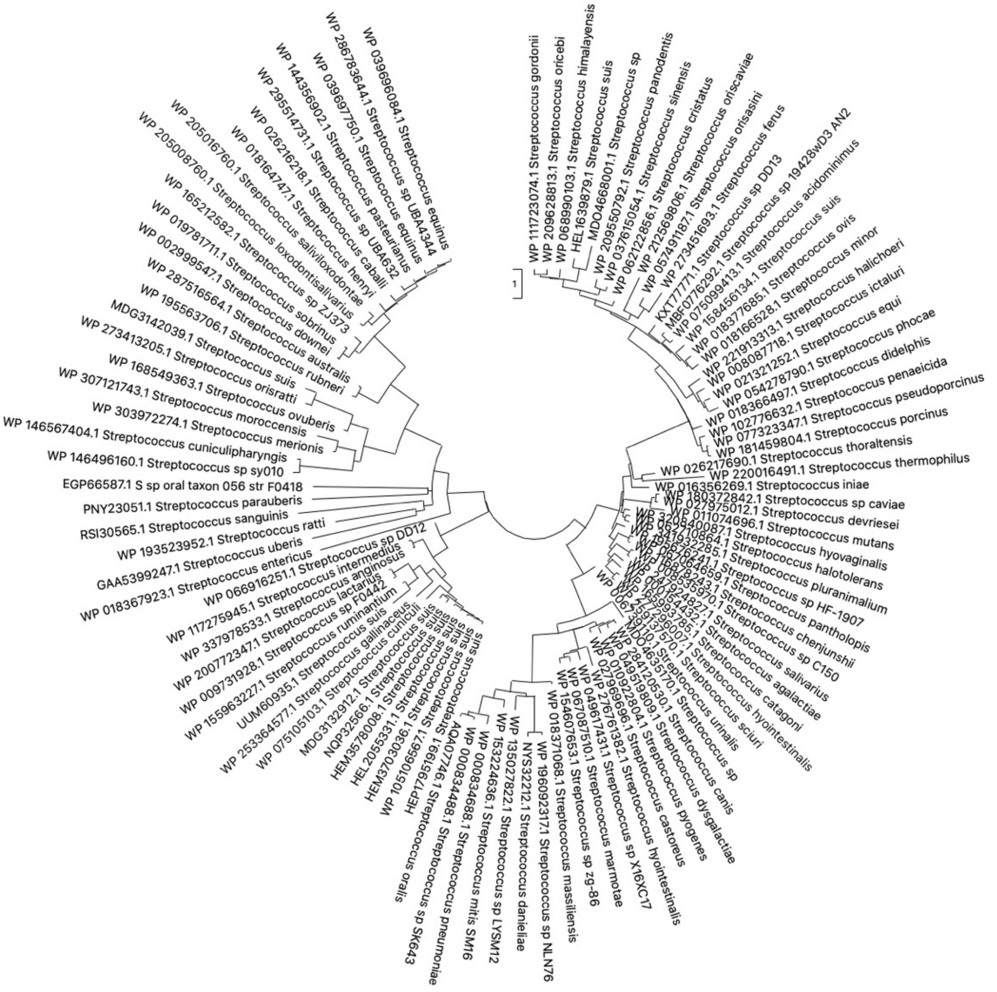


**Supplementary Fig. 4. The phylogenetic tree of PgtC2$^{Spy}$ homologs.**

Sequences of PgtC2$^{Spy}$ homologs from streptococci were retrieved from the NCBI database using BLAST [20]. The redundant sequences were removed using CD-HIT with 90% sequence identity cut-off [21]. The evolutionary history was inferred by using the Maximum Likelihood method and JTT matrix-based model [22]. The tree with the highest log likelihood (-156062.16) is shown. Initial tree(s) for the heuristic search were obtained automatically by applying Neighbor-Join and BioNJ algorithms to a matrix of pairwise distances estimated using the JTT model, and then selecting the topology with superior log likelihood value. The tree is drawn to scale, with branch lengths measured in the number of substitutions per site. This analysis involved 104 amino acid sequences. There were a total of 901 positions in the final dataset. Evolutionary analyses were conducted in MEGA X [23].

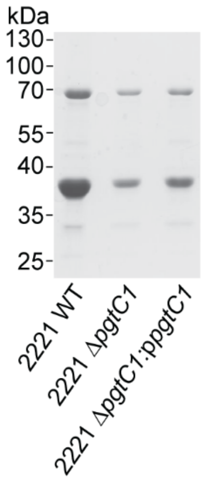


**Supplementary Fig. 5. Lectin affinity purification of *S. pyogenes* membrane glycoproteins.** Membrane proteins obtained from WT, Δ*pgtC1*, and Δ*pgtC1:*p*pgtC1* were purified by concanavalin A affinity chromatography as described in Methods. Proteins were separated on a 15% SDS-PAGE gel in Tris-glycine buffer and visualized by Coomassie staining. The experiment was performed independently two times and yielded the same results. A representative image from one experiment is shown. Source data are provided as a Source Data file.

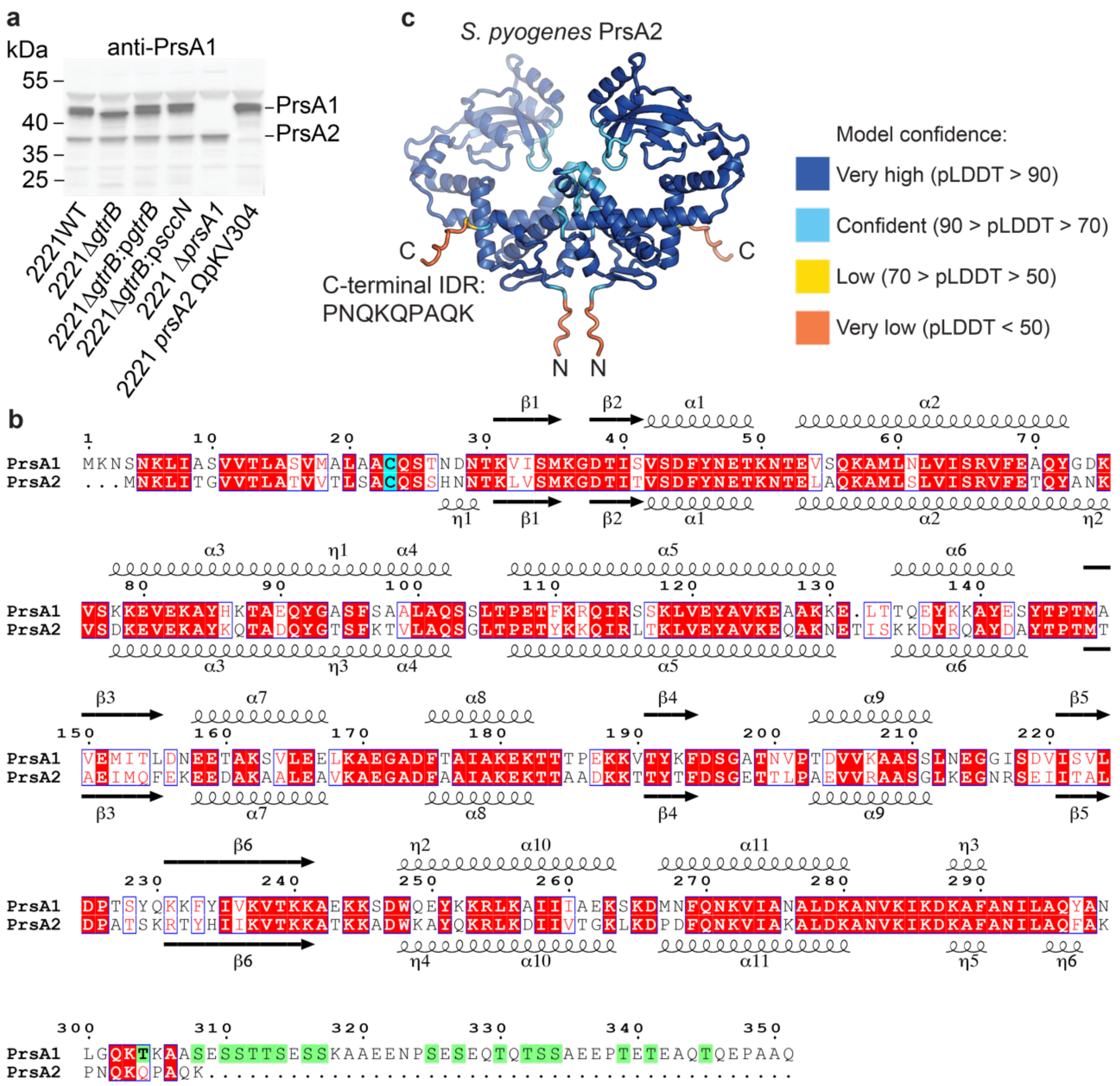


**Supplementary Fig. 6. *S. pyogenes* PrsA2 does not carry Ser/Thr IDR in contrast to PrsA1** (**a**) Immunoblot analysis of PrsA1 and PrsA2 in different *S. pyogenes* mutant strains. Proteins were detected by immunoblotting using anti-PrsA1 antibodies that recognize both PrsA1 and PrsA2, as these two proteins share approximately 65% sequence identity. Proteins were separated on a 4-12% SurePAGE™ Bis-Tris gel (Genscript) in MES buffer. The experiment was performed independently two times and yielded the same results. A representative image from one experiment is shown. (**b**) Sequence alignment of *S. pyogenes* PrsA1 and PrsA2. The AlphaFold2 predicted secondary structure elements of PrsA1 and PrsA2 are shown at the top and bottom, respectively. The acylated N-terminal cysteine residues are highlighted in cyan. The serine and threonine residues in the C-terminal IDR of PrsA1 are highlighted in green. The alignment was made and rendered using Clustal Omega and ESPript [24, 25]. (**c**) AlphaFold2 model of *S. pyogenes* PrsA2 dimer is shown in cartoon representation. The colors correspond to per-residue predicted local distance difference test (pLDDT) values. The amino and carboxy termini are labeled N and C, respectively. The 9 residue C-terminal IDR sequence of *S. pyogenes* PrsA2 is shown. Source data for **a** are provided as a Source Data file.

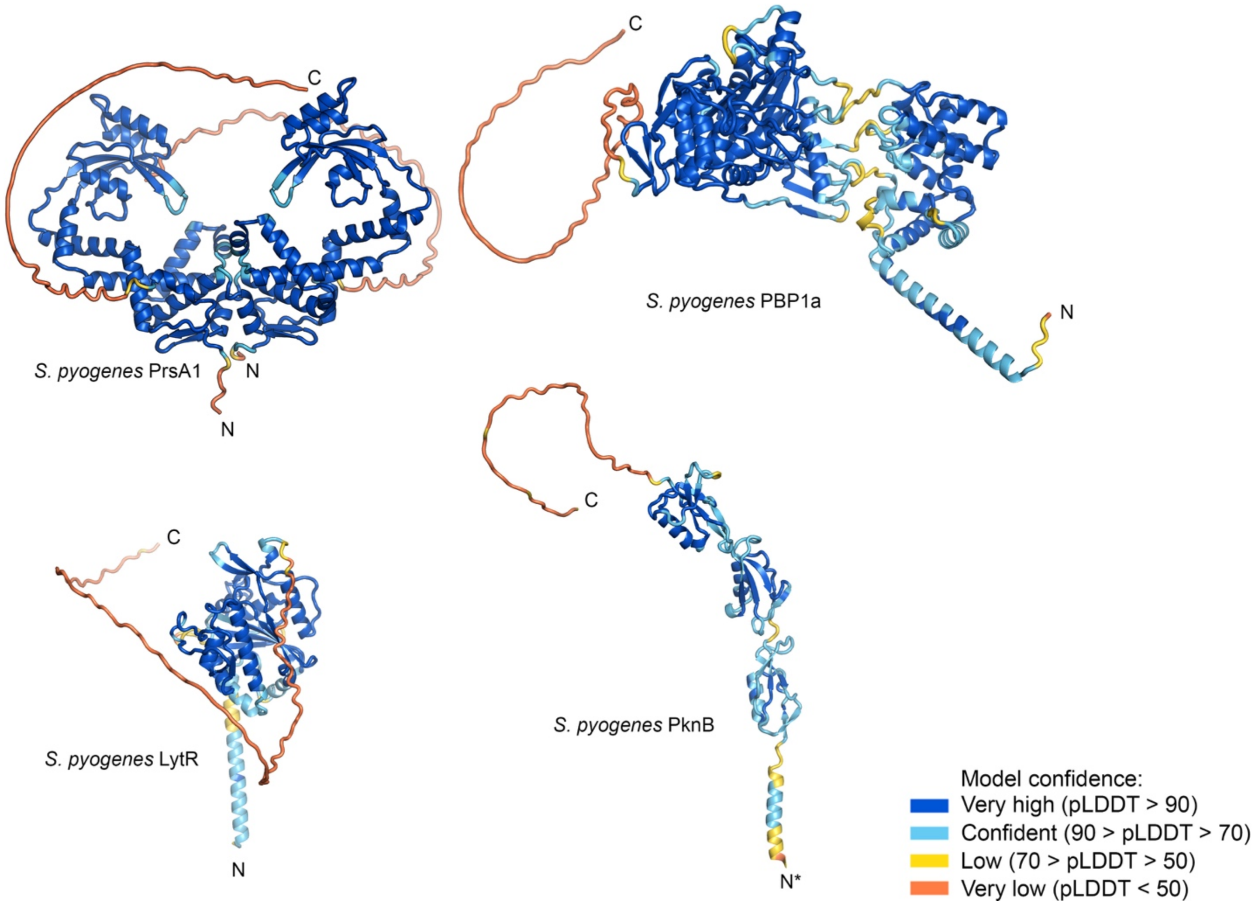


**Supplementary Fig. 7. AlphaFold2 structural models of glycoproteins identified in *S. pyogenes*.**

The colors correspond to pLDDT values. The proteins' amino and carboxy termini are labeled N and C, respectively. For clarity, one monomer of PknB is displayed for clarity. The cytoplasmic domain of PknB and the PrsA lipid anchor are not shown. An asterisk indicates the location of the cytoplasmic domain that is not shown in the image.

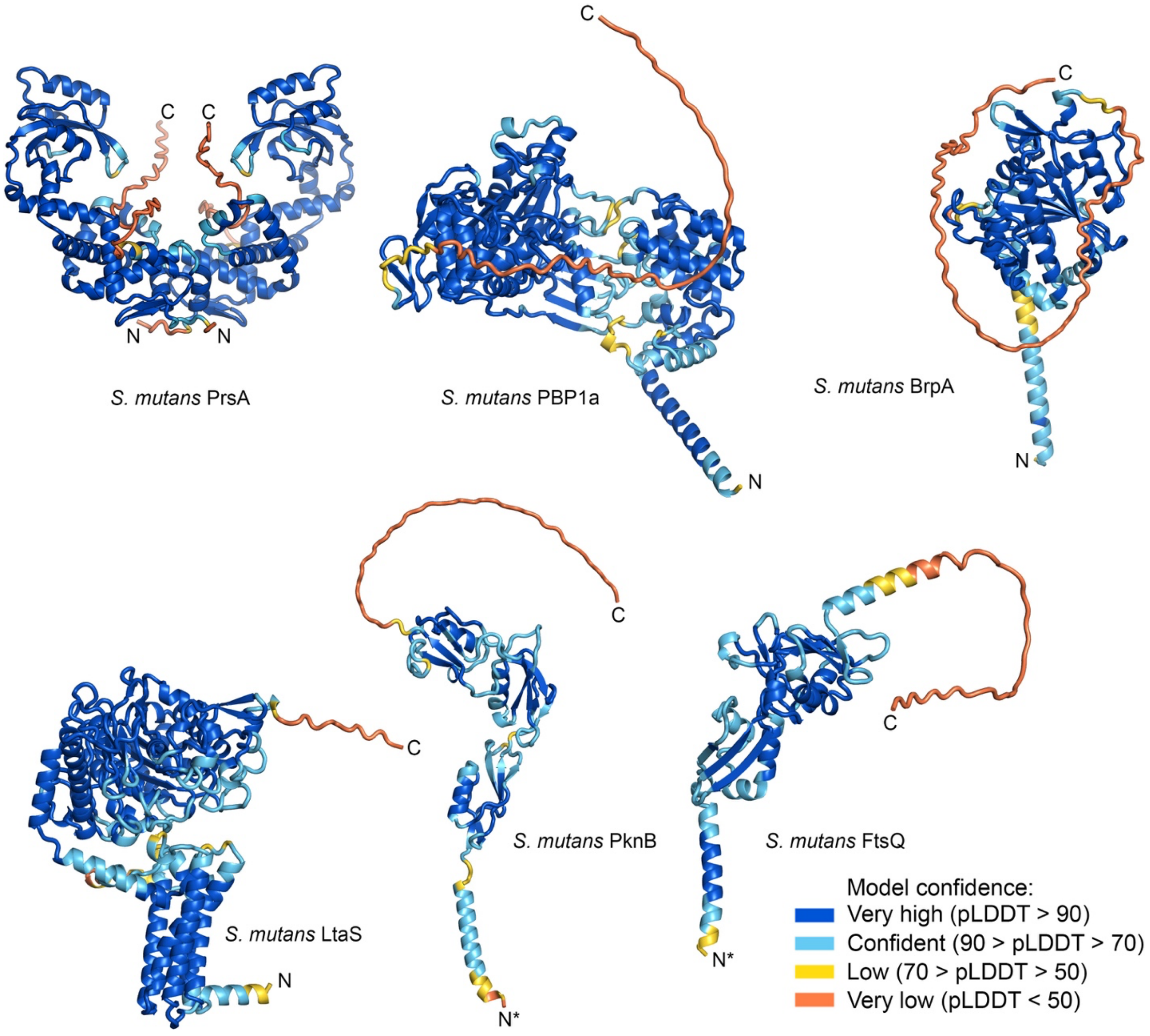


**Supplementary Fig. 8. AlphaFold2 structural models of glycoproteins identified in *S. mutans*.**

The colors correspond to pLDDT values. The cytoplasmic domains of PknB and FtsQ, and the PrsA lipid anchor are omitted for clarity. An asterisk indicates the location of the cytoplasmic domain that is not shown in the image. One monomer of PknB is shown for clarity. The proteins' amino and carboxy termini are labeled N and C, respectively.

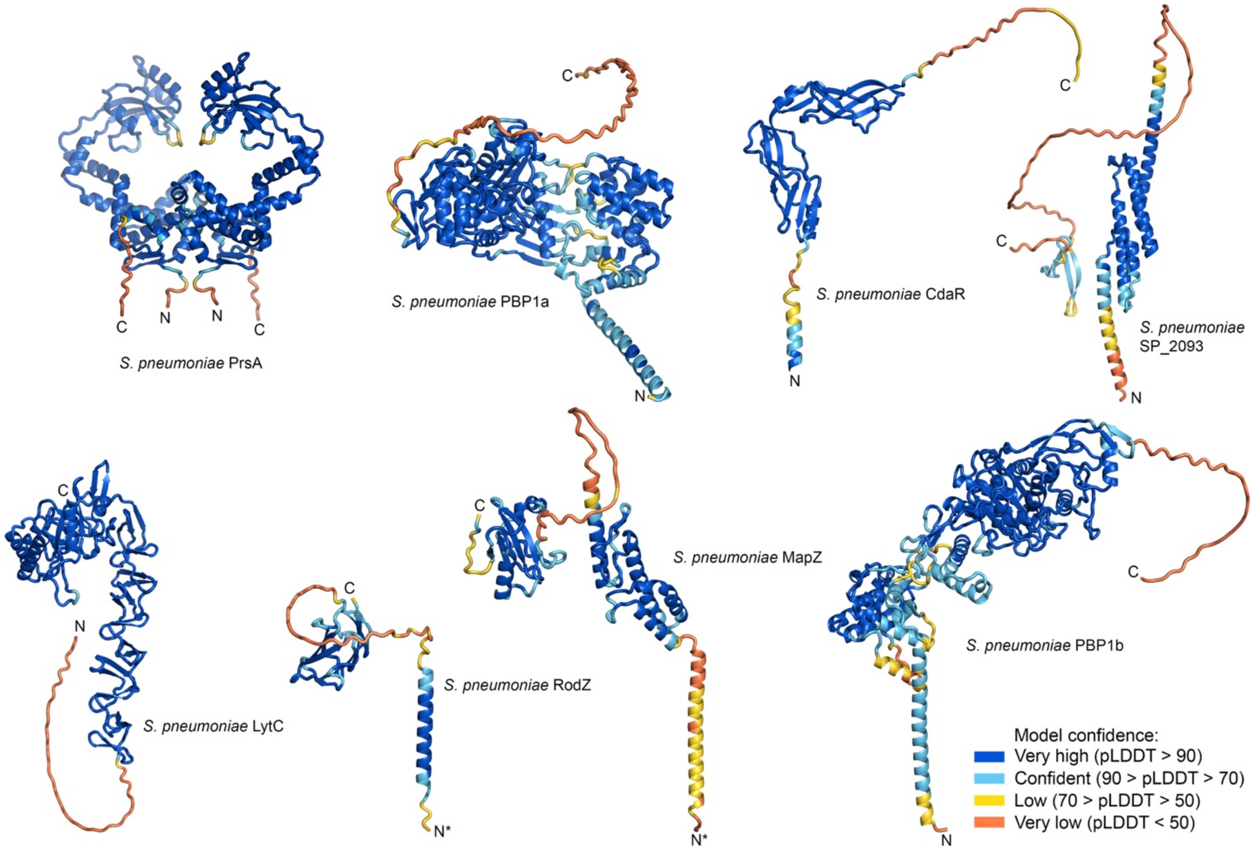


**Supplementary Fig. 9. AlphaFold2 structural models of glycoproteins identified in *S. pneumoniae*.**

The colors correspond to per-residue predicted local distance difference test (pLDDT) values. The cytoplasmic domains of RodZ and MapZ, as well as the lipid anchor of PrsA, are omitted for clarity. An asterisk indicates the location of the cytoplasmic domain that is not shown in the image. The proteins' amino and carboxy termini are labeled N and C, respectively.

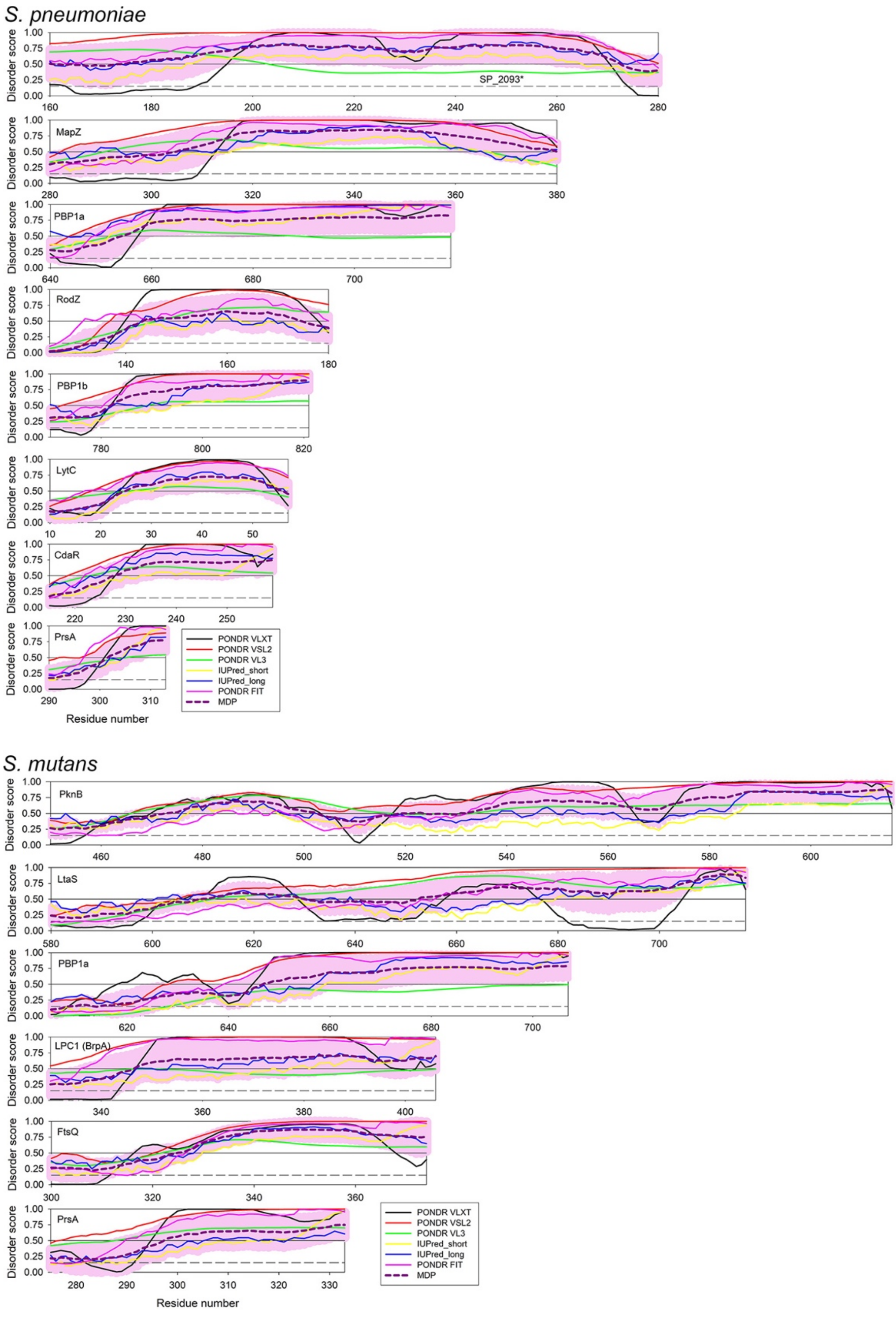


**Supplementary Fig. 10. Evaluation of the intrinsic disorder status of the Ser/Thr-rich regions in *S. pneumoniae* and *S. mutans* proteins analyzed in this study.**

Plots represent the per-residue disorder profiles generated for the indicated proteins by the Rapid Intrinsic Disorder Analysis Online (RIDAO) platform [26]. RIDAO integrates six established disorder predictors, such as PONDR® VLS2, PONDR® VL3, PONDR® VLXT, PONDR® FIT, IUPred-Long, and IUPred-Short into a single, unified platform. The mean disorder profiles (MDP) for these proteins were calculated for each protein by averaging the disorder profiles of individual predictors. The light pink shade represents the MDP error distribution. The thin black line at the disorder score of 0.5 is the threshold between order and disorder, where residues/regions above 0.5 are disordered, and residues/regions below 0.5 are ordered. The dashed line at the disorder score of 0.15 is the threshold between order and flexibility, where residues/regions above 0.15 are flexible, and residues/regions below 0.15 are mostly ordered.

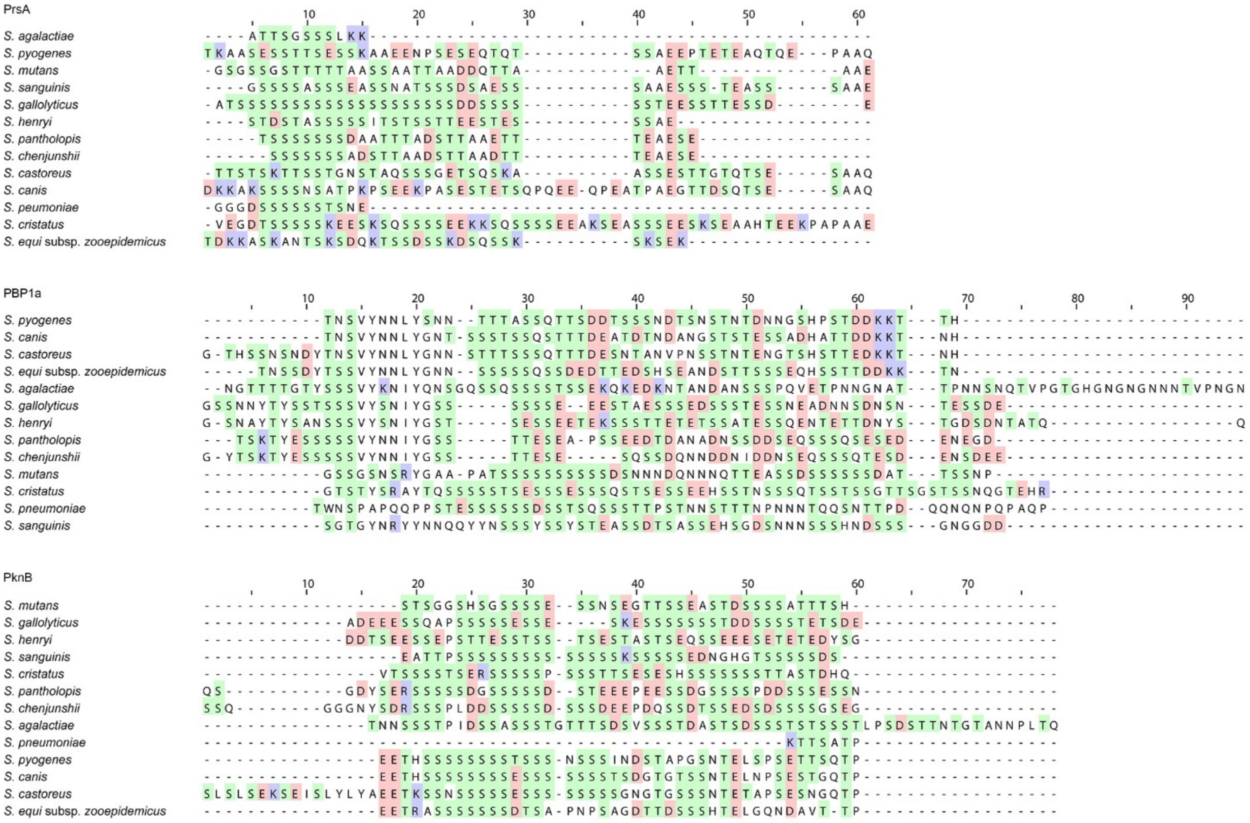


**Supplementary Fig. 11. Sequence alignments of IDRs identified in PrsA, PBP1a, and PknB homologs.**

Ser and Thr residues are highlighted in green, Arg and Lys residues in blue, and Asp and Glu residues in red.

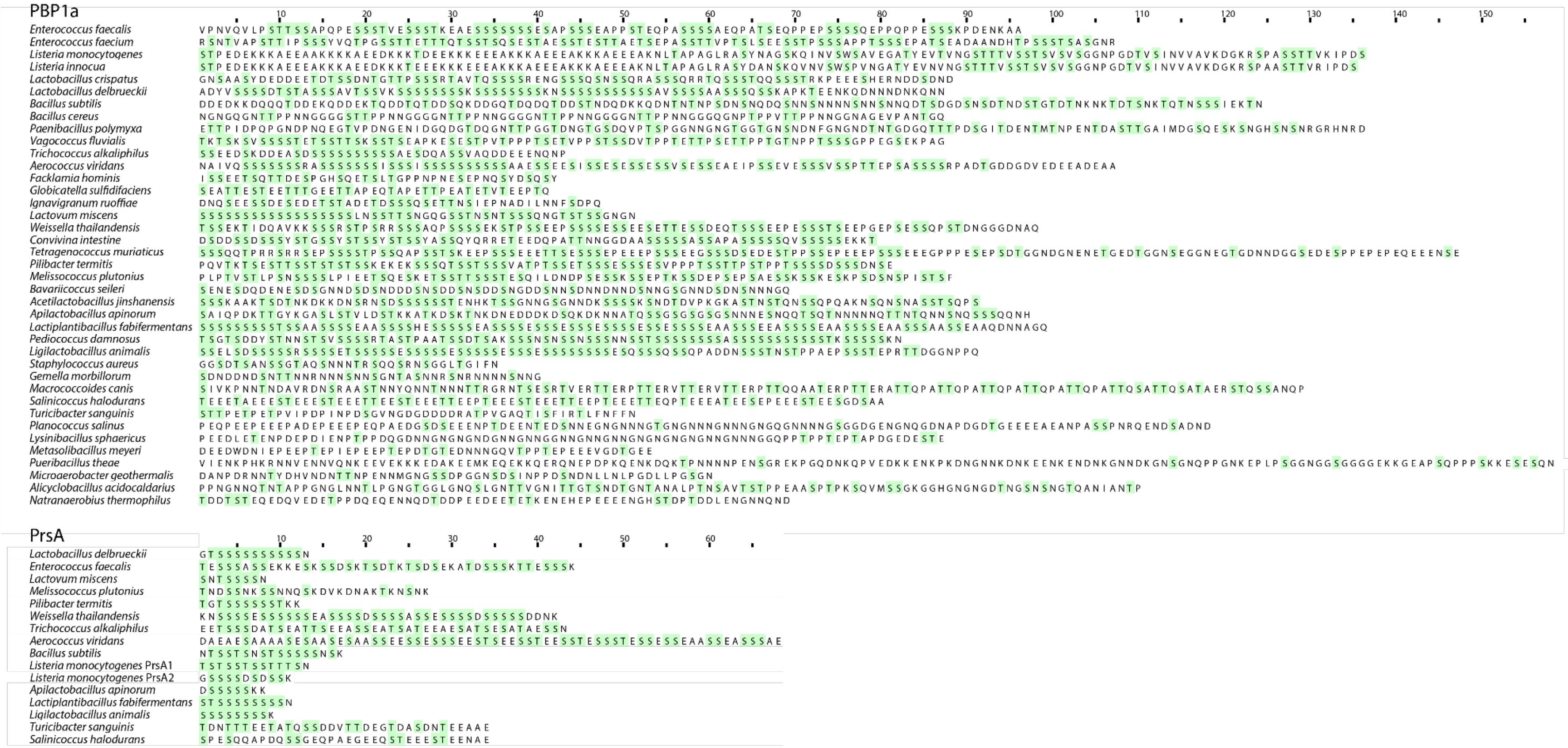


**Supplementary Fig. 12. Sequences of IDRs of PBP1a and PrsA homologs from Firmicutes.**

Ser and Thr residues are highlighted in green.

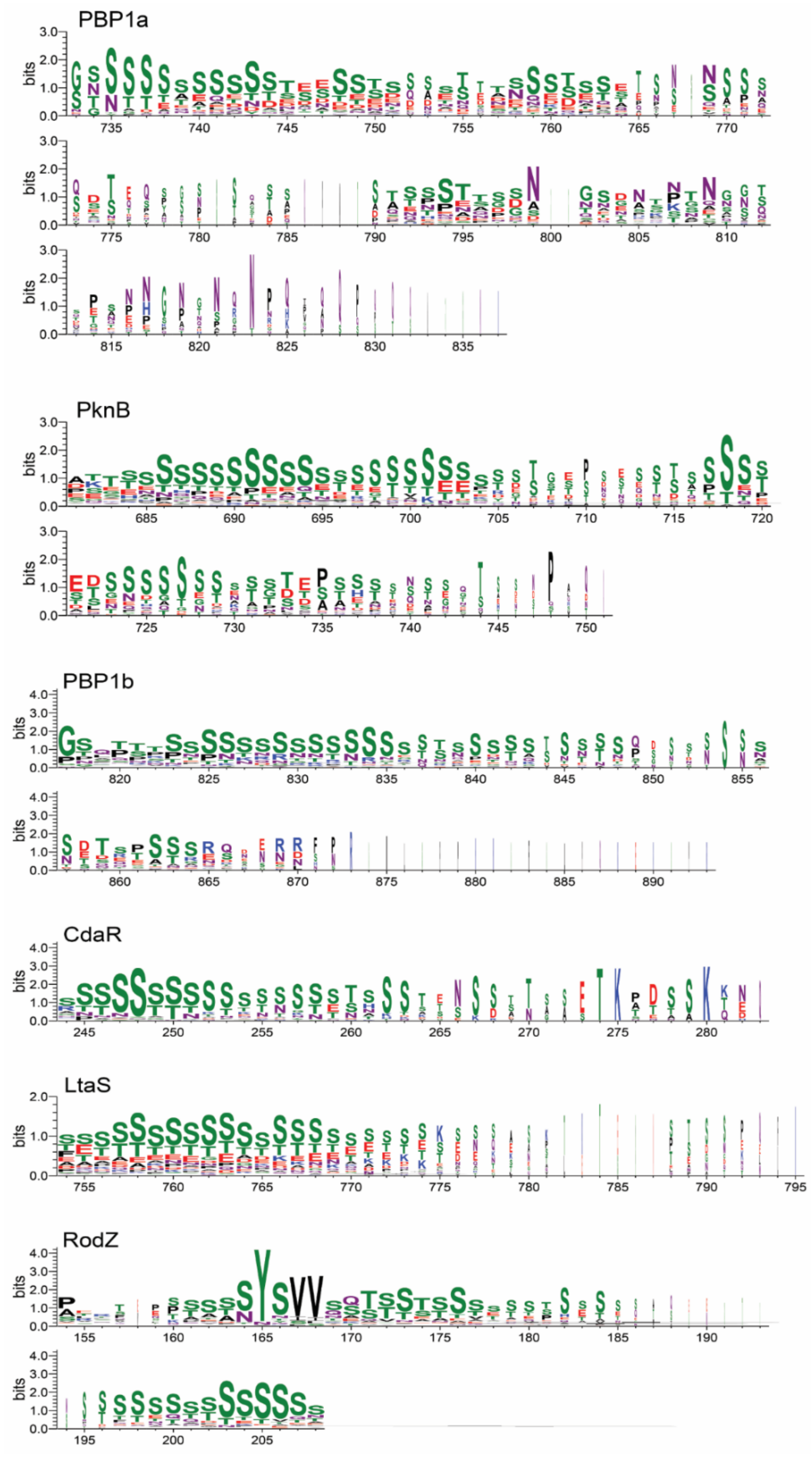


**Supplementary Fig. 13. The sequence logos of the IDRs of PBP1a, PknB, PBP1b, CdaR, LtaS and RodZ homologs from streptococcal species.**

The sequence logos are based on multiple sequence alignments of PrsA, PBP1a, PknB, PBP1b, CdaR, LtaS and RodZ homologs from streptococcal species and were generated by WebLogo [27].

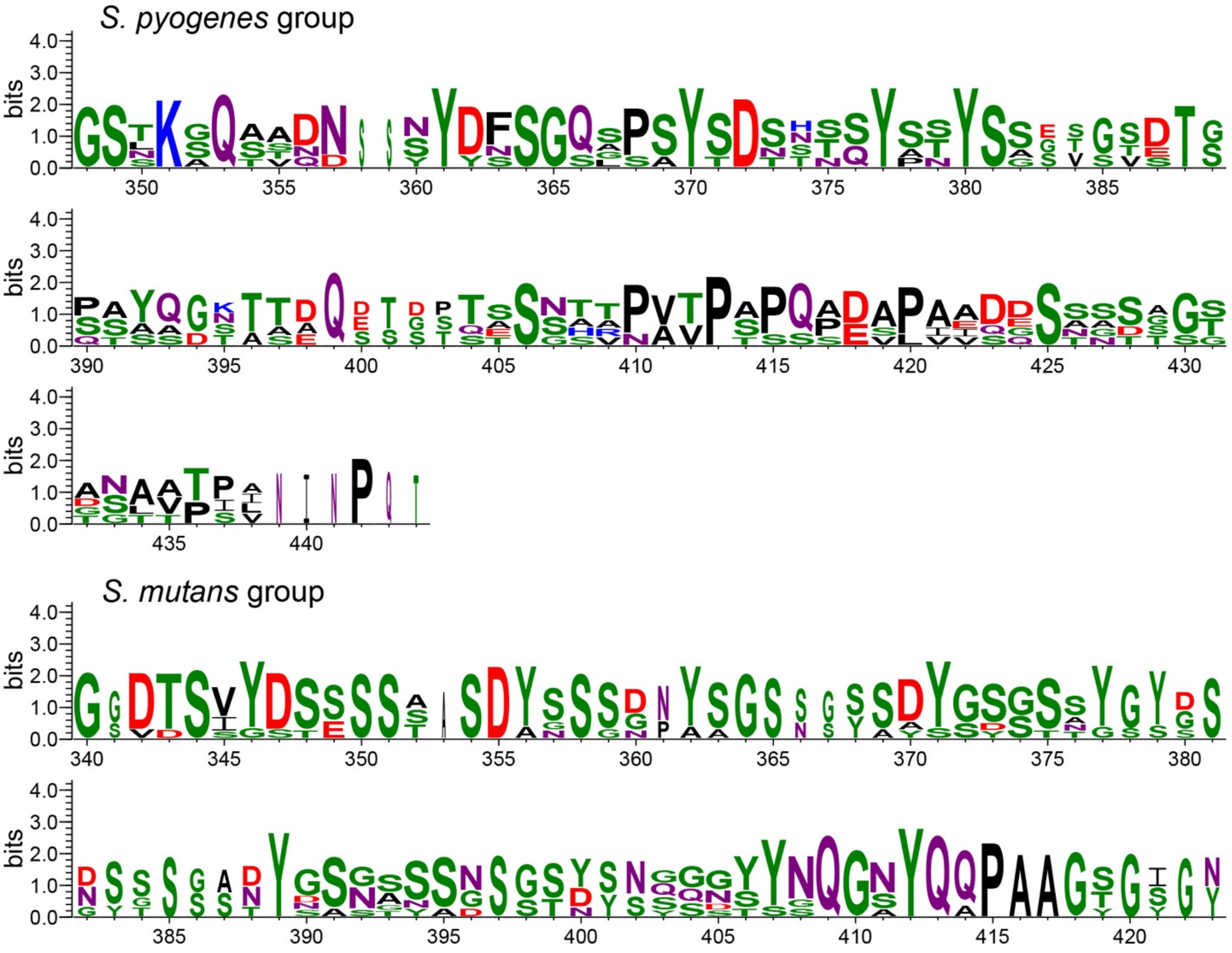


**Supplementary Fig. 14. The divergent IDRs of LytR homologs from streptococcal species.** The sequences of IDRs linked to LytR homologs from streptococcal species were separated into two groups based on the sequence similarity. The IDRs identified in the *S. mutans* LytR (known as BrpA) group contain an imperfect repeat motif (G/S)SSS(D/S)Y.

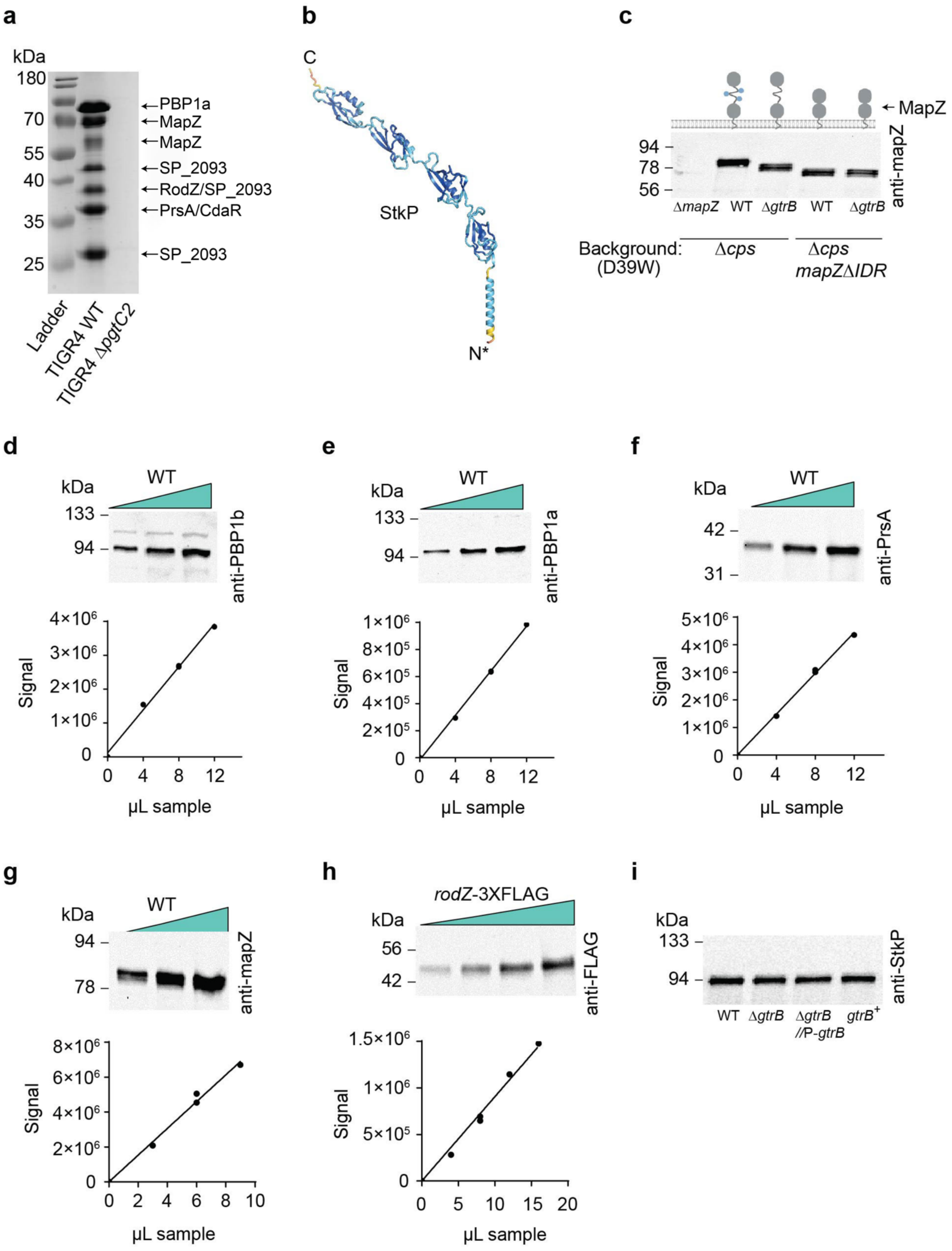

a
kDa
180
70
55
40
35
25
PBP1a
MapZ
MapZ
SP_2093
RodZ/SP_2093
PrsA/CdaR
SP_2093
Ladder
TIGR4 WT
TIGR4 ΔpgtC2
b
C
StkP
N*
c
MapZ
94
78
56
anti-mapZ
ΔmapZ
WT
ΔgtrB
WT
ΔgtrB
Background: (D39W)
Δcps
Δcps mapZΔIDR
d
WT
kDa
133
94
anti-PBP1b
4×10^6
3×10^6
2×10^6
1×10^6
0
Signal
0
4
8
12
μL sample
e
WT
kDa
133
94
anti-PBP1a
1×10^6
8×10^5
6×10^5
4×10^5
2×10^5
0
Signal
0
4
8
12
μL sample
f
WT
kDa
42
31
anti-PrsA
5×10^6
4×10^6
3×10^6
2×10^6
1×10^6
0
Signal
0
4
8
12
μL sample
g
WT
kDa
94
78
anti-mapZ
8×10^6
6×10^6
4×10^6
2×10^6
0
Signal
0
2
4
6
8
10
μL sample
h
rodZ-3XFLAG
kDa
56
42
anti-FLAG
1.5×10^6
1×10^6
5×10^5
0
Signal
0
5
10
15
20
μL sample
i
kDa
133
94
anti-StkP
WT
ΔgtrB
ΔgtrB //P-gtrB
gtrB+

**Supplementary Fig. 15. Purification, quantification, and validation of *S. pneumoniae* membrane glycoproteins.**

(**a**) A Coomassie-stained gel of *S. pneumoniae* TIGR4 membrane proteins purified by concanavalin A affinity chromatography. Proteins in excised bands were identified by LC-MS/MS analysis as described in Methods. The identified proteins are indicated. Predicted structural models of identified glycoproteins are shown in Supplementary Fig. 9. The experiments were performed independently two times in **a**. Representative image from one experiment is shown. (**b**) Predicted AlphaFold2 [28] structural model of *S. pneumoniae* StkP. The colors correspond to per-residue predicted local distance difference test (pLDDT) values. The amino- and carboxyl ends of proteins are labeled as N and C, respectively. The cytoplasmic domain of StkP is omitted for clarity (indicated by an asterisk). Just one monomer of the dimer for StkP is shown. (**c**) Immunoblot analysis of MapZ in the *S. pneumoniae* Δ*cps* WT (IU1824), Δ*gtrB*, *mapZ*ΔIDR and *mapZ*ΔIDRΔ*gtrB* strains. Schematic representation of MapZ is shown in the top panel and the cytoplasmic domain of this protein is omitted for clarity. The experiments were performed independently three times and a representative image from one experiment is shown. (**d**, **e**, **f**, **g** and **h**) Immunoblot of PBP1b (**d**), PBP1a (**e**), PrsA (**f**), MapZ (**g**) and RodZ (**h**) to show linear correlations between protein amounts and band signal intensity. Top panel, western blot images, and bottom panel, western standard curves generated by plotting different volumes (4, 8 or 12 μL) of WT lysates with antibody signals. The amounts of protein in the unknown samples shown in Fig. 3 **c**, **d, e, f,** and **g** were obtained from the linear regression interpolation values of the standard curves. The experiments were performed independently three times in **d**, **e, f, g,** and **h**. A representative image from one experiment is shown. (**i**) Immunoblot analysis of StkP in the *S. pneumoniae* D39W *cps*$^+$ WT (IU1781), and isogenic Δ*gtrB*, Δ*gtrB*//P*gtrB*, and *gtrB*$^+$ strains using *S. pneumoniae* anti-StkP antibodies. StkP is a serine-threonine kinase and is a homolog of PknB of *S. mutans* and *S. pyogenes*. The experiments were performed independently two times in **i**. A representative image from one experiment is shown. Proteins were separated on a 4-12% SurePAGE™ Bis-Tris gel (Genscript) in MES buffer in **a**, 7.5% Mini-PROTEAN® TGX™ precast protein gel (Biorad 4561025) in **d** and **e**, 10% precast protein gel (Biorad 4561035) in **f** and **i**, and 4–15% Mini-PROTEAN® TGX™ Precast Protein Gels (Biorad 4561085) in **c**, **g** and **h**. Source data for **a, c, d, e, f**, **g, h** and **i** are provided as a Source Data file.

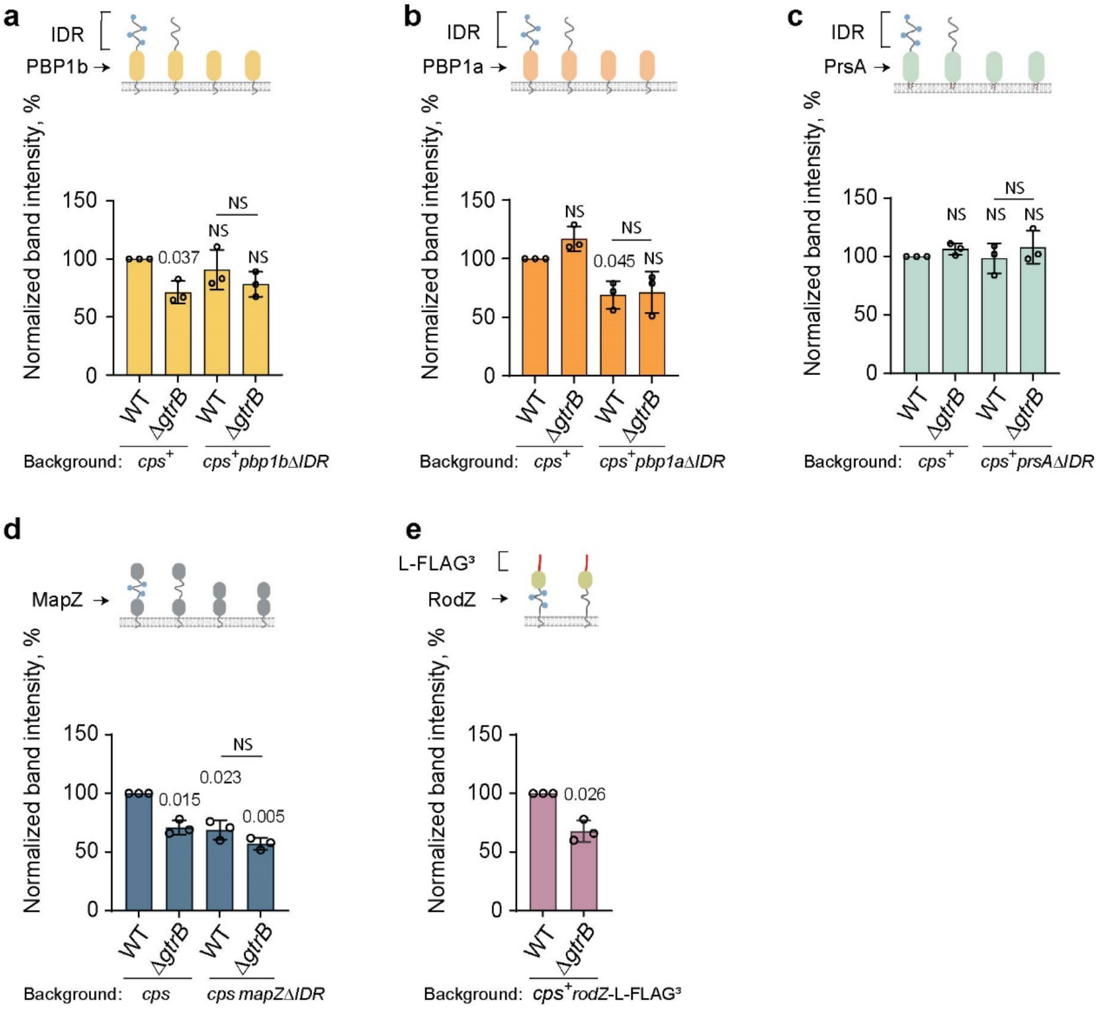


**Supplementary Fig. 16. Quantification of band intensities of *S. pneumoniae* PBP1b, PBP1a, PrsA, MapZ, and RodZ shown in representative immunoblots in Fig. 3c, d, e, g and Supplementary Fig. 15c.**

(**a**) Relative band intensities of PBP1b in the *S. pneumoniae* $cps^+$ WT (IU1781), and the isogenic strains, Δ*gtrB*, *pbp1b*ΔIDR, *pbp1b*ΔIDRΔ*gtrB*. (**b**) Relative band intensities of PBP1a in the *S. pneumoniae* $cps^+$ WT (IU1781), Δ*gtrB*, *pbp1a*ΔIDR and *pbp1a*ΔIDRΔ*gtrB* strains. (**c**) Immunoblot band intensities of PrsA in the *S. pneumoniae* $cps^+$ WT (IU1781), Δ*gtrB*, *prsA*ΔIDR and *prsA*ΔIDRΔ*gtrB* strains. (**d**) Immunoblot band intensities of MapZ in the *S. pneumoniae* Δ*cps* WT (IU1824), Δ*gtrB*, *mapZ*ΔIDR and *mapZ*ΔIDRΔ*gtrB* strains. (**e**) Relative band intensities of RodZ in the L-FLAG$^3$-tagged variants of the WT and Δ*gtrB* backgrounds. Schematic representations of PBP1b (**a**), PBP1a (**b**), PrsA (**c**), MapZ (**d**), and RodZ-L-FLAG$^3$ (**e**) are shown in the top panels. Each of the single blobs of PBP1b and PBP1a in the schematic represents two extracellular domains. Glc is shown as blue circles. Band intensities from each lane of the immunoblot were determined using Azure biosystem 600 software and normalized as described in Methods. Columns and error bars represent the mean and S.D., respectively (n = 3 biologically independent replicates). *P* values were determined by two-tailed one sample *t*-test to a reference value of 100. *P* values between ∆IDR strains in **a**, **b**, **c**, and **d** were determined by two sample *t*-test. Source data for **a, b, c, d,** and **e** are provided as a Source Data file.

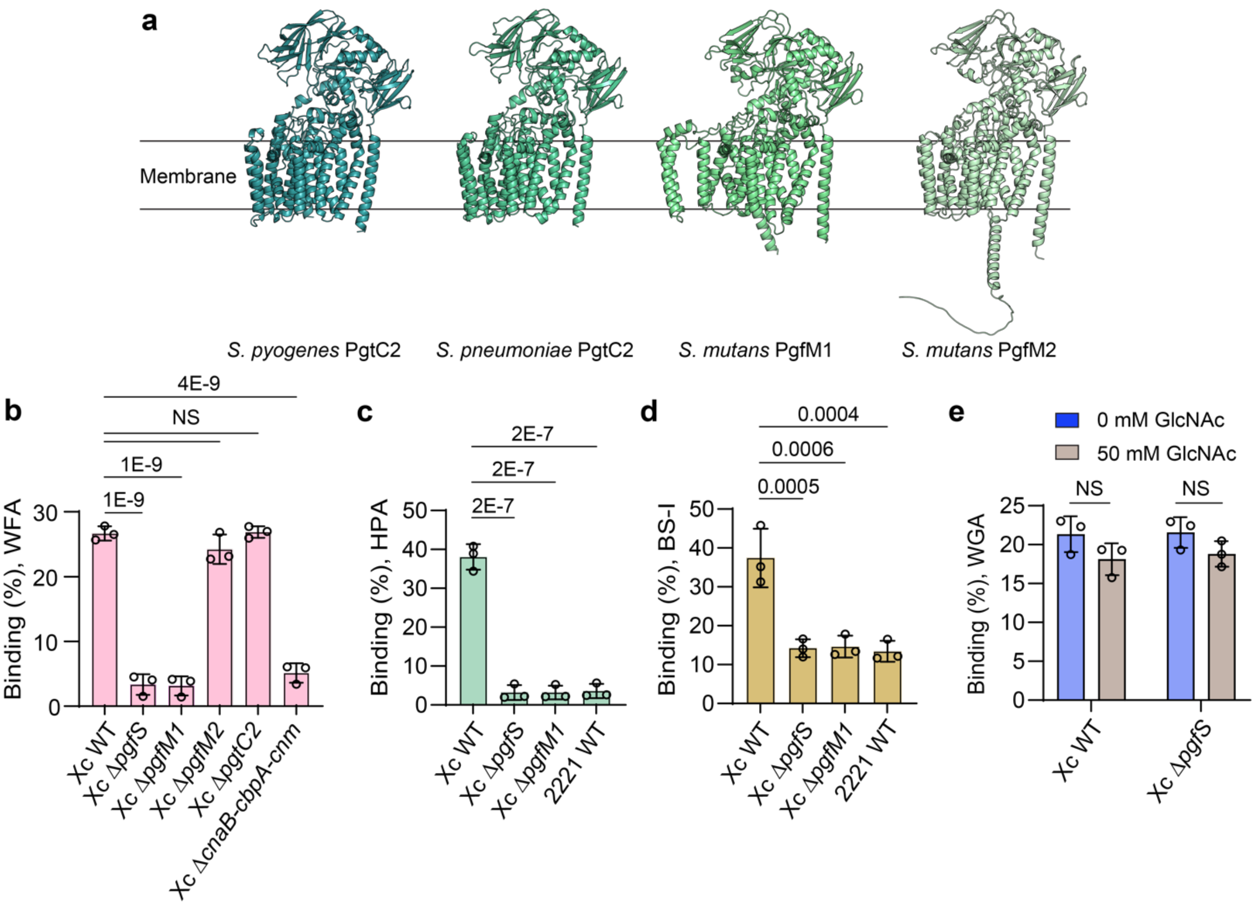


**Supplementary Fig. 17. The Pgf pathway is critical for *S. mutans* binding to the GalNAc-specific lectins.**

(**a**) AlphaFold2 structural models of *S. pyogenes* PgtC2, *S. pneumoniae* PgtC2, *S. mutans* PgfM1 and PgfM2 are shown in cartoon representation. The membrane topology was predicted using TOPCONS [29]. (**b**) Binding of *S. mutans* WT and the mutants to FITC-conjugated WFA. (**c**) Binding of bacteria to Alexa Fluor™ 488-conjugated HPA. (**d**) Binding of bacteria to FITC-conjugated BS-I. (**e**) Binding of *S. mutans* strains to Alexa Fluor™ 680-conjugated WGA. WFA and HPA selectively bind N-acetyl-α-D-galactosamine (GalNAc), whereas BS-I binds both α-D-galactose and GalNAc. WGA selectively binds N-acetylglucosamine (GlcNAc) and N-acetylneuraminic acid. WT is *S. mutans* Xc in **b**, **c**, **d**, and **e**. *S. pyogenes* MGAS 2221 (2221 WT) was used as a negative control in **c** and **d**. The following lectin concentrations were used: WFA, 50 µg $mL^{-1}$; HPA, 50 µg $mL^{-1}$; BS-I, 10 µg $mL^{-1}$; WGA, 50 µg $mL^{-1}$. Columns and error bars represent the mean and S.D., respectively (n = 3 biologically independent replicates). *P* values were calculated by one-way ANOVA with Dunnett's multiple comparisons test (**b**, **c**, and **d**) and by two-way ANOVA with Bonferroni's multiple comparisons test (**e**). NS indicates non-significant. Source data for **b, c, d** and **e** are provided as a Source Data file.

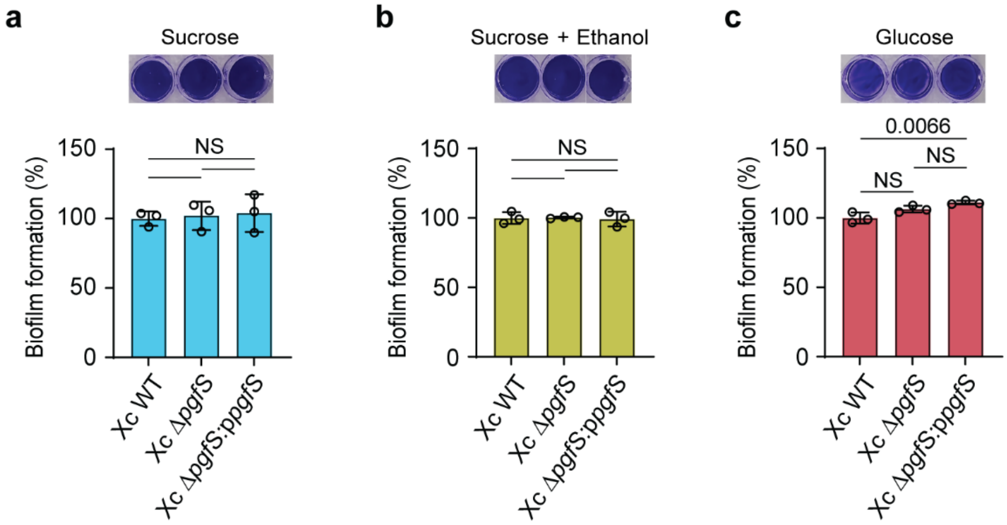


**Supplementary Fig. 18. Exopolysaccharide- and protein-based biofilm formation by *S. mutans* WT, XcΔ*pgfS*, and XcΔ*pgfS*:p*pgfS* strains.**

(**a** and **b**) Biofilm biomass of WT, XcΔ*pgfS*, and XcΔ*pgfS*:p*pgfS* strains grown in normal conditions (**a**), and with 3.5 % ethanol (**b**). Biofilms were grown 24 h in the UFTYE-sucrose medium. (**c**) Biofilm biomass of WT, XcΔ*pgfS*, and XcΔ*pgfS*:p*pgfS* strains grown 24 h in the UFTYE-glucose medium for 24 h. In **a**, **b** and **c**, Biofilms were stained with crystal violet followed by washes and resuspension in a de-staining solution. The absorbance of crystal violet in the de-staining solution was determined at 540 nm. Columns and error bars represent the mean and S.D., respectively (n = 3 biologically independent replicates). *P* values were calculated by one-way ANOVA with Tukey's multiple comparisons test. . Source data for **a, b** and **c** are provided as a Source Data file.

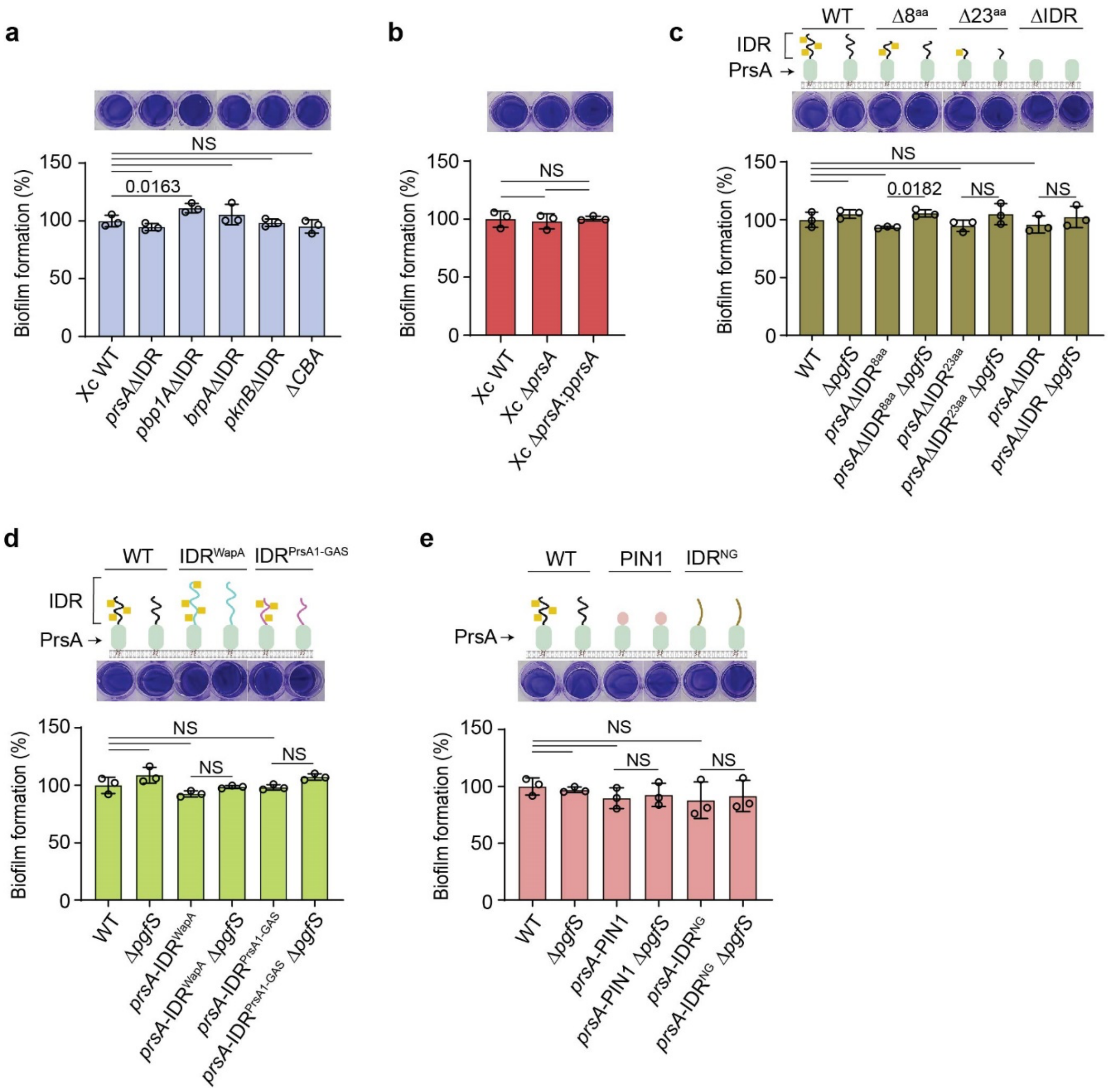


**Supplementary Fig. 19. Qunatification of *S. mutans* protein-based biofilm produced in normal conditions.**

(**a**) Biofilm formation of *S. mutans* WT and the isogenic mutants expressing the IDR-less variants of PrsA, PknB, BrpA, PBP1a (*prsA*ΔIDR, *pknB*ΔIDR, *brpA*ΔIDR and *pbp1a*ΔIDR) or deficient in four cell wall-associated adhesins WapA, Cnm, CnaA and CnaB (ΔCBA). The *pbp1a*ΔIDR mutant is the *pbp1a*ΔIDR-3×FLAG strain. (**b**) Biofilm formation of *S. mutans* WT, XcΔ*prsA,* and XcΔ*prsA*:p*prsA*. (**c**) Biofilm formation of *S. mutans* strains containing truncations in the C-terminal IDR of PrsA. (**d** and **e**) Biofilm formation of *S. mutans* strains expressing PrsA variants with the chimeric IDRs. In **c, d** and **e**, schematic representations of PrsA variants are shown (top panel). GalNAc is depicted as yellow squares. In **a**, **b**, **c**, **d**, and **e,** bacterial strains were grown in the

UFTYE-glucose medium for 24 h. Representative biofilms stained with crystal violet are shown in in the top panels. Quantification of biofilm formation is shown in the bottom panels. Columns and error bars represent the mean and S.D., respectively (n = 3 biologically independent replicates). *P* values were calculated by one-way ANOVA with Tukey's multiple comparisons test. Source data for **a, b, c, d** and **e** are provided as a Source Data file.

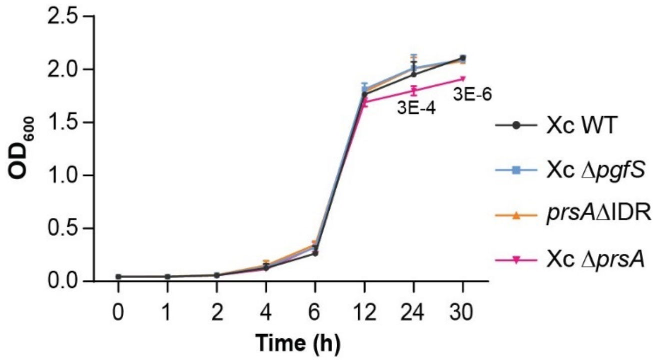


**Supplementary Fig. 20. Comparative growth analysis of the *S. mutans* WT, Xc∆*pgfS*, *prsA*∆IDR and Xc∆*prsA* strains.**

*S. mutans* strains were grown in UFTYE supplemented with 1% (wt/vol) glucose and 3.5% ethanol for 30 h at 37 °C. Then, the absorbance of cell suspensions was measured at 600 nm. The experiments were performed independently three times. Error bars represent the S.D. *P* values were calculated by a two-way ANOVA with Dunnett's multiple comparisons test. *P* values between Xc WT and Xc∆*prsA* are shown. Source data are provided as a Source Data file.

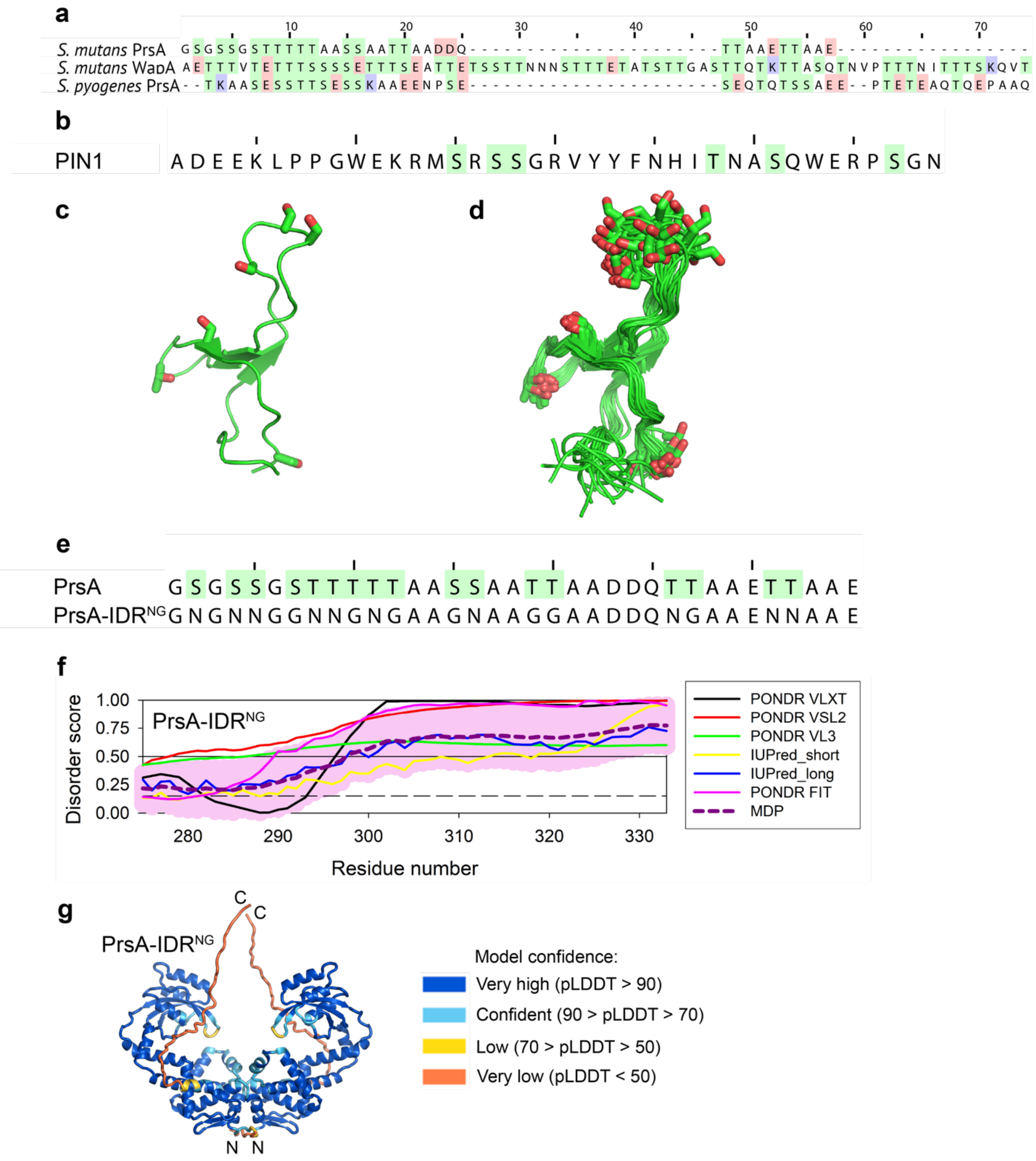


**Supplementary Fig. 21. Design of PrsA IDR fusion variants.**

(**a**) A sequence alignment of IDRs of *S. mutans* PrsA and WapA, and *S. pyogenes* PrsA1. Ser and Thr residues are highlighted in green, Arg and Lys in blue, and Asp and Glu in red. (**b**) The sequence of the PIN1 WW domain with serine and threonine residues highlighted in green. (**c**) The NMR structure of the PIN1 WW domain is shown in a cartoon representation (PDB ID 2KCF) [30]. The serine and threonine residues are shown as sticks. (**d**) A cartoon representation of 20

NMR conformers of the PIN1 WW domain. (**e**) The alignment of the wild-type PrsA IDR sequence with the NG substitution sequence where serine and threonine residues are randomly substituted with either asparagine (N) or glycine (G) residues. Analysis by DR-BERT indicated that the substituted sequence is predicted to be disordered, with per-residue DR-BERT scores >0.8. (**f**) Analysis of disorder propensity in PrsA-IDR$^{NG}$ variant. Plots represent the per-residue disorder profiles generated by the Rapid Intrinsic Disorder Analysis Online (RIDAO) platform [26]. (**g**) The AlphaFold2 model of the PrsA-IDR$^{NG}$ variant is shown in a cartoon representation. The colors correspond to per-residue predicted local distance difference test (pLDDT) values.

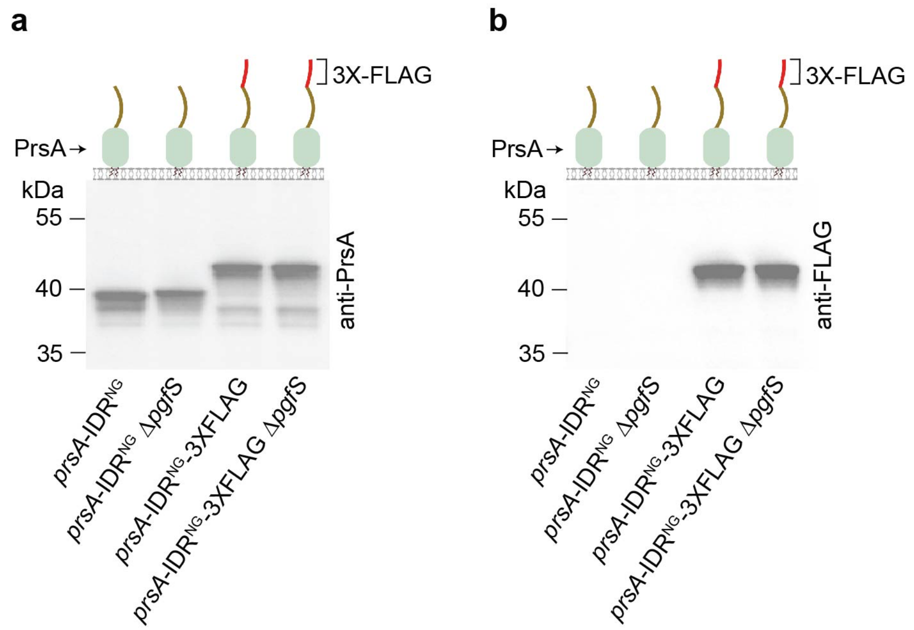


**Supplementary Fig. 22. Immunoblot analysis of *S. mutans* PrsA variants possessing the chimeric IDRs and 3×FLAG-tag in the WT and XcΔ*pgfS* strain backgrounds.**

(**a** and **b**) Immunoblot analysis of *S. mutans* PrsA variants probed with the anti-PrsA antibodies (**a**) and the anti-FLAG antibodies (**b**). Proteins were separated on a 4-12% SurePAGE™ Bis-Tris gel (Genscript) in MES buffer. The experiments were performed independently two times in **a** and **b**, and yielded the same results. A representative image from one experiment is shown. Source data for **a** and **b** are provided as a Source Data file.

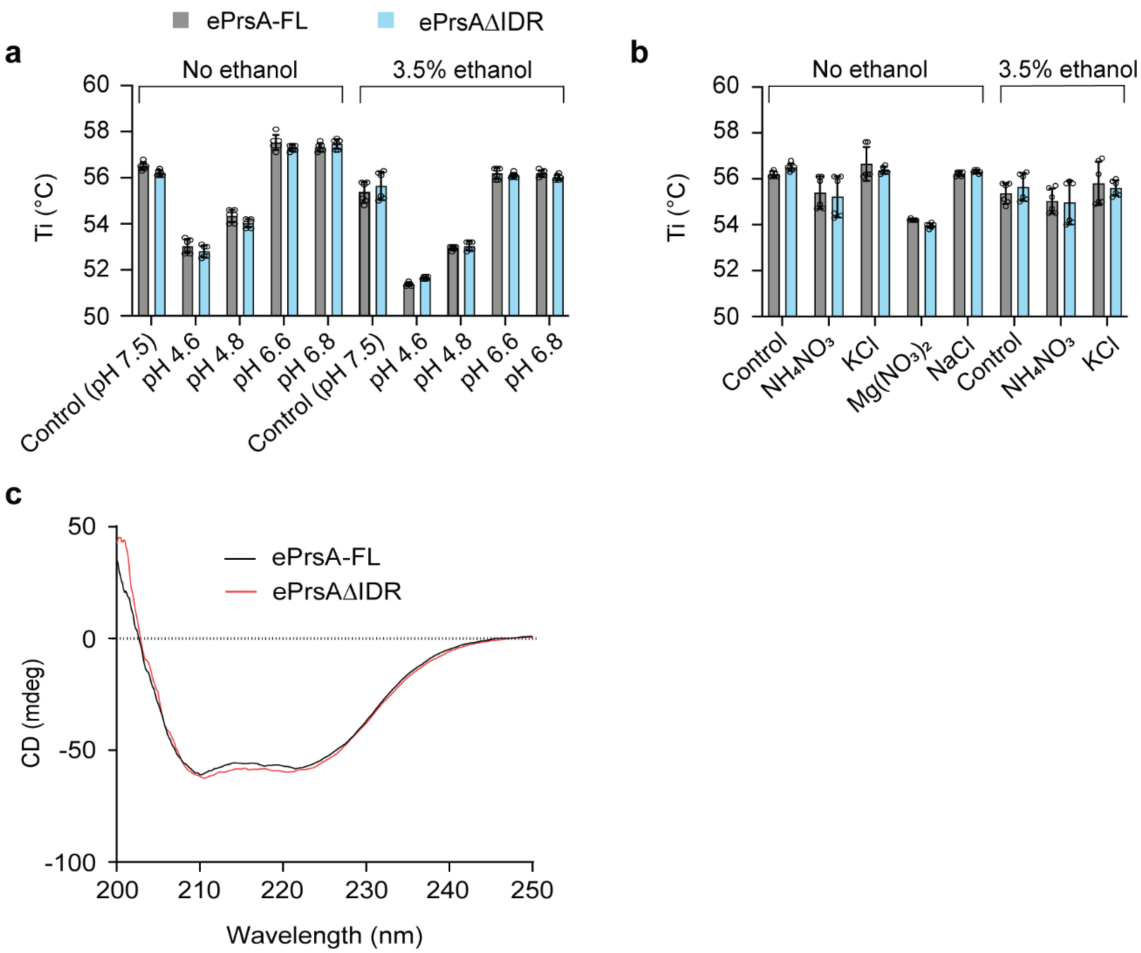


**Supplementary Fig. 23. Thermal denaturation and CD analysis of ePrsA-FL and ePrsAΔIDR.** (**a** and **b**) Comparison of the stability of ePrsA-FL and ePrsAΔIDR recombinant proteins in different pH- (**a**) and salts (**b**) buffers at normal and ethanol-stressed conditions. The control buffer contains 20 mM Tris HCl pH 7.5, 300 mM NaCl, and all tested buffers were adjusted in the control buffer background. 100 mM of tested salt concentration was used in **b**. Protein unfolding was recorded using intrinsic fluorescence at 330 nm and 350 nm, and the inflection temperatures (Ti) were determined by the fluorescence ratio (350/330 nm). Data presented are mean±S.D of six replicates from two independent protein purification samples. *P* values were determined by two-way ANOVA with Bonferroni's multiple comparisons test. All data were non-significant ($P>0.05$); therefore, they are not shown. (**c**) Superimposition of CD spectra of ePrsA-FL and ePrsAΔIDR recombinant proteins. Proteins were dialyzed against 20 mM sodium phosphate pH 7.4, and 150 mM sodium fluoride buffer. CD measurements were acquired on a JASCO J-810 spectropolarimeter over 200–250 nm wavelength range at 25 °C. Source data for **a, b** and **c** are provided as a Source Data file.

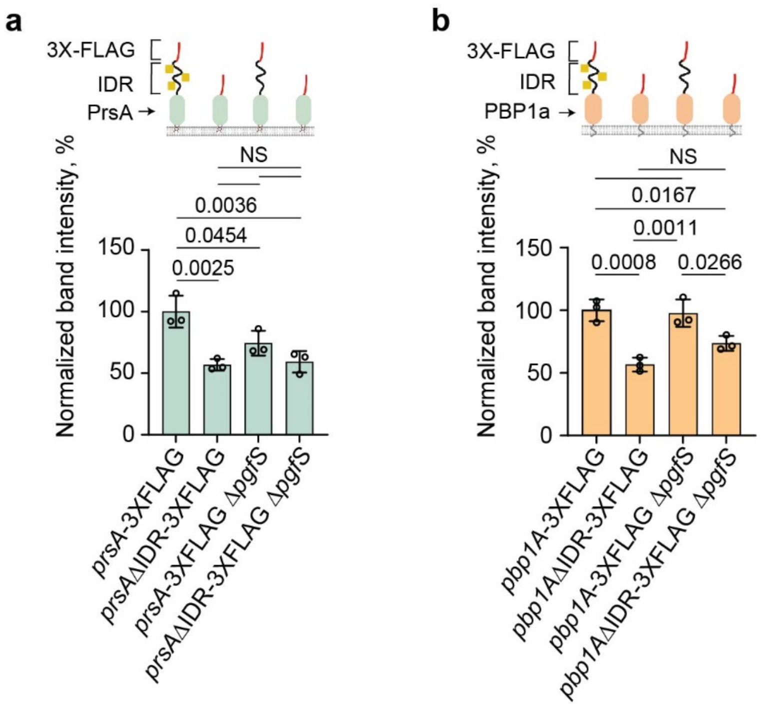


**Supplementary Fig. 24. Quantification of band intensities of *S. mutans* PrsA and PBP1a shown in representative immunoblots in Fig. 4h and i.**

(**a** and **b**) Relative band intensities of FLAG-tag fused variants of PrsA (**a**) and PBP1a (**b**) in the WT and Δ*pgfS* backgrounds. Schematic representations of PrsA and PBP1a variants are shown in the top panels. Each of the single blobs of PBP1a in **b** represents two extracellular domains. GalNAc is depicted as yellow squares. Band intensities from each lane of the immunoblot were calculated using ImageJ, and the data were normalized. Columns and error bars represent the mean and S.D., respectively (n = 3 biologically independent replicates). *P* values in **a** and **b** were determined by one-way ANOVA with Tukey's multiple comparisons test. Source data for **a** and **b** are provided as a Source Data file.

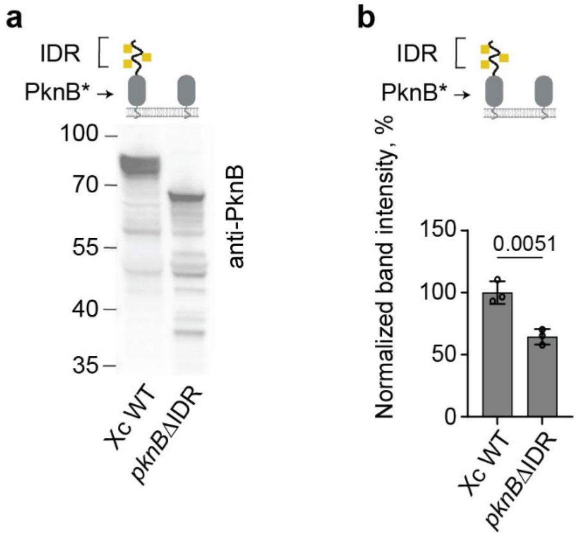


**Supplementary Fig. 25. Immunoblot analysis and quantification of band intensities of *S. mutans* PknB variants assessed by anti-PknB antibodies.**

(**a**) Immunoblot analysis of PknB variants probed by anti-PknB antibodies in *S. mutans*. The experiments were performed independently three times, and they yielded the same results. A representative image from one experiment is shown. Proteins were separated on a 4-12% SurePAGE™ Bis-Tris gel (Genscript) in MES buffer. (**b**) Immunoblot band intensities of PknB variants in *S. mutans*. Band intensities from each lane of the immunoblot were calculated using ImageJ, and the data were normalized. Columns and error bars represent the mean and S.D., respectively (n = 3 biologically independent replicates). *P* values were determined by two-tailed Student's *t*-test. In **a** and **b**, Schematic of PknB variants is shown in top panel. * The cytoplasmic domain of PknB in schematic is not shown. Each of the single blobs of PknB represents three extracellular domains. GalNAc is depicted as yellow squares. Source data for **a** and **b** are provided as a Source Data file.

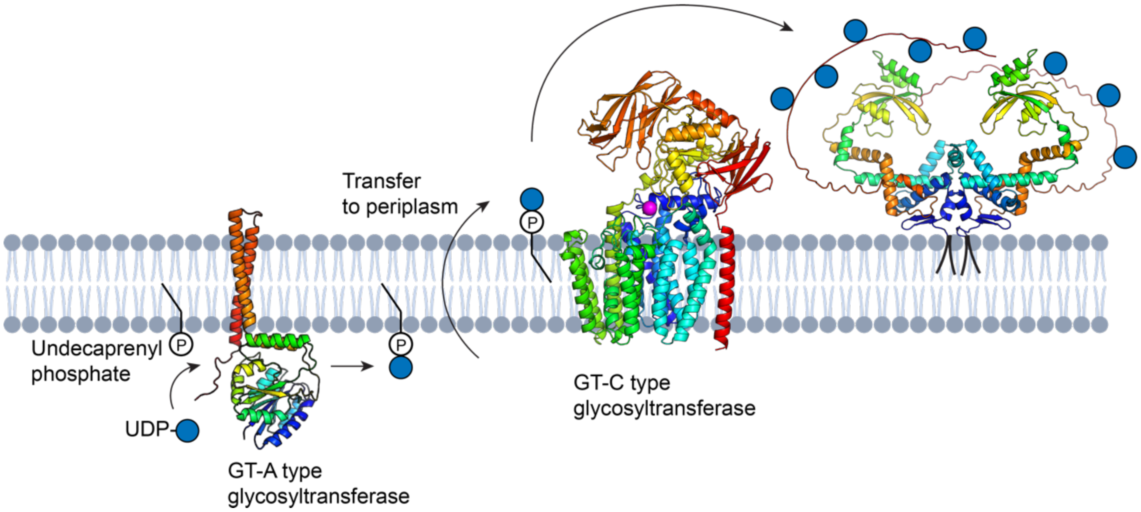


**Supplementary Fig. 26. *O*-glycosylation of extracytoplasmic proteins in streptococci.**
GT-A type glycosyltransferase (GtrB in *S. pyogenes* and *S. pneumoniae* and *pgfS* in *S. mutans*) transfers a sugar moiety (Glc in *S. pyogenes* and *S. pneumoniae*) from activated nucleotide-sugar (UDP-Glc in *S. pyogenes* and *S. pneumoniae*) to a lipid carrier, undecaprenyl phosphate. Undecaprenyl phosphate-sugar is transferred into the extracellular space by an unknown transporter, where it serves as a sugar donor (Glc in *S. pyogenes* and *S. pneumoniae* and GalNAc in *S. mutans*) for GT-C type glycosyltransferase (PgtC2 in *S. pyogenes* and *S. pneumoniae* and PgfM1 in *S. mutans*) to glycosylate the Ser/Thr-rich IDR of PrsA. AlphaFold2 models of *S. pyogenes* GtrB, PgtC2 and PrsA1 are shown as cartoons colored in rainbow from N-terminus in blue to C-terminus in red. Glc residues are depicted as blue circles. A model of PrsA1 dimer shows the diacylated lipoprotein embedded in the cell membrane. The magenta sphere highlights the key active-site Asp residue of PgtC2.

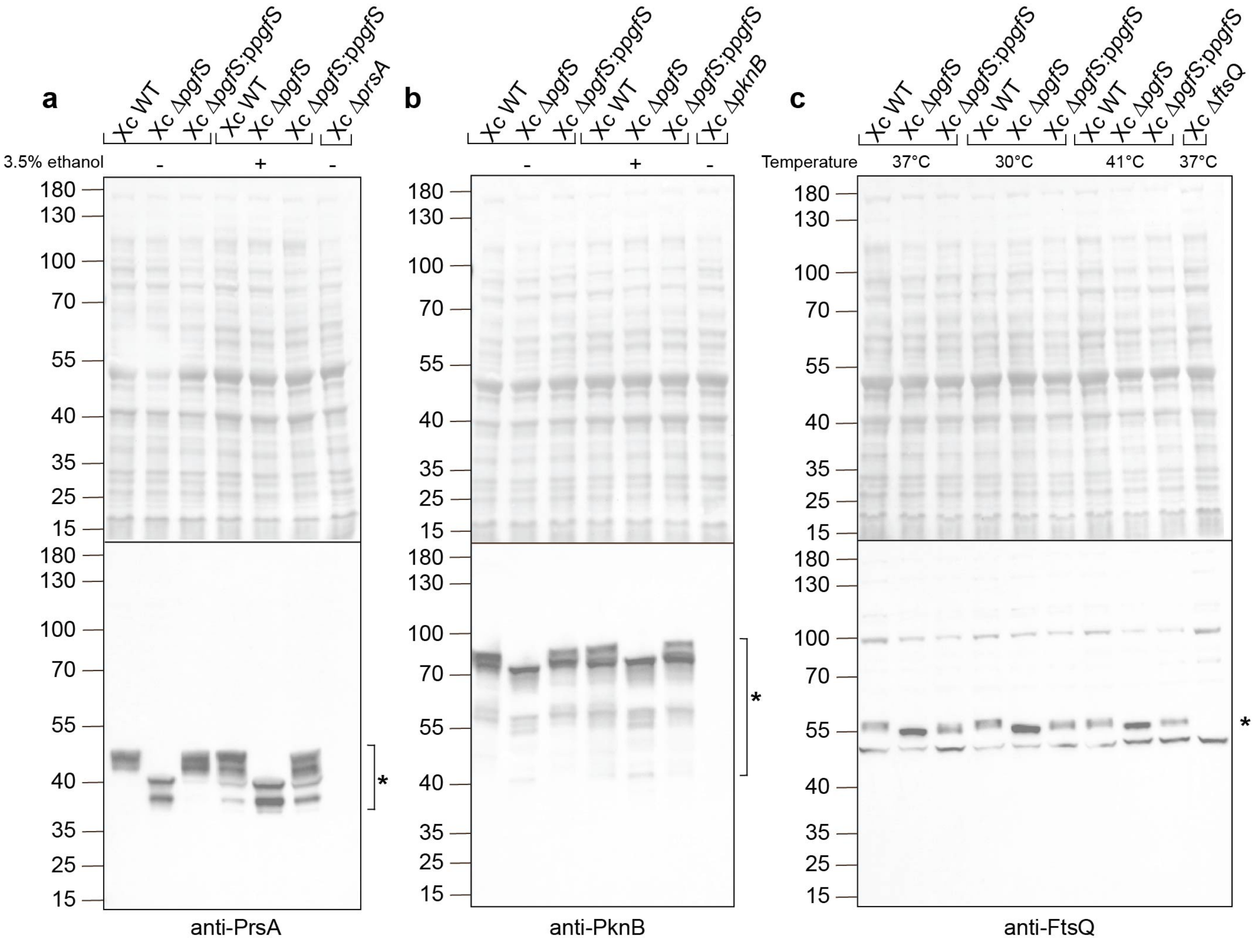


**Supplementary Fig. 27. Validation of antisera against *S. mutans* PrsA (a), PknB (b), and FtsQ (c).**

Antisera against purified recombinant PrsAΔIDR, FtsQΔIDR, and PknBΔIDR$^{Smu}$ were produced in rabbits by Cocalico Biologicals, Denver. In **a** and **b**, bacteria were grown in THY ± 3.5% ethanol medium at 37 °C, whereas in **c** and **d**, cells were grown in THY at different temperatures (37 °C, 30 °C and 40 °C) to $OD_{600}$ ~0.80. 10-15 μg of total protein was loaded in each lane. Proteins were separated on a 4-12% SurePAGE™ Bis-Tris gel (Genscript) in MES buffer and transferred onto a nitrocellulose membrane. Protein transfer was verified by Ponceau staining. The membrane was incubated with antisera dilutions: anti-PrsA, 1:1,000; anti-PknB$^{Smu}$, 1:500; anti-FtsQ, 1:500, and secondary antibodies goat anti-rabbit IgG conjugated with HRP 1:10,000 (ThermoFisher Scientific). The top panels in **a**, **b**, and **c** are images of Ponceau-stained blots, whereas the bottom panels are immunoblotting images. Protein MW standards are shown in kDa. Targeted protein bands are shown as a star symbol (*). Source data for **a, b** and **c** are provided as a Source Data file.
